# Discovery of evolutionarily extended *cis*-regulatory overlapping genes expanding the protein universe from animals to humans

**DOI:** 10.1101/2025.07.27.667088

**Authors:** Yuhta Nomura, Naoshi Dohmae

## Abstract

A recent proteogenomics-driven approach has uncovered actively translated alternative open reading frames (altORFs) that complement reference protein-coding sequences (refCDSs). Here, we further explored proteogenomic data to identify hidden proteomes, specifically short ORF-encoded polypeptides (SEPs) and longer altORF-encoded proteins (LEPs) previously overlooked in human genomes. We discovered a novel class of proteomes originating from SEP/LEP-coding upstream overlapping altORFs (oORFs) associated with refCDSs, which *cis*-regulate refCDS translation and have undergone evolutionary selection favoring C-terminal extension for the enhanced *cis*-regulation. These translatable oORFs occur in signal effectors of the Hippo-YAP/TAZ, p53, Wnt, and TGF-β crosstalk pathways and frequently arise from intragenic frameshift polymorphisms closely linked to human diseases. The intragenic frameshift mutations divide refCDSs and capture the N-terminal regions to form the new oORFs. Consequently, we termed these entities <u>ups</u>tream region-<u>u</u>surping repurposed <u>p</u>roteins (USURPs). These findings offer new insights into birth and evolution of proteomes, broadening our understanding of the protein universe influenced by genome dynamics.

**Teaser:** Unveiling a novel category of proteomes from animals to humans and elucidating its birth, evolution, function, and relation to human diseases

## Introduction

The central dogma of molecular biology, which includes the fundamental processes of transcription and translation, describes the unidirectional flow of genetic information for protein expression in all living cells. Traditionally, it has been assumed that eukaryotic proteins are produced from monocistronic transcripts, each containing a single protein-coding open reading frame (ORF), unlike their bacterial counterparts (*1*). However, recent advances in translatomics, specifically ribosome profiling (Ribo-Seq), and a proteogenomics strategy integrating mass spectrometry (MS)-based proteomics with RNA sequencing (RNA-Seq), have unveiled the presence of actively translated alternative ORFs (altORFs) on eukaryotic transcripts (*2–9*). These altORFs, distinct from primary ORFs (*i.e.*, reference protein-coding sequences; refCDSs), suggest more densely packed genetic information in genomes than previously anticipated. This multiomics approach is uncovering previously unrecognized biological entities, termed dark proteomes and peptidomes, encoded within eukaryotic genomes (*7*).

The proteogenomic method, bolstered by MS-based proteomics, offers a unique advantage in the discovery of translatable altORFs over traditional genomic or transcriptomic techniques that do not incorporate proteomic data. This is because MS-based proteomics can definitively identify peptides and proteins through direct detection. In contrast, traditional methods often exclude short ORFs from further study due to stringent criteria such as a cutoff of <150 amino acids, leading to the misclassification of many RNAs as non-coding (*2*). Additionally, only refCDSs, typically the longest ORFs on their transcripts, have been acknowledged and annotated as genes. This has resulted in the exclusion of almost all altORFs from consideration as gene candidates.

These sampling biases may contribute to the misconception that unexplored altORFs are merely short, insignificant ORFs, only acknowledged when their translation products are identified as short ORF-encoded polypeptides (SEPs) (*2*). The arbitrary threshold of <150 amino acids for SEPs potentially introduces further bias in uncovering hidden alternative genes (alt-genes). However, exceptions likely exist in the amino acid lengths of altORFs, which could contribute to an expanded protein universe and necessitate more precise genome annotation and a deeper understanding of genome evolution. In this study, we propose that altORFs of ≥150 amino acids be categorized into longer altORF-encoded proteins (LEPs), which may often overlap out-of-frame with refCDSs due to their extended lengths. We investigated both SEPs and LEPs to establish an unbiased approach for alt-gene discovery.

We previously identified a SEP-coding upstream overlapping altORF (oORF), *oSCRIB*, which is out-of-frame with a refCDS, the Scribble (SCRIB) protein, a key player in the Hippo-YAP/TAZ signaling pathway in humans (*10*). In this study, we expanded our proteogenomic data mining to include both SEPs and LEPs using publicly available, high-quality datasets of MS-based deep proteomics and RNA-Seq in humans. We discovered a new category of dark proteomes, consisting of SEPs and LEPs, encoded as *cis*-regulatory oORFs of biologically significant refCDSs. Furthermore, we investigated the molecular mechanism underlying the *de novo* birth and evolution of SEP/LEP-coding oORFs. Our analyses showed that these translatable oORFs have evolved to become longer at the C-terminal and are frequently generated in humans through frameshift polymorphisms. These findings suggest a further expansion of the protein universe influenced by genome dynamics.

## Results

### Transcriptomic and proteomic data mining for new SEP/LEP-coding altORFs in humans

Human-derived cell lines, such as neuroblastoma SH-SY5Y and lung adenocarcinoma A549, serve as *in vitro* models for human organs and tissues. SH-SY5Y cells resemble immature sympathetic neurons and express neuroproteins in culture (*11*), while A549 cells are a more differentiated cell line representing type II alveolar epithelial cells (*12*). Consequently, these cell lines are frequently used to study the characteristics of brain and lung cells, respectively.

To discover new translatable altORFs encoding SEPs or LEPs in these distinct cell lines (Fig. 1A), we assembled publicly available RNA-Seq data from these cell lines *de novo* and searched the resulting contigs for putative short ORFs of <150 amino acids and longer ORFs of ≥150 amino acids in humans. Next, we compared the translated nucleotide sequences (*i.e.*, amino acid sequences) of the identified short/longer ORFs with a set of human reference protein sequences (RefSeq Proteins (*13*); assembly accession no. GCF_000001405.39) available from the National Center for Biotechnology Information (NCBI). This comparison aimed to isolate short/longer ORFs not currently recognized as genes in the human RefSeq Proteins database. Consequently, the final custom sequence databases for each cell line included 5,279,489 and 1,731,286 SEPs, and 16,157 and 13,570 LEPs, for SH-SY5Y and A549, respectively.

**Fig. 1.**
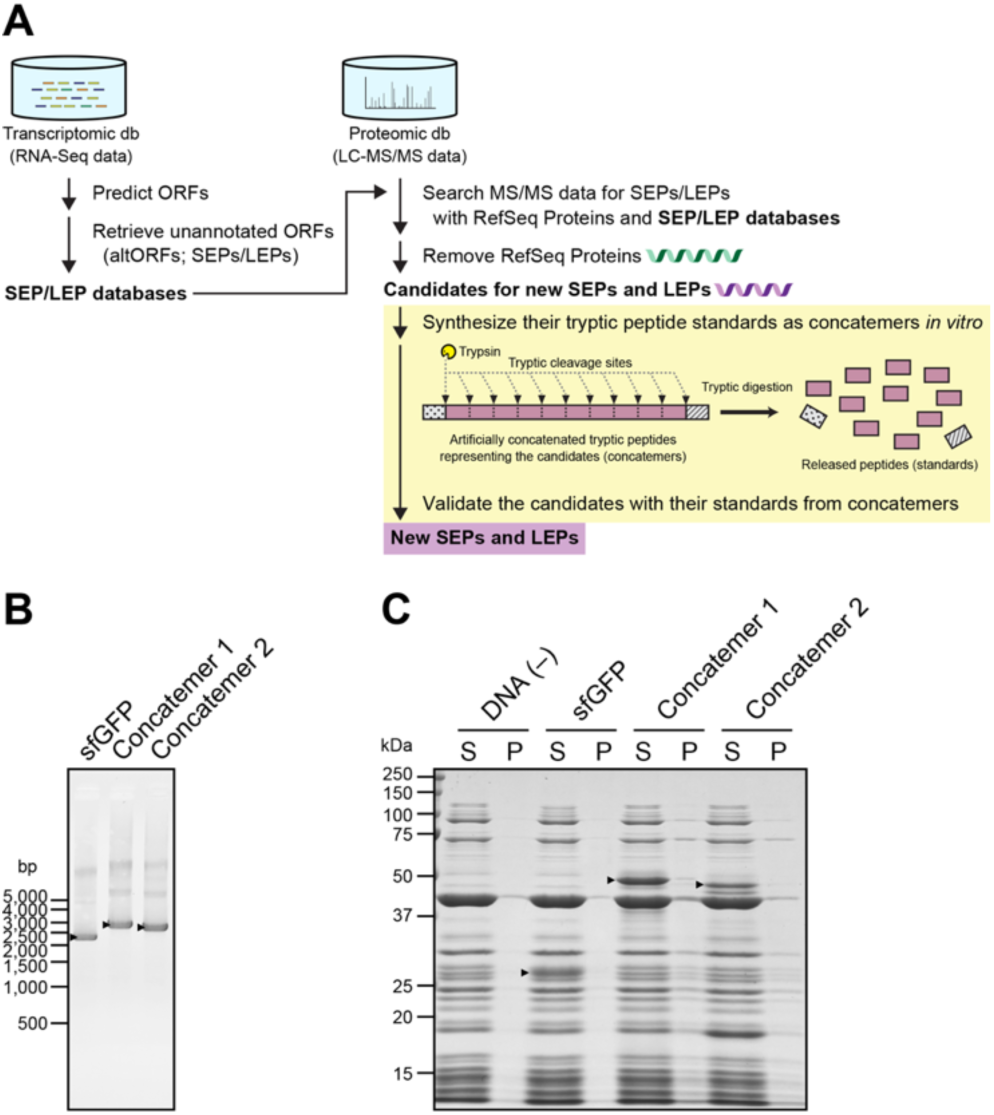
Proteogenomics-driven alt-gene discovery in humans. (**A**) The alt-gene discovery workflow implemented in this study, consisting of proteogenomic data mining for new candidate SEPs and LEPs and validation of their expression using artificially concatenated peptides (concatemers) to release multiple peptide standards. (**B**) Agarose gel electrophoresis of DNA fragments (*sfGFP* and concatemers) prepared for *in vitro* transcription and translation. DNA fragments were visualized under ultraviolet light on a gel stained with GelRed; the positions of molecular standards are indicated. Arrowheads indicate the DNA fragment bands. (**C**) SDS-PAGE analysis of *in vitro*-synthesized proteins (sfGFP and concatemers). Coupled *in vitro* reactions of transcription and translation with and without the DNA fragment encoding sfGFP protein were used as positive and negative control reactions, respectively. The gel was stained with Coomassie Brilliant Blue G-250; the positions of molecular standards are indicated. Arrowheads indicate the bands of proteins synthesized in vitro, whereas other bands were derived from enzymes used for transcription and translation and partial-length proteins. S, soluble fractions; P, insoluble fractions.

Next, we conducted proteogenomic analyses using high-quality, publicly available proteomic data (ProteomeXchange dataset identifier PXD004452 (*14*)) from SH-SY5Y and A549 cell lines (Fig. 1A). MS/MS peak lists, generated from proteomic mass spectra using Thermo Proteome Discoverer (Thermo Fisher Scientific), were searched against two distinct sequence databases: the human RefSeq Proteins combined with the cell-type-matched custom sequence database of SEPs or LEPs. This search, conducted using the Mascot engine (*15*), involved peptide-spectrum matching (PSM) followed by target-decoy based false discovery rate (FDR) filtering. This approach predicted the presence of 6 SEPs and 6 LEPs in either SH-SY5Y or A549 proteomes with high confidence, indicated by a low FDR ≤=1% at the peptide level and ≤=0.1% at the protein level (Table 1 and table S1).

**Table 1.**
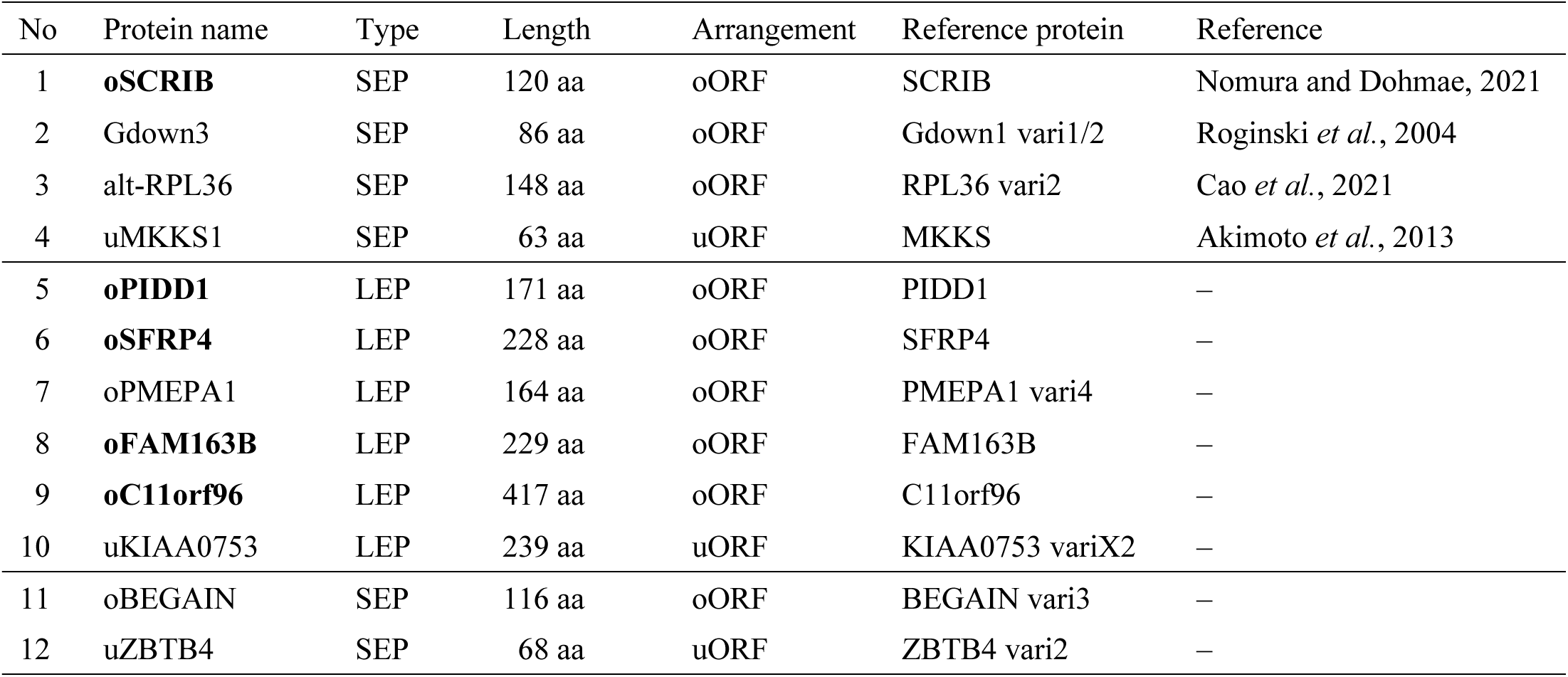
List of SEPs and LEPs in humans.

Our results include previously reported SEPs that are encoded as oORFs of the *SCRIB*, *Gdown1* (*POLR2M*) transcript variant (vari)1/2, and *RPL36* vari2, along with an upstream ORF (uORF) of *MKKS*. These SEPs, not included in the human RefSeq Proteins database, have been identified as oSCRIB, Gdown3, alt-RPL36, and uMKKS1, respectively (*10, 16–18*) (Table 1). In addition to these SEPs, our results also reveal the presence of new LEPs that are encoded as oORFs of *PIDD1*, *SFRP4*, *PMEPA1* vari4, *FAM163B*, and *C11orf96* (*AG2*), and a uORF of *KIAA0753* variX2. We also detected new SEPs encoded as an oORF of *BEGAIN* vari3 and a uORF of *ZBTB4* vari2. These have been designated as oPIDD1, oSFRP4, oPMEPA1, oFAM163B, oC11orf96, uKIAA0753, oBEGAIN, and uZBTB4, respectively (Table 1). Additionally, publicly available Ribo-Seq-based translatomic data for human cell lines also indicated preferential translational initiation at a SEP-coding oORF (*oSCRIB*) (*10*) and LEP-coding oORFs (*oPIDD1* and *oC11orf96*) (fig. S1). Notably, all of the oORFs detected (*oSCRIB*, *Gdown3*, *alt-RPL36*, *oPIDD1*, *oSFRP4*, *oPMEPA1*, *oFAM163B*, *oC11orf96*, and *oBEGAIN*) were +2 out-of-frame relative to their respective refCDSs. Conversely, the SEP-coding uORFs (*uMKKS1* and *uZBTB4*) and an LEP-coding uORF (*uKIAA0753*) were in-frame and +1 out-of-frame with their refCDSs, respectively.

Among these, oSCRIB, Gdown3, uMKKS1, oPIDD1, and uZBTB4 were detected in both SH-SY5Y and A549 cell lines. By contrast, alt-RPL36, oC11orf96, and uKIAA0753 were detected only in SH-SY5Y, while oSFRP4 and oPMEPA1 were found only in A549. The remaining LEP, oFAM163B, encoded as an oORF of a refCDS for the brain-specific protein FAM163B (*19*), was detected only in the brain-related SH-SY5Y cell line. Similarly, the remaining SEP, oBEGAIN, encoded as an oORF of a refCDS for the brain-enriched protein BEGAIN (*20*), was also identified only in SH-SY5Y. These findings highlight cell-type-specific expression of altORFs. Overall, our proteogenomics-based approach in alt-gene discovery effectively refined large alt-gene sequence datasets to promising candidates, indicating the presence of both SEPs and LEPs in human cells.

### Conclusive validation of probability-based identification of new SEPs and LEPs in humans

We previously demonstrated that cell-free production of authentic proteins, followed by MS, accelerates the validation of protein identifications predicted *in silico* by the Mascot engine (*10*). Synthesizing many authentic proteins, particularly their unique tryptic peptide regions, facilitates efficient validation of proteogenomic data mining.

In this study, we engineered two artificial genes encoding a concatemer of many unique tryptic peptides representing each protein (Fig. 1A, B and table S2). These concatenated peptides were synthesized *in vitro* using the Protein synthesis Using Recombinant Elements (PURE) system (*21*), a commercial *in vitro* translation system based on bacterial 70S ribosomes (Fig. 1C). Next, we conducted tryptic digestion of the concatenated peptides that were predominantly detected in the soluble fraction (Fig. 1C). The resulting tryptic peptides, derived from these concatenated peptides, were analyzed via LC-MS/MS under data-dependent acquisition (DDA) control (*22*). The spectra featured profiles of product ions generated through higher-energy collisional dissociation (HCD) (*22, 23*) of the protonated precursor ions from the tryptic peptides (table S2). By manually comparing the spectra with those from corresponding peptides in the previously mentioned proteome datasets (identifier PXD004452), we confirmed equivalent fragmentation patterns (figs. S2–S13). This comparison provided evidence for the existence of the new LEPs (oPIDD1, oSFRP4, oPMEPA1, oFAM163B, oC11orf96, and uKIAA0753) and the new SEPs (oBEGAIN and uZBTB4), as well as the previously reported SEPs (oSCRIB, Gdown3, alt-RPL36, and uMKKS1) in human cells at the protein level (Table 1).

### Common characteristics of the newly discovered SEPs and LEPs in humans

Next, we investigated whether any known proteins exhibit significant sequence similarity to the newly discovered SEPs and LEPs, as well as those SEPs identified previously. This was done by searching the NCBI nonredundant protein sequences and the Universal Protein Resource (UniProt) database (UniProt Knowledgebase) using the Basic Local Alignment Search Tool (BLAST) (*24*). The search results revealed that only uKIAA0753 shares high sequence similarity with known proteins, specifically collagen proteins. The others did not show significant similarity to any known proteins, suggesting that they might have emerged *de novo* from the genomes. Additionally, there was no similarity in nucleotide and translated sequences of these SEPs and LEPs as well as their associated refCDSs, indicating their independent origins.

We further examined whether SEPs and LEPs in human cells share common characteristics. Gene Ontology enrichment analysis of the refCDSs associated with the translatable altORFs encoding SEPs or LEPs identified in this study was conducted using the Database for Annotation, Visualization, and Integrated Discovery (DAVID) (*25*) (Fig. 2A). This analysis showed significant enrichment in cell signaling, with an enrichment score of 1.47—greater than 1.30 threshold for further evaluation (*25*). In fact, the majority of refCDSs (*SCRIB*, *PIDD1*, *SFRP4*, *PMEPA1*, *BEGAIN*, *MKKS*, and *Gdown1*) clustered together, indicating their involvement in cell signaling. Additionally, the results showed that *SCRIB*, *PIDD1*, and *SFRP4*, as well as *MKKS*, *Gdown1*, *ZBTB4*, and *RPL36*, were related to gene expression and regulation; the remaining refCDSs (*FAM163B*, *C11orf96*, and *KIAA0753*) were not clustered in this analysis.

**Fig. 2.**
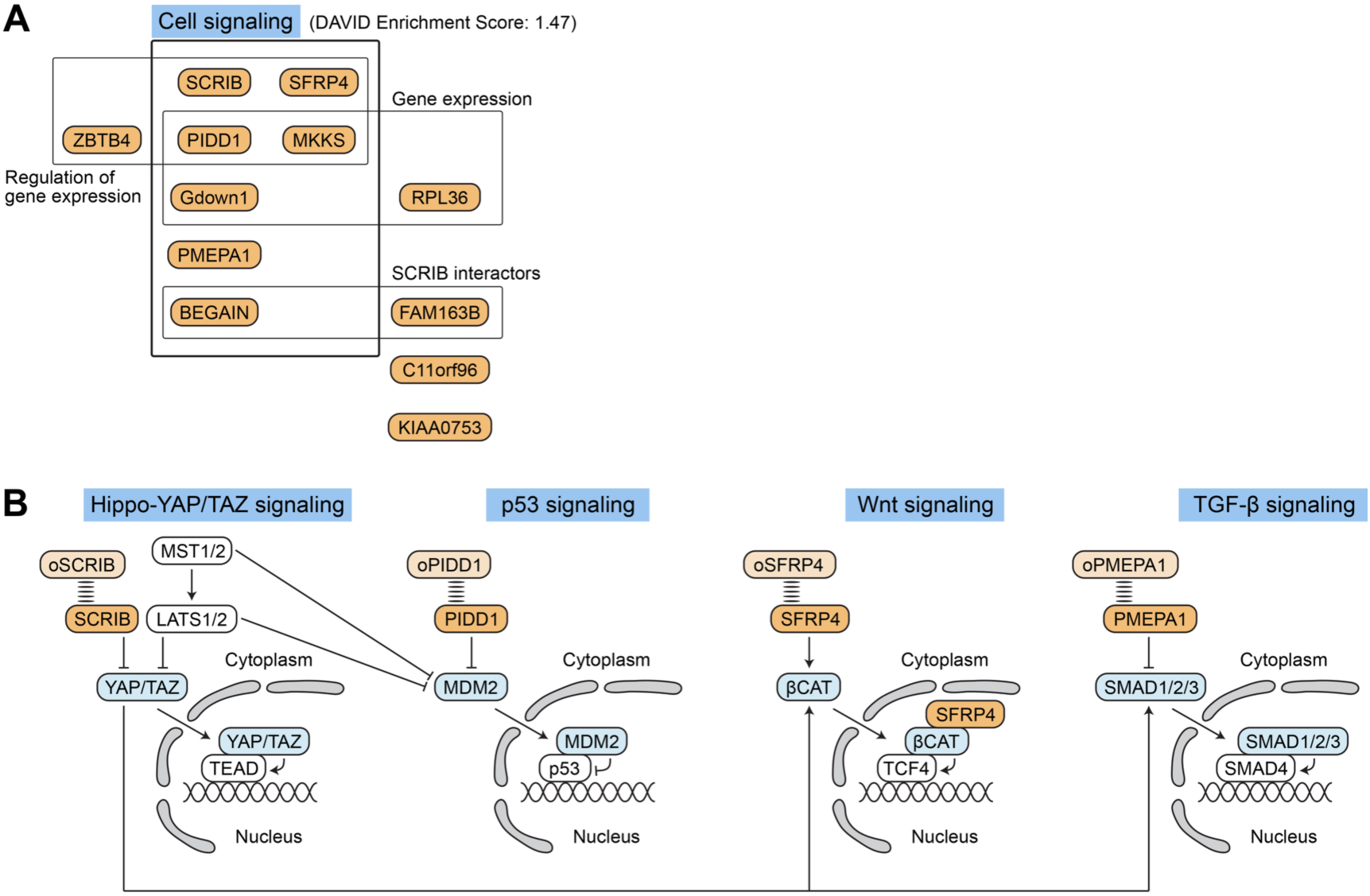
Common characteristics of SEPs and LEPs in humans. (**A**) Gene Ontology enrichment analyses of refCDSs associated with SEP/LEP-coding altORFs (uORFs/oORFs) in humans. Significant enrichment scores (>1.30) calculated by the Database for Annotation, Visualization, and Integrated Discovery (DAVID) are indicated. Potential protein relationships are outlined. (**B**) Signal regulatory functions of refCDSs associated with SEP/LEP-coding oORFs in the Hippo-YAP/TAZ, p53, Wnt, and TGF-β crosstalk pathways in humans. Black dotted lines indicate potential *cis*-regulatory roles of oORFs (*oSCRIB*, *oPIDD1*, *oSFRP4*, and *oPMEPA1*) in the expression of refCDSs (*SCRIB*, *PIDD1*, *SFRP4*, and *PMEPA1*).

Intriguingly, we further observed that the refCDSs with the identified oORFs, particularly those encoding signaling effectors (SCRIB, PIDD1, SFRP4, and PMEPA1), share close relationships. First, the signaling pathways influenced by SCRIB, PIDD1, SFRP4, and PMEPA1, namely, Hippo-YAP/TAZ, p53, Wnt, and TGF-β, are exclusive to multicellular animals (Metazoa) and play pivotal roles in regulating cell–cell interactions, cell proliferation, and body size (*26–28*). Second, the Hippo-YAP/TAZ pathway, in which SCRIB is a key actor, engages in crosstalk with the PIDD1, SFRP4, and PMEPA1 pathways (p53, Wnt, and TGF-β, respectively) (*29, 30*) (Fig. 2B). Third, all of these signaling effectors (SCRIB, PIDD1, SFRP4, and PMEPA1) are known to modulate the nuclear import of transcriptional effectors (coactivators/repressors such as YAP/TAZ, MDM2, βCAT, and SMAD1/2/3) that act on specific transcription factors (TEAD, p53, TCF4, and SMAD4, respectively) (*31–34*) (Fig. 2B). Moreover, FAM163B and BEGAIN, each exhibiting a translatable oORF, were previously identified as potential interactors of SCRIB and SCRIB-interacting DLG, respectively (*20, 35–37*). These shared characteristics of the refCDSs with translatable oORFs suggest that the translatable oORFs play a role in *cis*-regulating the translation of their downstream, biologically significant refCDSs.

### SEP/LEP-coding oORFs function as *cis*-regulatory elements of refCDS translation

In eukaryotic cells, translation initiation involves the 5’-3’ leaky scanning of translational start codons by ribosomes on polycistronic mRNAs (*3*). This process inherently allows ribosomes to skip the start codon of oORFs and reach that of refCDSs, thereby enabling the translation of both oORF and refCDS on the same transcripts. However, because ribosomes translating oORFs cannot simultaneously engage in refCDS translation, oORF translation may impede refCDS translation. Additionally, the start codons of refCDSs are typically embedded within the coding regions of oORFs, potentially leading to ribosomal collision (*38, 39*) and further hindering refCDS translation. Recent studies have indeed shown that SEP-coding oORFs can negatively *cis*-regulate the expression of refCDSs (*40, 41*). We have characterized an SEP-coding oORF, *oSCRIB*, as a negative *cis*-regulatory element affecting *SCRIB* translation (*10*).

To determine whether the translational regulation of refCDSs by their oORFs is common among LEP-coding oORFs, as observed with SEP-coding oORFs (Fig. 3A), we performed *in vitro* expression of DNA fragments encoding a full-length LEP-coding oORF (*oPIDD1* or *oSFRP4*) and a C-terminally truncated, partial refCDS (*PIDD1* or *SFRP4*) fused to the reporter gene *sfGFP* (Fig. 3B, C). The translational efficiency of *PIDD1* and *SFRP4* was measured by the fluorescence intensity of the sfGFP reporter protein. Furthermore, to evaluate the functionality of LEP-coding oORFs (*oPIDD1* and *oSFRP4*), we mutated their translational start codon (AUG) to AGG, a sequence not recognized as a start codon by eukaryotic ribosomes (*42, 43*) (Fig. 3B, C). Changing this to AGG resulted in increased fluorescence intensity of sfGFP (Fig. 3D, E), suggesting that eliminating the translatability of *oPIDD1* and *oSFRP4* increased ribosomal translation of *PIDD1* and *SFRP4*. These findings indicated that LEP-coding oORFs could play a role in *cis*-regulating translation of the respective refCDSs.

**Fig. 3.**
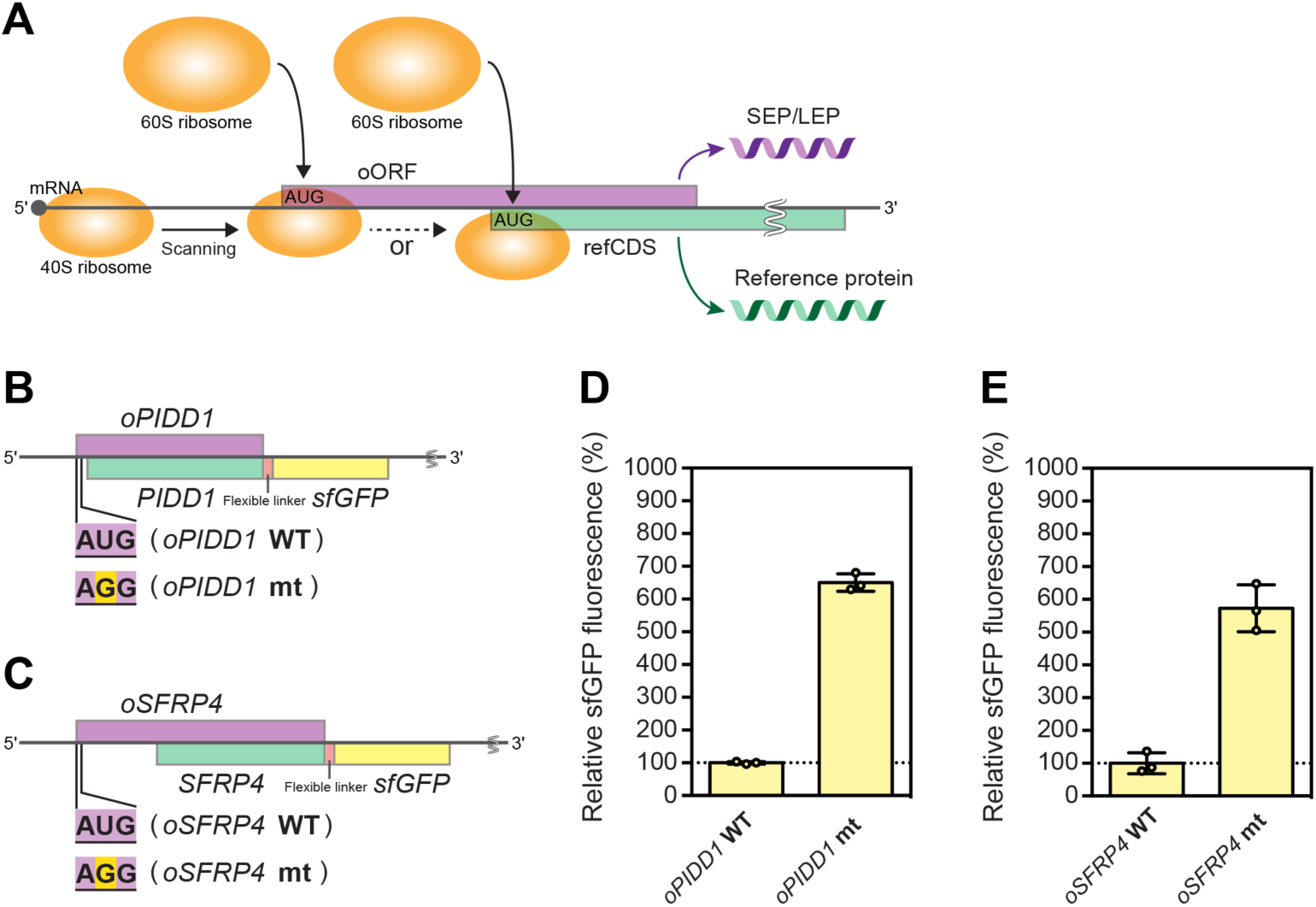
*Cis*-regulatory roles of SEP/LEP-coding oORFs in the translation of signal regulatory refCDSs. (**A**) Proposed bicistronic translation of SEP/LEP-coding oORFs and refCDSs. (**B, C**) Designs of DNAs for *in vitro* transcription and translation of oORFs (*oPIDD1* and *oSFRP4*) and refCDSs (*PIDD1* and *SFRP4*). (**D, E**) Translation efficiency of refCDSs (*PIDD1* and *SFRP4*), which was quantified based on the fluorescence intensity of the reporter protein sfGFP. Data are the mean ± SD of three independent reactions.

### Evolutionary changes in the length of *cis*-regulatory oORFs in mammals

To understand the evolutionary aspects of *cis*-regulatory oORFs, we investigated whether translatable oORFs underwent any evolutionary changes after their origin. First, we observed that oORFs, which are out-of-frame with refCDSs, have coding regions that partly overlap the refCDSs. The degree of overlap appeared to be related to the length of the oORFs, whereas the extent of non-overlap did not show such a correlation. Notably, these oORFs encode not only SEPs but also LEPs, with some oORFs (*alt-RPL36*, *oFAM163B*, and *oC11orf96*) entirely encompassing refCDSs in humans. This led us to the hypothesis that C-terminal extension might be a common evolutionary target for oORFs. To explore this, we compared the length of oORF homologs. We found that the majority of oORFs (*oSCRIB*, *oPIDD1*, *oSFRP4*, *oPMEPA1*, *oFAM163B*, *oBEGAIN*, and *oC11orf96*) have significantly elongated C-terminals after their origin as *cis*-regulatory oORFs, incorporating downstream regions that are out-of-frame with refCDSs during animal evolution (Fig. 4 and figs. S14A, S15A, S16A, and S17). The remaining oORFs (*Gdown3* and *alt-RPL36*), which were previously found only in specific transcript variants (vari1/2 and vari2, respectively), did not show such a trend.

**Fig. 4.**
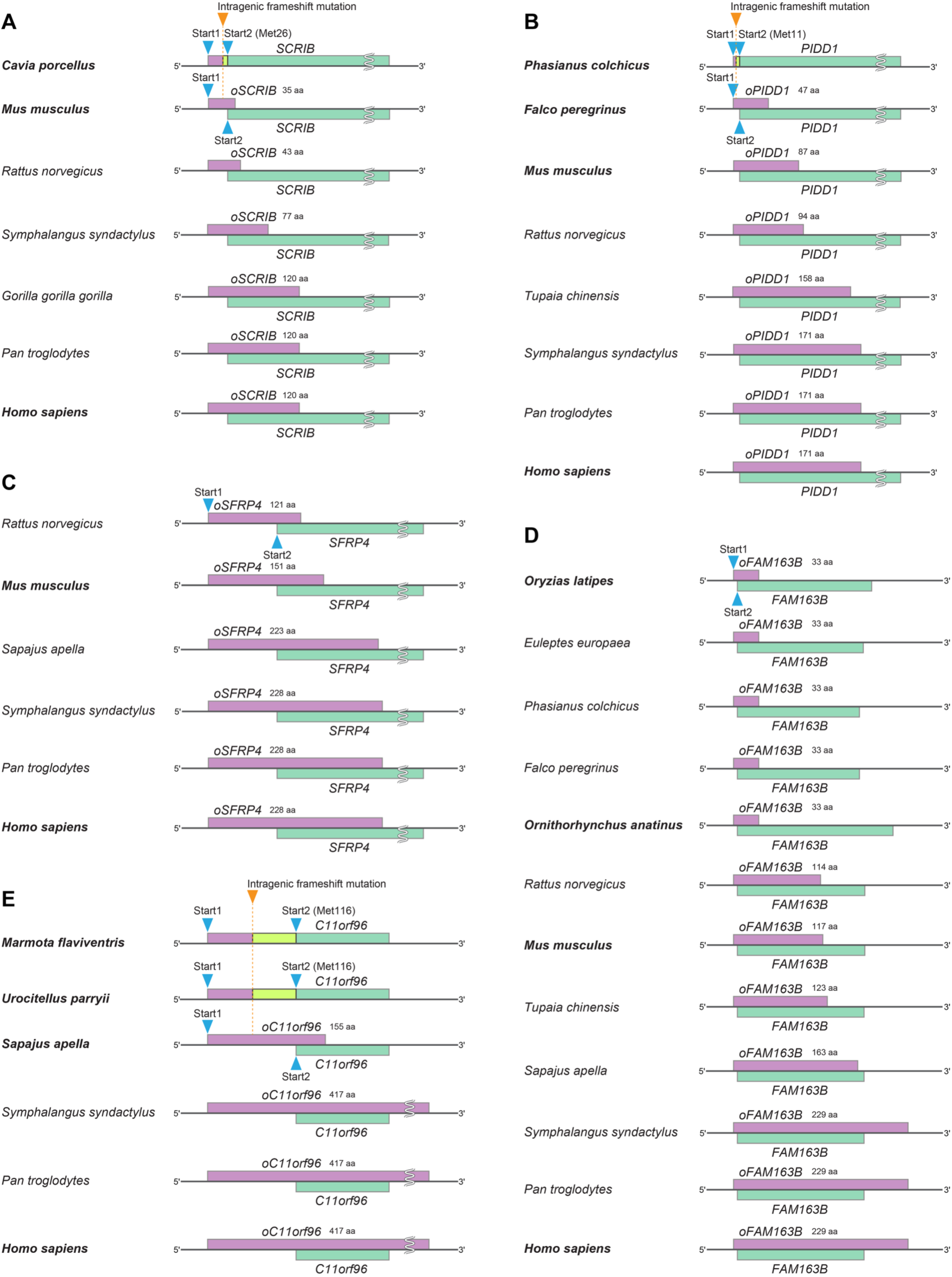
Evolution of SEP/LEP-coding oORFs by C-terminal extension. (**A–E**) Observed evolution of SEP/LEP-coding oORFs in animals. Blue arrowheads indicate translational start codons; orange arrowheads indicate frameshift sites.

Our results demonstrate that large C-terminal extensions occurred during the evolution of animals, particularly in mammals; both *oFAM163B* and *oC11orf96* came to encompass their associated refCDSs during primate evolution (Fig. 4 and figs. S14A, S15A, S16A, and S17). The oORFs, including human oSCRIB (120 amino acids), oPIDD1 (171 amino acids), and oC11orf96 (417 amino acids), are >3, >3.5, and >2.5 times longer than their homologs in *Mus musculus* (35 amino acids), *Falco peregrinus* (47 amino acids), and *Sapajus apella* (155 amino acids), respectively. Human oFAM163B (229 amino acids) is nearly 7 times longer than its homologs (33 amino acids) in fish (*Oryzias latipes*), reptiles (*Euleptes europaea*), birds (*Phasianus colchicus* and *Falco peregrinus*), and mammals (*Ornithorhynchus anatinus*). These findings indicate that *cis*-regulatory oORFs have undergone C-terminal extension during evolution.

### C-terminal extension of oORFs enhances their *cis*-regulatory impact on refCDS translation

Because oORF-translating ribosomes are not simultaneously available for refCDS translation, longer oORFs should prolong the duration of ribosome occupancy, thereby reducing the frequency of refCDS translation. Consequently, the *cis*-regulatory effect of oORFs on refCDS translation may be amplified by C-terminal extension. To test this hypothesis, we expressed plasmid DNAs encoding two fluorescent reporter genes (*AmCyan1* and *DsRed-Express*) in human embryonic kidney cells (HEK293T). *AmCyan1* was engineered to contain an oORF of 14, 31, 60, 182, or 244 amino acids, without any change in the amino acid sequence of *AmCyan1* itself, whereas *DsRed-Express* was used as an internal control to evaluate the transfection efficiency of the plasmid DNAs (Fig. 5A). The translational efficiency of the reporter genes was measured by fluorescence intensity. This measurement showed that the introduction of a longer oORF into *AmCyan1* resulted in lower fluorescence intensity of *AmCyan1* (Fig. 5B), indicating a decrease in its translational efficiency due to longer oORF translation. These findings suggest that longer oORFs due to C-terminal extension have an enhanced *cis*-regulatory effect on refCDS translation in the cells.

**Fig. 5.**
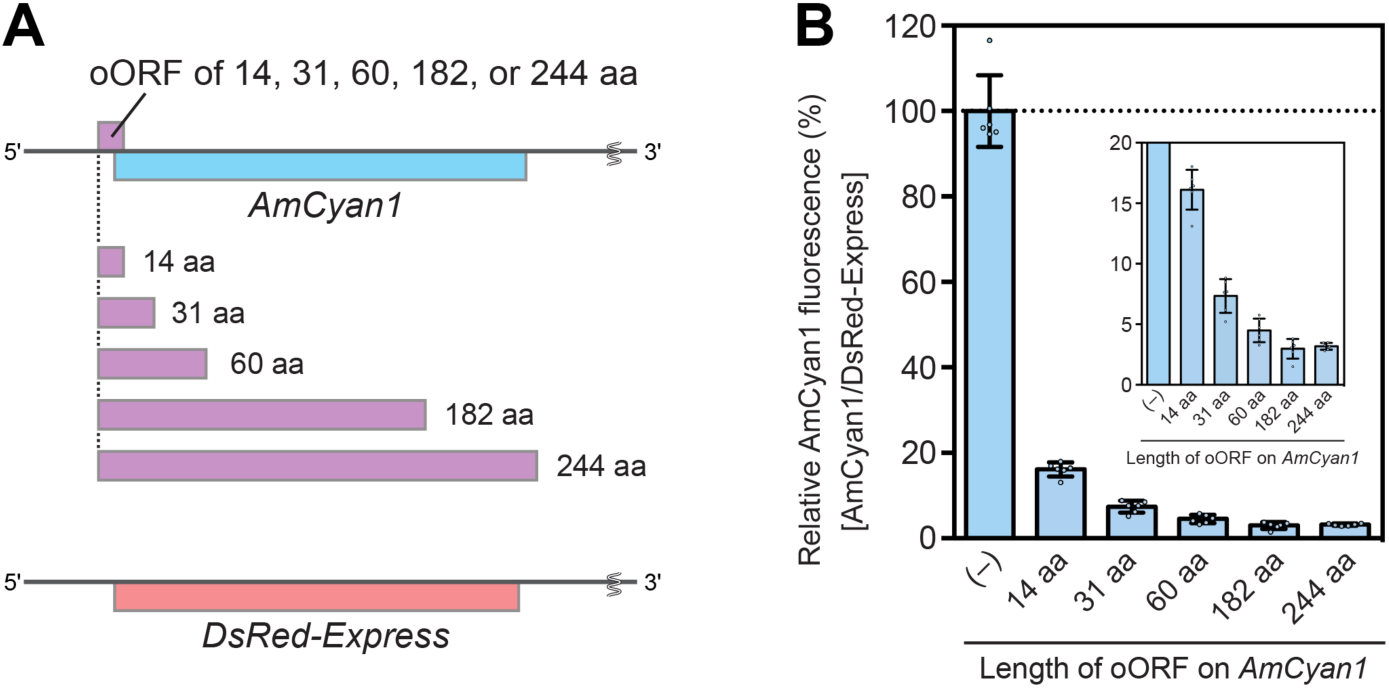
*cis*-Regulatory impact of C-terminally extended oORFs on refCDS translation. (**A**) Design of plasmid DNAs encoding two fluorescent reporter genes, *AmCyan1* and *DsRed-Express*, expressed in human embryonic kidney cells (HEK293T). *AmCyan1* contained an oORF of 14, 31, 60, 182, or 244 amino acids. *DsRed-Express* was used as an internal control to evaluate plasmid DNA transfection efficiency. (**B**) Translation efficiency of *AmCyan1* with an oORF, quantified by fluorescence intensity. Data are the mean ± SD of six independent cell cultures.

### Longer oORFs are frequently +2 out-of-frame relative to refCDSs

Our proteogenomic study revealed the existence of *cis*-regulatory oORFs that have undergone C-terminal extension during evolution and consequently code for LEPs in humans. Intriguingly, all of them were +2 out-of-frame relative to refCDSs. Accordingly, we wondered whether longer oORFs are frequently +2 out-of-frame relative to refCDSs. To address this question, we inspected the results of the Ribo-Seq-based translatomic analysis conducted by Yang et al. (*44*), which serves as an important complement to our proteogenomic study. We first retrieved translatable oORFs that start with AUG and end with stop codons from human translatomic data. Of the 335 oORFs retrieved, only eight— including oPIDD1—coded for LEPs ≥150 amino acids, whereas 292 oORFs were SEPs of <100 amino acids and 35 were ≥100 but <150 amino acids (data S1). We then evaluated the ratio of +2 out-of-frame oORFs to +1 out-of-frame oORFs. Although the numbers of +1 and +2 out-of-frame oORFs were equal (50:50) for oORFs encoding SEPs <100 amino acids, the ratio of +2 out-of-frame oORFs to +1 out-of-frame oORFs increased with oORF length (60% for oORFs encoding SEPs ≥100 but <150 amino acids and 75% for LEPs ≥150 amino acids) (fig. S18 and data S1). These data support our hypothesis that longer oORFs are frequently +2 out-of-frame relative to refCDSs.

### Variable proteoform and translatability of oORFs by genomic dynamics

Additionally, when we searched for homologs of oSCRIB and oPIDD1 in rodents (order Rodentia) and birds (class Aves), respectively, with the shortest oORFs in our dataset, we observed that *Cavia porcellus* (Rodentia) and *Phasianus colchicus* (Aves) have unique refCDSs (SCRIB and PIDD1, respectively) fused with partial oORFs (oSCRIB and oPIDD1, respectively) (Fig. 4A, B and figs. S14A–C and S15A–C). Similarly, a partial oC11orf96 was fused with C11orf96 in *Marmota flaviventris* (Rodentia) and *Urocitellus parryii* (Rodentia) (Fig. 4E and fig. S16A–C). Although it is difficult to trace ancient gene birth/loss events, these cases—suggesting a common mechanism for the appearance and disappearance of oORFs—resulted from frameshift mutations between the start codons of oORFs and refCDSs (figs. S14D, S15D, and S16D). These findings indicate that translatable oORFs are strongly influenced by genomic alterations.

Therefore, we investigated whether genetic variation was present in the coding regions of the identified oORFs. Using NCBI dbSNP (*45*), a database of genetic variation, such as single nucleotide polymorphisms (SNPs) and short insertions and deletions (indels), we assessed whether SEP/LEP-coding oORFs—including human *oSCRIB*, *oPIDD1*, *oSFRP4*, *oFAM163B*, and *oC11orf96*—are affected by known genetic variation. Our search confirmed the existence of oORFs elongated by SNPs that eliminate oORF stop codons, as well as oORFs shortened by single nucleotide insertions that shift the downstream reading frames (Fig. 6). We also observed the disappearance of oORFs due to single nucleotide deletions causing a +1 frameshift and fusing them with the refCDSs (Fig. 6A, C, E). These findings further demonstrate the variable proteoform and translatability of oORFs due to genomic dynamics, which is a driving force behind the formation of taxonomically restricted genes, or orphan genes (*46*).

**Fig. 6.**
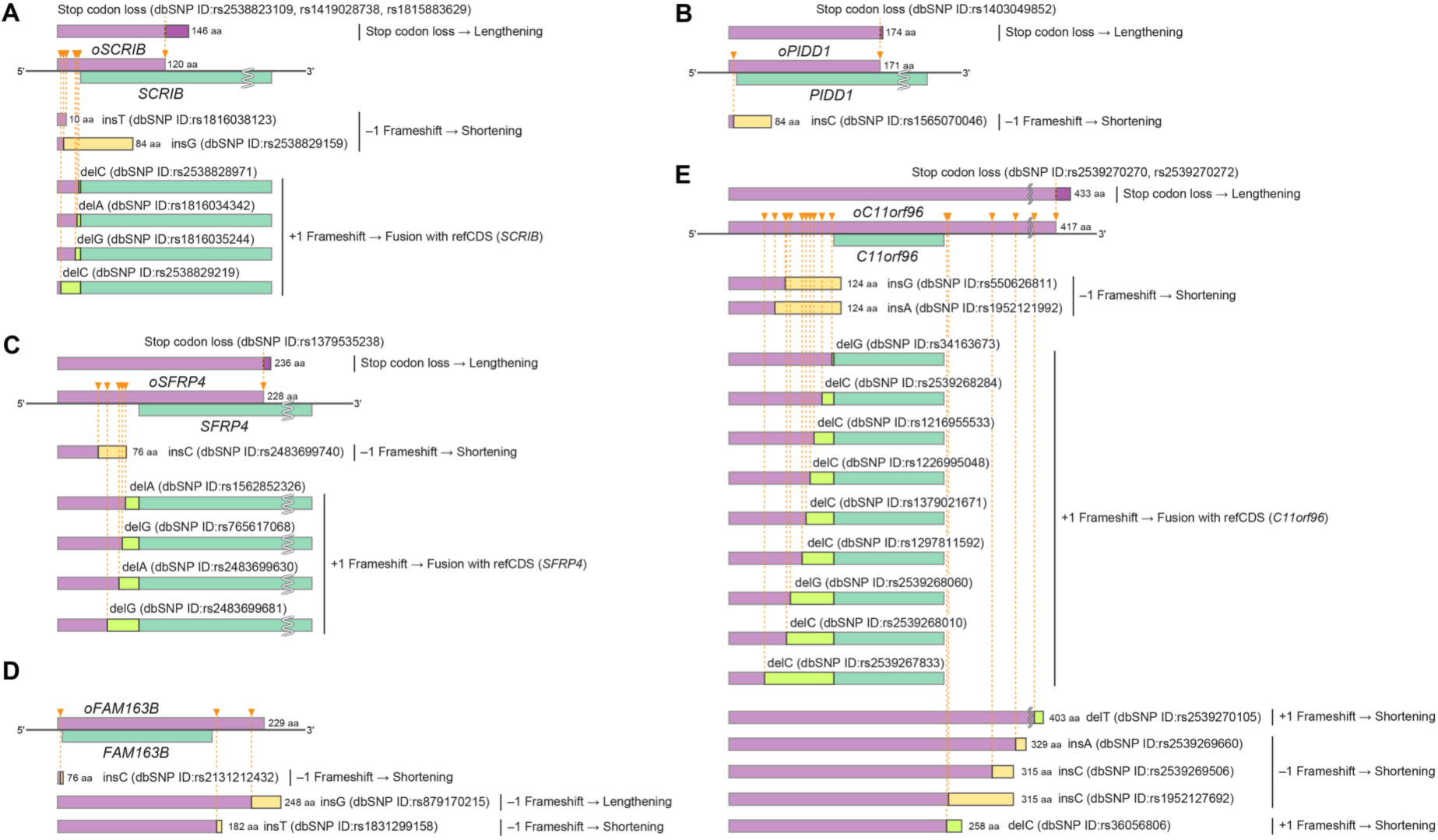
Variable proteoform and translatability of oORFs due to genetic variation in humans. (**A–E**) Detected genetic variations, including single nucleotide polymorphisms (SNPs) and short insertions and deletions (indels), that alter the coding regions of SEP/LEP-coding oORFs in humans. Positions of SNPs and indels retrieved from NCBI dbSNP are indicated by orange arrowheads.

### Prevalence of overlapping gene birth in humans due to frameshift polymorphisms

Next, we considered how translatable oORFs emerged in animal genomes. The *de novo* birth of oORFs has previously been explained by two distinct modes (*47–49*). The first mode involves gene overlap between a refCDS and its uORF through terminal extensions. This overlap can occur through the conversion of uORFs into oORFs by C-terminal extension, either via SNPs that eliminate uORF stop codons or indels within uORF regions that shift their downstream reading frames. It may also arise from N-terminal extension of refCDSs through the creation of new refCDS start codons within uORF regions. The second mode could originate from intergenic start codon-forming mutations located upstream, but near refCDSs, leading to the *de novo* creation of oORFs that use a downstream stop codon out-of-frame with the refCDSs.

In addition to these two modes of origin, we observed that intragenic frameshift mutations within refCDS regions also create oORFs, which could represent a new third mode of origin. Intragenic frameshift mutations within refCDS regions would split the refCDSs at the mutation sites, leading to the emergence of oORFs that usurp the start codons of the refCDSs, along with surrounding sequences, including the Kozak sequence (*50, 51*). The resulting oORFs would then incorporate N-terminal coding regions from the refCDSs located upstream of the frameshift site, while exploiting their downstream regions that are out-of-frame with the refCDSs. This process could lead to the emergence of N-terminally truncated refCDSs, which may use preexisting or potentially usable internal start codons located near, yet downstream from, the frameshift sites (*52, 53*). The newly formed oORFs, partially composed of sequences from the original refCDSs, are then destined to *cis*-regulate the refCDS translation. Based on their origin and functionality, we propose that these SEP- and LEP-coding oORFs be designated as <u>ups</u>tream region-<u>u</u>surping <u>r</u>epurposed <u>p</u>roteins (USURPs).

Using NCBI ClinVar (*54*), a database of genetic variants in humans—particularly those related to diseases—we investigated whether translatable oORFs, specifically *USURPs* with intragenic frameshifts in refCDSs, emerge *de novo* from genomes. We focused on identifying *USURPs* emerging through a –1 frameshifting single nucleotide insertion near the start codons of refCDSs (Fig. 7A), based on our observation of the opposite event, *i.e.*, disappearance of oORFs via +1 frameshifting single nucleotide deletions (Fig. 6A, C, E). We first extracted human nucleotide sequences meeting this criterion and identified eighteen candidate refCDSs reported to harbor pathogenic single nucleotide insertions within their first 49 amino acids, without exceeding N-terminal truncations of 49 amino acids (Fig. 7B and data S2). These candidates revealed the emergence of 9 SEP-coding uORFs (ranging from 10 to 29 amino acids), 8 SEP-coding oORF (*USURPs*; 35–98 amino acids), and an LEP-coding *USURP* (166 amino acids) in humans. This indicates the frequent occurrence of *USURPs* due to frameshift polymorphisms in the human genome.

**Fig. 7.**
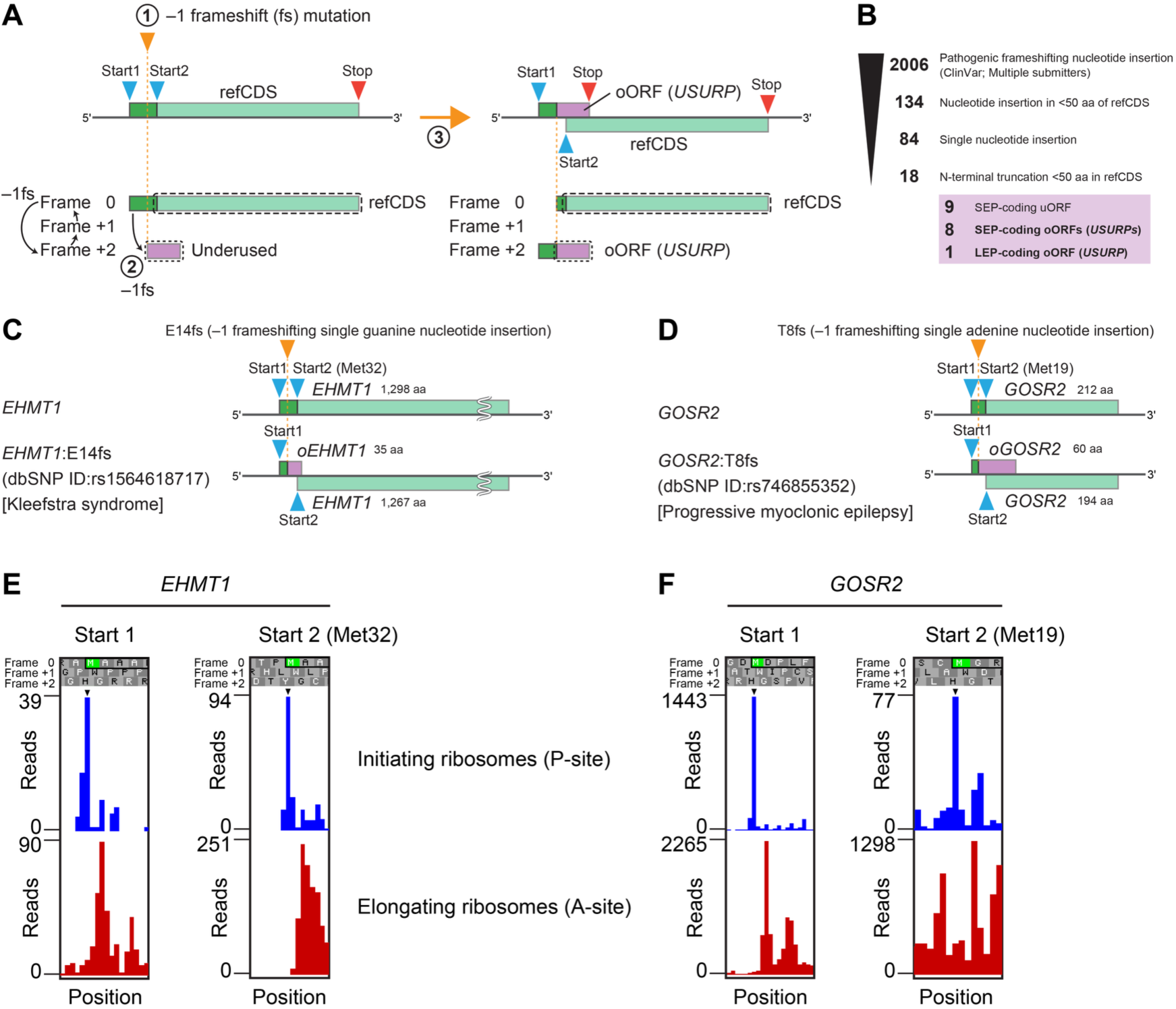
Emergence of *USURP* oORFs in humans by disease-associated frameshift polymorphisms. (**A**) Proposed mode of emergence of translatable oORFs (*USURPs*) through intragenic frameshifts in refCDSs. Positions of translational start and stop codons are indicated by blue and red arrowheads, respectively. Frameshift sites are indicated by orange arrowheads. (**B**) Data mining scheme of NCBI ClinVar resulting in the identification of oORFs (*USURPs*) in human *EHMT1* and *GOSR2* via pathogenic frameshift polymorphisms. (**C, D**) Representative *USURP* oORFs (*oEHMT1* and *oGOSR2*) that emerged from human disease-related refCDSs (*EHMT1* and *GOSR2*) through frameshift polymorphisms. Positions of translational start codons and frameshift sites are indicated by blue and orange arrowheads, respectively. (**E, F**) Survey of public Ribo-Seq data. Aggregated profiles of initiating and elongating ribosomes were obtained from GWIPS-viz (https://gwips.ucc.ie).

Among these, two new *USURPs* were identified, both coding for SEPs of 35 and 60 amino acids. These emerged from the refCDSs for euchromatic histone-lysine *N*-methyltransferase 1 (EHMT1; 1,298 amino acids) and Golgi soluble *N*-ethylmaleimide-sensitive factor attachment protein (SNAP) receptor (SNARE) complex member 2 (GOSR2; 212 amino acids) through –1 frameshifting single nucleotide insertions of guanine and adenine (E14fs and T8fs), respectively (Fig. 7C, D). After their emergence, the expression of the EHMT1 and GOSR2 refCDSs, implicated in Kleefstra syndrome (*55*) and progressive myoclonic epilepsy (*56*), could potentially be compensated for by internal start codons downstream of the frameshift sites. Notably, the possible internal start codons for *EHMT1* and *GOSR2* were identified at ATG_94-96_ (Met32) and ATG_55-57_ (Met19), respectively, leading to the production of proteins truncated by 31 and 18 amino acids (Fig. 7C, D).

Supporting the functional potential of the internal start codon of *EHMT1*, ATG_94-96_ (Met32), publicly available Ribo-Seq-based translatomic data indicated efficient translational initiation at both the primary (ATG_1-3_) and internal start codons in human cell lines (Fig. 7E). Additionally, transcript variant E of *GOSR2* is known to utilize ATG_55-57_ (Met19) as a start codon (NCBI accession number NM_001321134), in line with the translatomic data that demonstrated its availability (Fig. 7F). These findings highlighted the translatability of not only the newly emerged *USURPs* but also the N-terminally truncated refCDSs. Therefore, the pathogenicity of frameshift polymorphisms in *EHMT1* and *GOSR2* likely results not from the disruption and untranslatability of refCDSs but from reduced expression of refCDSs due to *USURP* translation. Overall, our research reveals that *USURPs* are currently emerging in human genomes and suggests the involvement of these *cis*-regulatory *USURPs* in human diseases.

## Discussion

We expanded our investigation into newly active altORFs on human mRNAs using a proteogenomic approach, complemented by validation of their expression using artificially concatenated peptides for the release of multiple peptide standards. These methodologies enabled us to recognize that the coding regions of these altORFs, particularly oORFs, encompass not only short ORFs encoding SEPs <150 amino acids but also longer altORFs encoding LEPs ≥150 amino acids.

A recent study highlighted the presence of uORFs involved in the translational regulation of refCDSs within the Hippo-YAP/TAZ and Wnt signaling pathways (*57*). We expanded on this by revealing that SEP/LEP-coding oORFs are present in transcripts encoding biologically important proteins, including signal effectors of key crosstalk pathways such as Hippo-YAP/TAZ, p53, Wnt, and TGF-β in humans. This expanded group of translatable oORFs is characterized by the inclusion of LEP-coding longer altORFs.

AltORFs, particularly oORFs, have traditionally been assumed to be short as stop codons are believed to accumulate more frequently in reading frames than in those coding for functional proteins (refCDSs). This accumulation of premature stop codons was thought to limit the length of oORFs. However, our recent discoveries challenge this notion. We found that newly identified oORFs exhibit a higher tolerance to truncation mutations than previously anticipated and predominantly code for LEPs. These oORFs have been evolutionarily selected for C-terminal extension through the elimination of stop codons, indicating that C-terminal extension is a common evolutionary target for these sequences.

Our research also demonstrated that translatable oORFs play a role in the *cis*-regulation of refCDS translation and their *cis*-regulatory impact is amplified by the C-terminal extension of the oORFs. In scenarios where stronger *cis*-regulation is advantageous, this could serve as an evolutionary driving force, leading to natural positive (Darwinian) selection favoring the C-terminal extension for its beneficial effects. This phenomenon may also present opportunities to acquire cellular functions via the +2 underused frame of refCDSs. Indeed, three-dimensional structural modeling and functional domain analysis using AlphaFold2 and InterProScan revealed that human oC11orf96 contains a short α-helical structure, which potentially forms a transmembrane domain in its C-terminal region (fig. S19).

We found that longer oORFs are frequently +2 out-of-frame relative to refCDSs. Accordingly, we recognized that +2 out-of-frame oORFs are more amenable to C-terminal extension than +1 out-of-frame oORFs. The loss of translational stop codons (TAA, TAG, and TGA) in +2 out-of-frame oORFs can occur without impacting their associated refCDSs, because the first thymine base of these stop codons is a wobble base in the frame for refCDSs (fig. S20A). Changing this first thymine to adenine, cytosine, or guanine results in the loss of the stop codons. In fact, we observed frequent removal of stop codons due to mutations in this first thymine (fig. S20B). Conversely, for +1 out-of-frame oORFs, a change in the second adenine or guanine base of the stop codons to cytosine or thymine is necessary (fig. S20C), which limits the likelihood of losing these stop codons. Moreover, in single-nucleotide mutations, transition between thymine and cytosine (*i.e.*, one-ring pyrimidines) or between adenine and guanine (*i.e.*, two-ring purines) is more common than transversion between one-ring and two-ring structures (*58*). This reduces the frequency of eliminating the stop codons in +1 out-of-frame oORFs, as the second adenine or guanine base of the stop codons is frequently swapped between adenine and guanine (*i.e.*, two-ring purines), which does not abolish these codons (fig. S20C). This constraint on the C-terminal extension of +1 out-of-frame oORFs may reduce the prevalence of LEP-coding longer oORFs that are +1 out-of-frame with refCDSs. This also aligns with reports of SEP-coding short altORFs nested within refCDSs, often occurring +1 out-of-frame with refCDSs (*59, 60*).

We also observed that translatable oORFs coding for SEPs and LEPs are strongly influenced by genomic changes, such as SNPs and indels; their birth/loss events frequently involve intragenic frameshift mutations in humans. Supporting our concept of upstream region-snatching *USURPs* as a third mode of overlapping gene birth, we revealed the frequent occurrence of *USURPs* due to frameshift polymorphisms in humans, some of which are closely linked to human diseases. The one-step birth of *USURPs* via intragenic frameshifts in refCDSs circumvents the need for evolutionary enhancement of Kozak sequence contexts, thereby explaining the efficient translation of *USURPs*. Although the concurrent N-terminal truncation of refCDSs—especially when the truncations are large—is potentially detrimental to protein functionality and likely subject to natural negative (or purifying) selection that hinders the spread of harmful alleles, previous studies observed that potentially usable start codons are abundant near the primary start codons of eukaryotic proteins, including those in humans (*52, 53*). These preexisting internal start codons of refCDSs are believed to have evolved for the generation of different proteoforms or to ensure the expression of refCDSs, particularly their core regions (*61*). This fundamental characteristic of refCDSs appears to facilitate the emergence of *USURPs* without compromising the translatability of refCDSs. Traditionally, frameshift polymorphisms are considered detrimental mutations that disrupt refCDSs and their translatability. However, our findings suggest the existence of human diseases stemming from the emergence of *cis*-regulatory *USURPs* without compromising the translatability of refCDSs using preexisting internal start codons.

In conclusion, our investigation extended beyond SEPs to include LEPs, uncovering previously overlooked dark proteomes. We demonstrated that the *de novo* birth and evolution of proteomes have been driven by genome dynamics, thereby enriching the protein universe. Their evolution by C-terminal extension is unique among proteins encoded in refCDSs. Additionally, the one-step birth of *USURPs* facilitates introduction of an additional layer of *cis*-translational regulation. These findings underscore the high density of genetic information in the human genome and fuel our ongoing research into alt-gene discovery, which may prove invaluable in identifying future drug targets.

## Materials and Methods

### Transcriptomic data processing

We utilized publicly available paired-end RNA-Seq data of polyadenylated mRNAs from SH-SY5Y and A549 cell lines, including three biological replicates for each cell line. The data were sourced in two formats: as a single Sequence Read Archive (SRA) file or a set of FASTQ files. Specifically, we obtained paired-end SRA format data of SH-SY5Y cells (with 2.25× 10^7^, 2.22× 10^7^, and 9.2× 10^7^ read pairs) from the NCBI BioProject (accession PRJNA301726 and PRJNA352080); paired-end SRA format data of A549 cells (1.61 × 10^8^ read pairs) from the NCBI BioProject (accession PRJNA523380) by the Cancer Cell Line Encyclopedia (CCLE); and paired-end FASTQ format data of A549 cells (7.11 × 10^7^ and 7.72 × 10^7^ read pairs) from the Encyclopedia of DNA Elements (ENCODE) database (https://www.encodeproject.org). The SRA data were converted to FASTQ files using the fasterq-dump utility from the NCBI SRA Toolkit 2.9.6-1. Then we processed the raw paired-end reads in the FASTQ files using Trim Galore! 0.6.2 for trimming, followed by *de novo* assembly of the trimmed reads with Trinity 2.8.5 using default settings.

To identify potential ORFs within the assembled sequences and human RefSeq Transcripts (NCBI RefSeq assembly accession GCF_000001405.39), we used TransDecoder.LongOrfs 5.5.0, setting a parameter for a minimum codon length of 10 (*i.e.*, a minimum protein length of 9 amino acids). Then we extracted all possible short/longer ORFs starting and stopping with AUG and stop codons, respectively, using SeqKit 0.12.0. The aggregated datasets of shorter and longer ORFs from the three replicates of each cell line were combined into a nonredundant dataset for each cell line. These translated nucleotide sequences of short/longer ORFs, representing amino acid sequences of putative SEPs and LEPs, were further compared against the human RefSeq Proteins (NCBI RefSeq assembly accession GCF_000001405.39) using BLAST+ 2.9.0. The resulting datasets, including SEP/LEP sequences and their protein annotations from BLAST+, were combined with SeqKit. Only the unannotated sequences were retained in each dataset (for SH-SY5Y, A549, and RefSeq Transcripts). These datasets included millions of short ORFs potentially encoding SEPs (4,669,292 in SH-SY5Y, 1,029,024 in A549, and 1,146,023 in RefSeq Transcripts) and thousands to tens of thousands of longer ORFs, *i.e.*, longer altORFs potentially encoding LEPs (8,602 in SH-SY5Y, 5,984 in A549, and 10,181 in RefSeq Transcripts). To improve the accuracy of proteogenomics-based identification of SEPs and LEPs in humans, the dataset of RefSeq Transcripts was integrated into that of each cell line, with the resultant nonredundant datasets serving as the custom SEP/LEP sequence databases for SH-SY5Y and A549 cells.

### DNA construction for in vitro transcription and translation

We synthesized DNA fragments corresponding to concatenated peptides-coding genes (table S3) and *sfGFP* (*10*), as well as constructs *oPIDD1*-*PIDD1*-*sfGFP* and *oSFRP4*-*SFRP4*-*sfGFP* (table S3). These fragments were introduced into a pEX-A2J2 vector by Eurofins Genomics (Tokyo, Japan), and their sequences were confirmed by the same company. Then we amplified the coding regions along with their 3’-untranslated regions (3’-UTRs) derived from the pEX-A2J2 vector. This amplification was carried out using polymerase chain reaction (PCR) with Tks Gflex DNA Polymerase (Takara Bio, Shiga, Japan). Specific forward and reverse primers were used, as listed in table S4 and in our previous paper (*10*): Fw1-C and Rv1 for concatenated peptide-coding genes, Fw1-G and Rv1 for *sfGFP*, Fw1-PG1 and Rv1 for *oPIDD1* (WT; ATG^start^)-*PIDD1*-*sfGFP*, Fw1-PG2 and Rv1 for *oPIDD1* (mt; ATG^start^ to AGG)-*PIDD1*-*sfGFP*, Fw1-SFG1 and Rv1 for *oSFRP4* (WT; ATG^start^)-*SFRP4*-*sfGFP*, and Fw1-SFG2 and Rv1 for *oSFRP4* (mt; ATG^start^ to AGG)-*SFRP4*-*sfGFP*. The first PCR products were used as templates for a second PCR with primers as listed in our previous paper (*10*): split forward primers, Fw2-E and Fw3-E (3 and 300 nM, respectively), and a nested reverse primer Rv2 (300 nM) for concatenated peptide-coding genes and *sfGFP*, and other split forward primers, Fw2-W and Fw3-W (3 and 300 nM, respectively), and the nested reverse primer Rv2 (300 nM) for *oPIDD1* (WT and mt)-*PIDD1*-*sfGFP* and *oSFRP4* (WT and mt)-*SFRP4*-*sfGFP*. The resulting second PCR products were purified and concentrated using the Illustra GFX PCR DNA and gel band purification kit (Cytiva, Tokyo, Japan). To confirm their molecular sizes, the DNA fragments underwent 1% (w/v) agarose gel electrophoresis and were visualized under ultraviolet light after staining with GelRed nucleic acid gel stain (FUJIFILM Wako Chemicals, Osaka, Japan). These DNA fragments were subsequently used for *in vitro* transcription and translation.

### DNA construction for protein expression in human cells

Two DNA fragments were amplified from the original pDsRed-Express-N1 (Clontech, Mountain View, CA, USA) using two pairs of primers with small modifications necessary for DNA construction: DsRed-Fw1 & DsRed-Rv1 and DsRed-Fw2 & DsRed-Rv2 (table S4). Amplification was performed by PCR using PrimeSTAR Max DNA Polymerase (Takara Bio). The two DNA fragments were reassembled into a circular plasmid using the NEBuilder HiFi DNA Assembly Kit (New England Biolabs, Ipswich, MA, USA). The modified pDsRed-Express-N1 was then sequenced.

Next, the modified pDsRed-Express-N1 and the original pAmCyan1-N1 (Clontech) were used as PCR templates for *DsRed-Express* and *AmCyan1*, respectively. Specific forward and reverse primers were used, as listed in table S4: DsRed-Fw3 & DsRed-Rv3 for *DsRed-Express* and AmCyan-Fw & AmCyan-Rv for *AmCyan1*. The two DNA fragments were reassembled into a circular plasmid using the NEBuilder HiFi DNA Assembly Kit. The resulting pAmCyan1-DsRed-Express plasmid was then sequenced.

To introduce an oORF into *AmCyan1*, pAmCyan1-DsRed-Express was digested with *Sal*I (Takara Bio) and *Age*I-HF (New England Biolabs); the resultant DNA fragment was mixed with a single-stranded oligonucleotide, oORF-244aa-Fw (table S4), for assembly using the NEBuilder HiFi DNA Assembly Kit. Introduction of the oORF into *AmCyan1* was confirmed by sequence analysis. The length of the oORF in *AmCyan1* was then modified by site-directed mutagenesis to introduce a new stop codon, using four pairs of primers: oORF-14aa-Fw & oORF-14aa-Rv; oORF-31aa-Fw & oORF-31aa-Rv; oORF-60aa-Fw & oORF-60aa-Rv; and oORF-182aa-Fw & oORF-182aa-Rv (table S4). These plasmids were then sequenced.

To prepare an empty vector lacking *AmCyan1* and *DsRed-Express*, the modified pDsRed-Express-N1 was digested with *Mlu*I (Takara Bio) and *Asc*I (New England Biolabs), then subjected to self-ligation using Ligation high Ver.2 (Toyobo, Osaka, Japan). The resulting pEmpty plasmid was then sequenced.

### In vitro transcription and translation

Bacterial transcription coupled with translation was conducted *in vitro* using the PURE*frex* 2.0 system (GeneFrontier, Chiba, Japan). The reaction was initiated by adding 5 μL DNA fragment (either concatenated peptides-coding genes or *sfGFP*) into a 15 μL mixture of RNase-free water and PURE*frex* 2.0 Solutions I, II, and III. The mixtures were incubated at 37°C for 5 h. After incubation, the mixtures were centrifuged at 20,400 × *g* for 20 min at 4°C. The resulting supernatants (soluble fractions) and pellets (insoluble fractions) were analyzed by sodium dodecyl sulfate-polyacrylamide gel electrophoresis (SDS-PAGE), followed by staining with Coomassie Brilliant Blue G-250. For eukaryotic translation, the WEPRO7240H Expression Kit (CellFree Sciences, Matsuyama, Japan) was used as described previously (*10*) with minor modifications. The reaction mixtures were then utilized to measure the fluorescence intensity of the reporter protein sfGFP at excitation/emission wavelengths of 485/535 nm using a Wallac 1420 Multilabel Counter ARVO MX (PerkinElmer, Waltham, MA, USA).

### Sample preparation for LC-MS/MS analysis

The protein bands corresponding to *in vitro* synthesized concatenated peptides (artificial proteins) were excised from the SDS-PAGE gels. These gel slices were destained and subjected to reduction and alkylation (carbamidomethylation) treatments, first with 50 mM dithiothreitol and then with 100 mM sodium iodoacetate. Following this, the gel slices were washed with water. Then the proteins within the gel were digested using *N*-tosyl-L-phenylalanine chloromethyl ketone-treated trypsin (Worthington Biochemical, Freehold, NJ, USA). This was carried out in a solution containing 20 mM Tris-HCl buffer (pH 8.0) and 0.05% (w/v) *n*-dodecyl-β-D-maltoside at 37°C for 16 h. The resulting peptides were prepared for analysis by liquid chromatography-tandem MS (LC-MS/MS).

### LC-MS/MS analysis

The peptide samples were applied to a packed nano-capillary C18 column (NTCC-360/75-3-105, 0.075 × 105 mm, 3 μm particle size; Nikkyo Technos, Tokyo, Japan) for LC-MS/MS analysis using an Easy-nLC 1000 LC system (Thermo Fisher Scientific, Waltham, MA, USA). Two types of eluents were used: eluent A (water with 0.1% (v/v) formic acid) and eluent B (acetonitrile with 0.1% (v/v) formic acid). The column temperature was maintained at room temperature, and eluent A was initially applied at a flow rate of 300 nL/min. Elution involved increasing the proportion of eluent B from 0–100% over 12 min, reaching 35% at 10 min and 100% at 12 min, and this final condition was held for an additional 8 min. The eluate was analyzed using a Q Exactive hybrid Quadrupole-Orbitrap mass spectrometer (Thermo Fisher Scientific) in electrospray ionization (ESI) positive-ion mode. The MS/MS spectra were acquired in DDA mode, automatically alternating between a full scan of precursor ions (*m/z* 300–2000) in the Orbitrap (resolution 70,000; automatic gain control target 3,000,000; maximum injection time 60 ms) and subsequent HCD MS/MS scans of the top 10 most abundant precursor ions (*m/z* 200–2000) in the Orbitrap (resolution 17,500; automatic gain control target 500,000; maximum injection time 100 ms; isolation window 4.0 *m/z*; normalized collision energy 30%).

### Proteomic data processing

We processed all raw MS/MS files, including those analyzed previously and publicly available proteome datasets of human cell lines (SH-SY5Y and A549; two biological replicates for each cell line), deposited in ProteomeXchange (dataset identifier PXD004452 (*14*)). These files were converted into MGF format using Thermo Proteome Discoverer 2.2.0.388 (Thermo Fisher Scientific). The public proteomic data files underwent precursor mass recalibration before being converted into MGF files with Thermo Proteome Discoverer. The MGF files were then submitted to an in-house Mascot Server 2.7.0 (Matrix Science, London, UK) for PSM searches via the Mascot algorithm, employing Thermo Proteome Discoverer with both target and decoy sequence databases. The search parameters included variable modifications (acetyl [protein N-term], oxidation [M], and Gln to pyro-Glu conversion [N-term Q]), static modification (carbamidomethyl [C]), a maximum of three missed cleavages by trypsin digestion, a precursor mass tolerance of 15 ppm, and a fragment mass tolerance of 30 mmu.

Separate PSM searches for target decoy-based FDR estimation were conducted using the following target sequence databases and their corresponding decoy databases (created by reversing the target sequences with Thermo Proteome Discoverer): for proteogenomic data mining, two different target databases were used in parallel (combination of human RefSeq Proteins and the custom SEP or LEP sequence database for SH-SY5Y or A549, respectively); for detecting *in vitro* synthesized proteins, the target database consisted of concatenated peptides (artificial proteins) sequences and the whole protein sequences of *Escherichia coli* BL21 (DE3) (NCBI GenBank accession CP001509; version CP001509.3 (*62*)). The search results were filtered using the Percolator algorithm to maintain an estimated target decoy-based FDR of 1% and 0.1% at the peptide and protein levels, respectively. Identified peptides/proteins were further searched against the UniProt Knowledgebase. Manual inspection of MS/MS spectra was conducted with Thermo Proteome Discoverer and Thermo Xcalibur Qual Browser 3.1.66.10 (Thermo Fisher Scientific). The results were visualized using the integrative proteomics data viewer PDV 1.8.1 (*63*). Mapping of DNA sequences corresponding to peptides/proteins of interest onto the human genome was performed using the Integrative Genomics Viewer (IGV) 2.8.0.

### Protein expression in human cells

Human embryonic kidney cells (HEK293T) were grown at 37°C in 5% CO_2_ to 70–90% confluence in Dulbecco’s Modified Eagle Medium supplemented with 10% heat-inactivated fetal bovine serum. DNA was transfected in 96-well tissue culture plates using Lipofectamine 3000 (Thermo Fisher Scientific) according to the manufacturer’s instructions. The cell cultures were then incubated for an additional 2 days and utilized to measure the fluorescence intensity of the reporter proteins AmCyan1 and DsRed-Express at respective excitation/emission wavelengths of 458/489 nm and 554/586 nm, using a Varioskan LUX Multimode Microplate Reader (Thermo Fisher Scientific).

## Supporting information

Supplementary materials (DataS1)

Supplementary materials (DataS2)

## Acknowledgments

The authors thank Dr. Makoto Muroi (RIKEN), Dr. Makoto Kawatani (RIKEN), Dr. Takehiro Suzuki (RIKEN), Ms. Emiko Sanada (RIKEN), Ms. Yumi Sato (RIKEN), and Ms. Tamayo Oishi (RIKEN) for their technical support.

## Author contributions

Y.N. and N.D. conceived the research; Y.N. conducted the experiments; Y.N. and N.D. analyzed the results; Y.N. wrote the paper.

## Data and materials availability

All data supporting the findings of this study are available within the paper and its supplementary materials.

