## Supplementary materials (Figs and Tables) for "Discovery of evolutionarily extended *cis*-regulatory overlapping genes expanding the protein universe from animals to humans"

**This PDF file includes:**

Figs. S1 to S20
Tables S1 to S4
Data S1 to S2

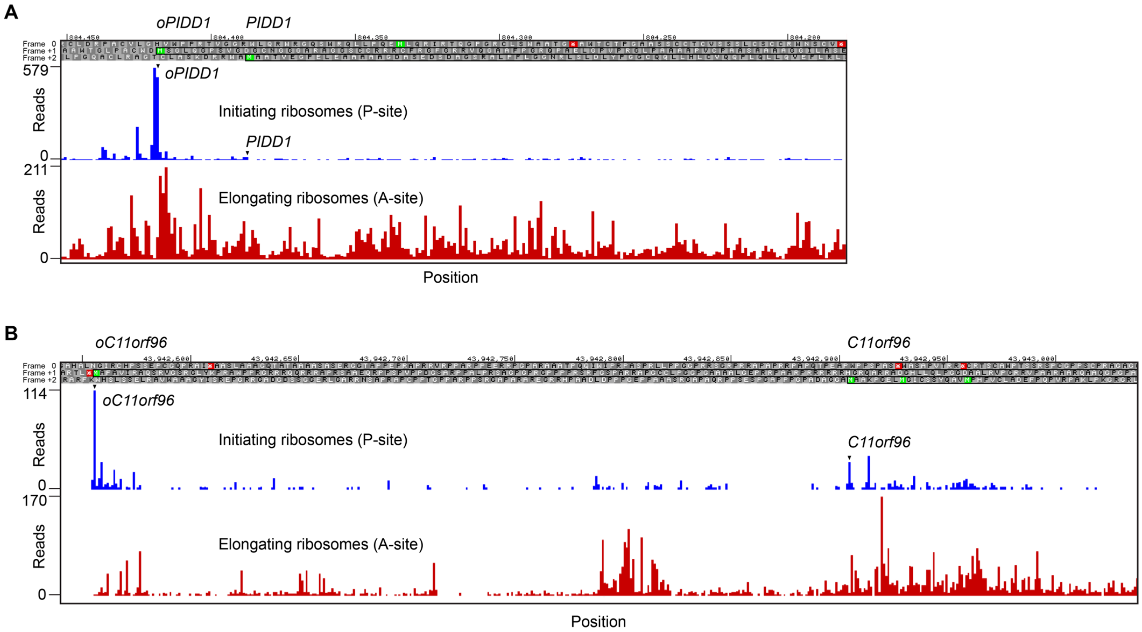


**Fig. S1. Survey of public Ribo-Seq data on *oPIDD1* and *oC11orf96*.** Aggregated profiles of initiating and elongating ribosomes at *oPIDD1* (**A**) and *oC11orf96* (**B**) were obtained from GWIPS-viz (https://gwips.ucc.ie).

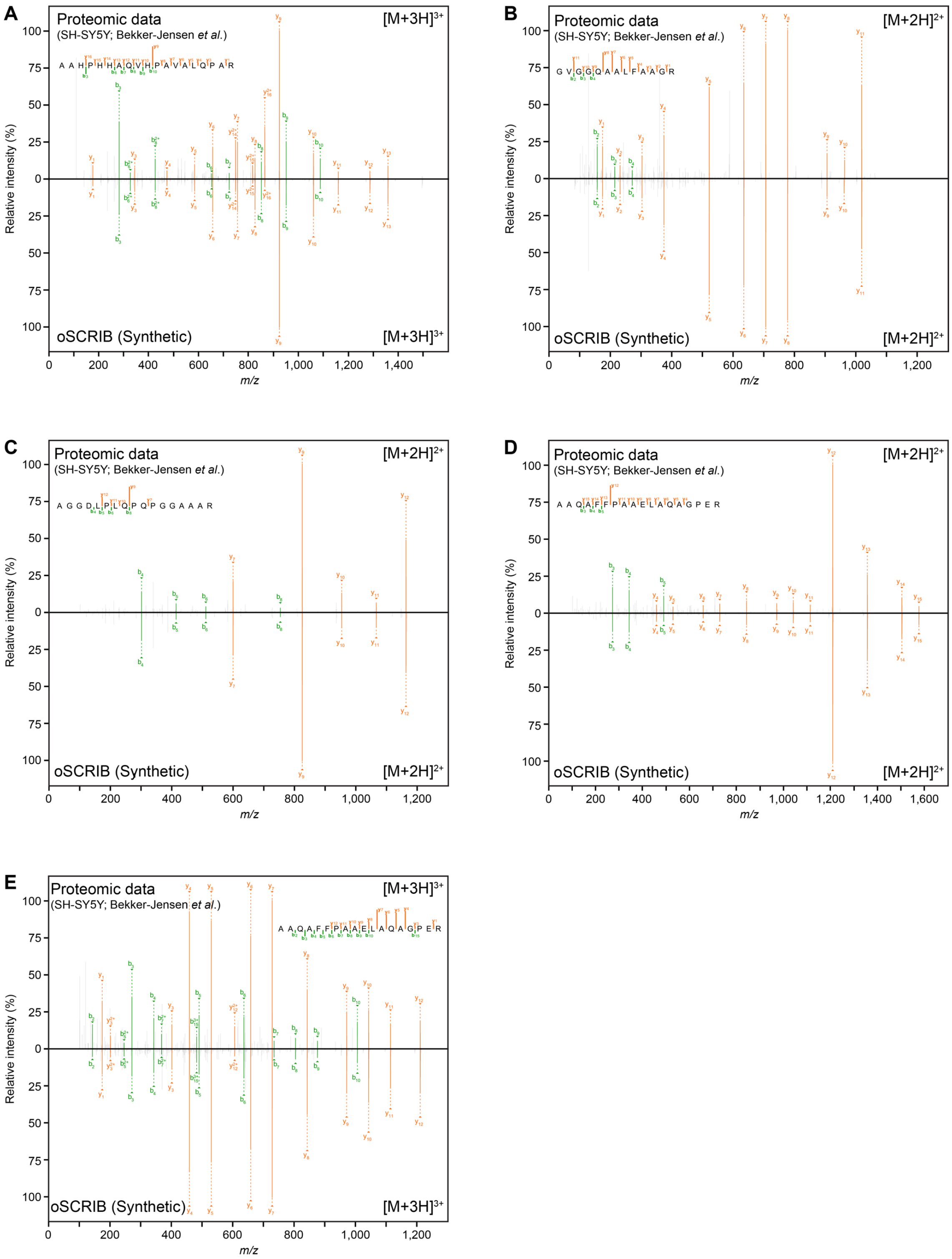


**Fig. S2. Mirror plots of MS/MS spectra from oSCRIB.**

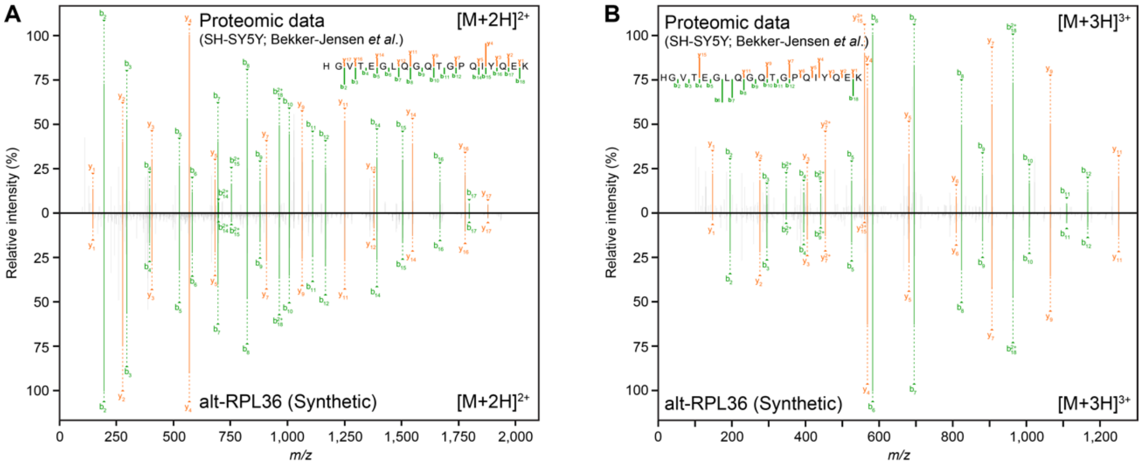


**Fig. S3. Mirror plots of MS/MS spectra from alt-RPL36.**

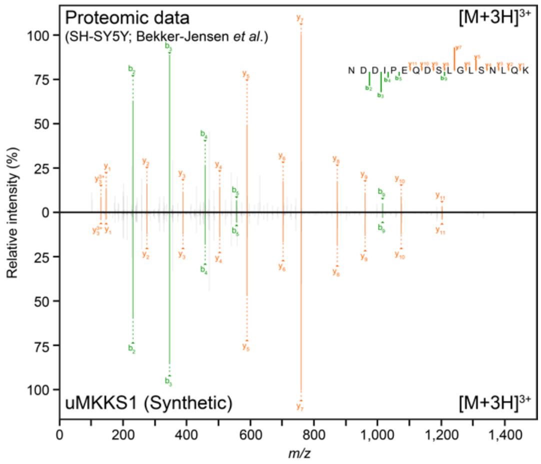


**Fig. S4. Mirror plots of MS/MS spectra from uMKKS1.**

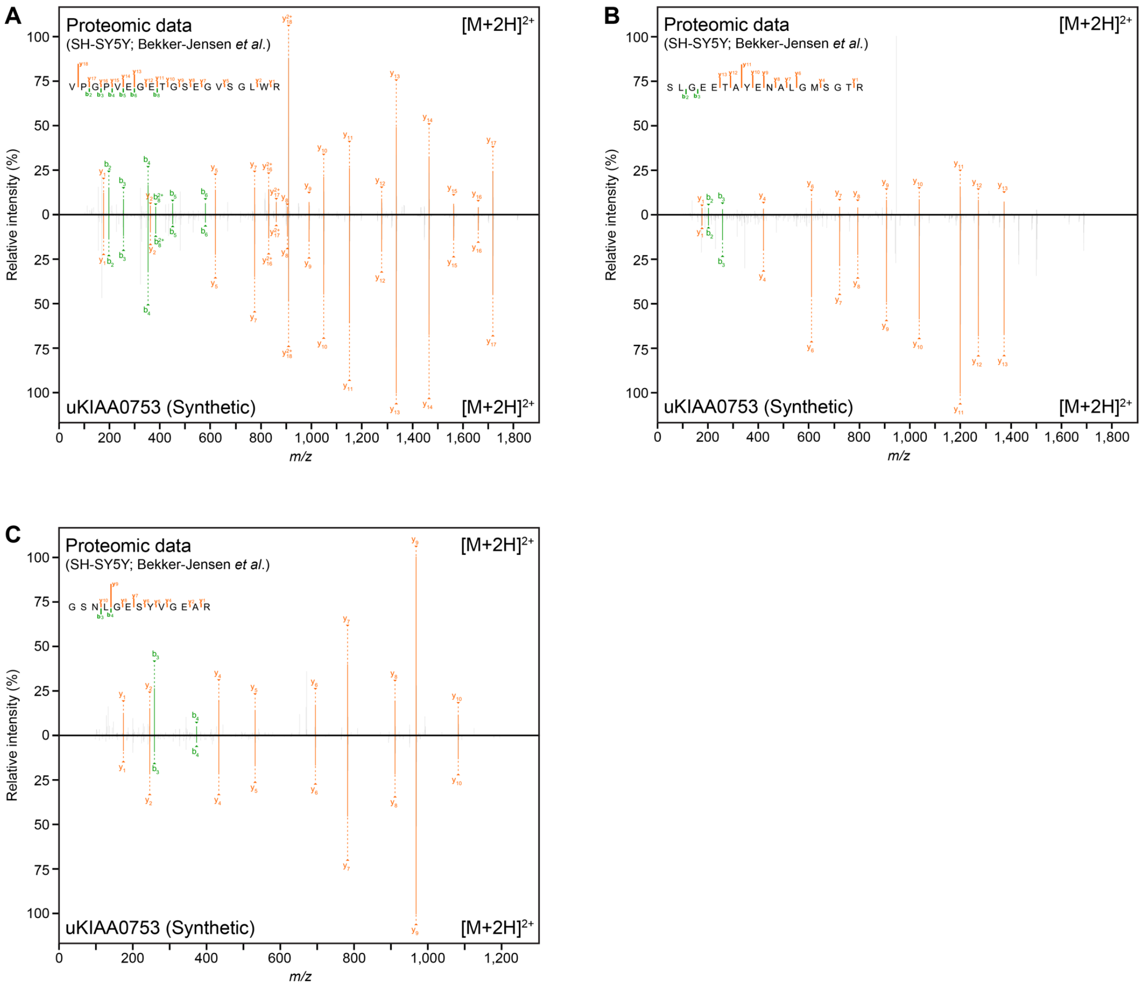


**Fig. S5. Mirror plots of MS/MS spectra from uKIAA0753.**

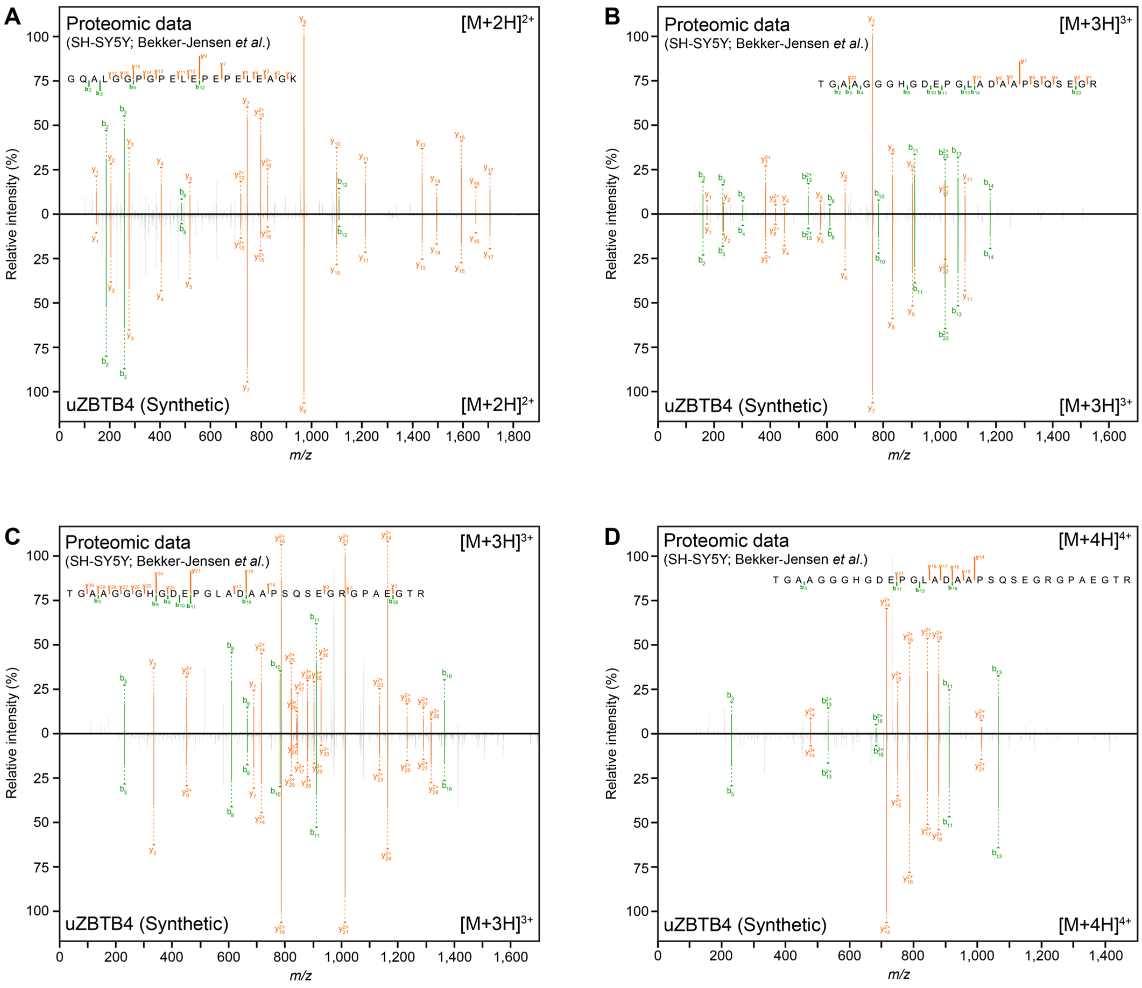


**Fig. S6. Mirror plots of MS/MS spectra from uZBTB4.**

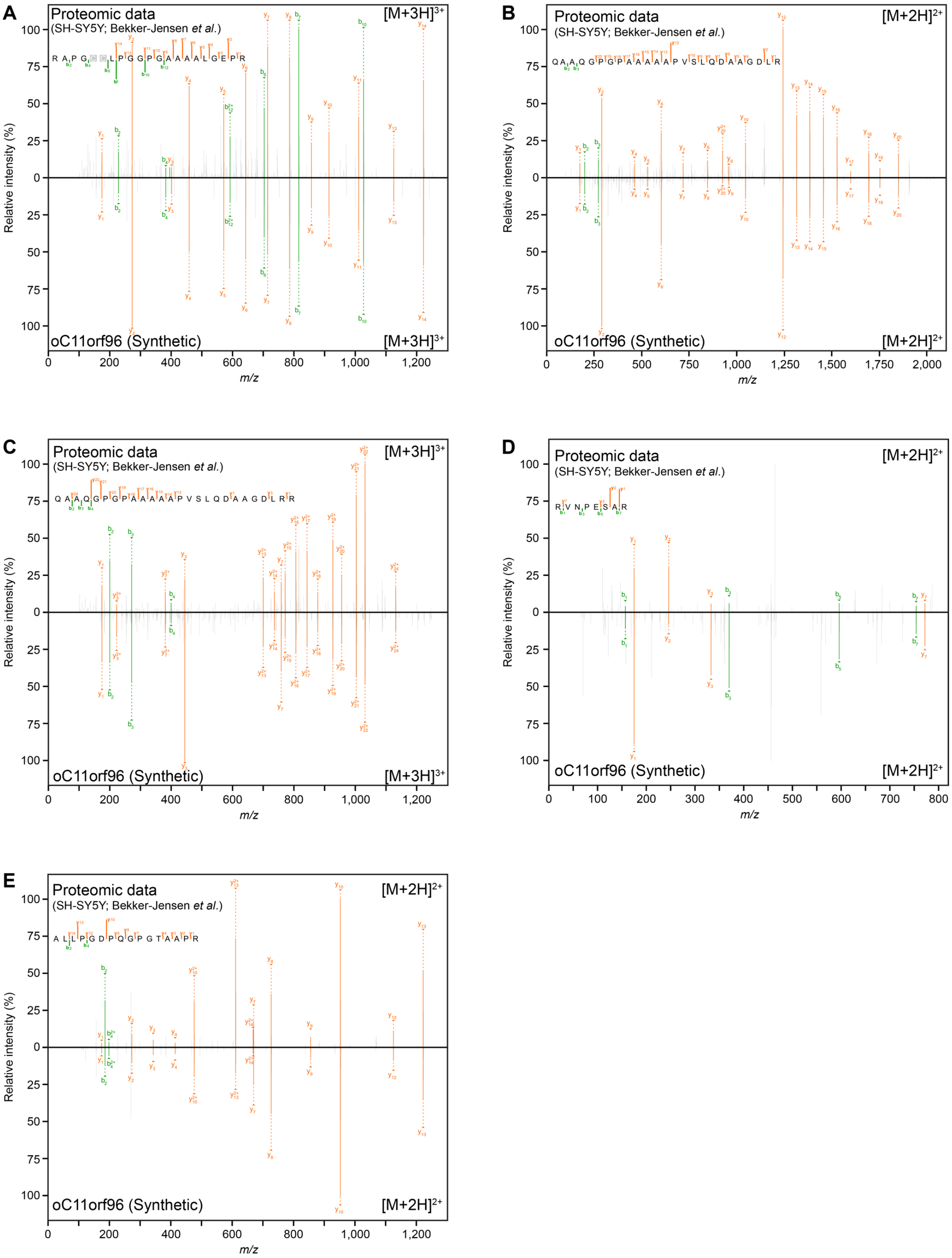


**Fig. S7. Mirror plots of MS/MS spectra from oC11orf96.**

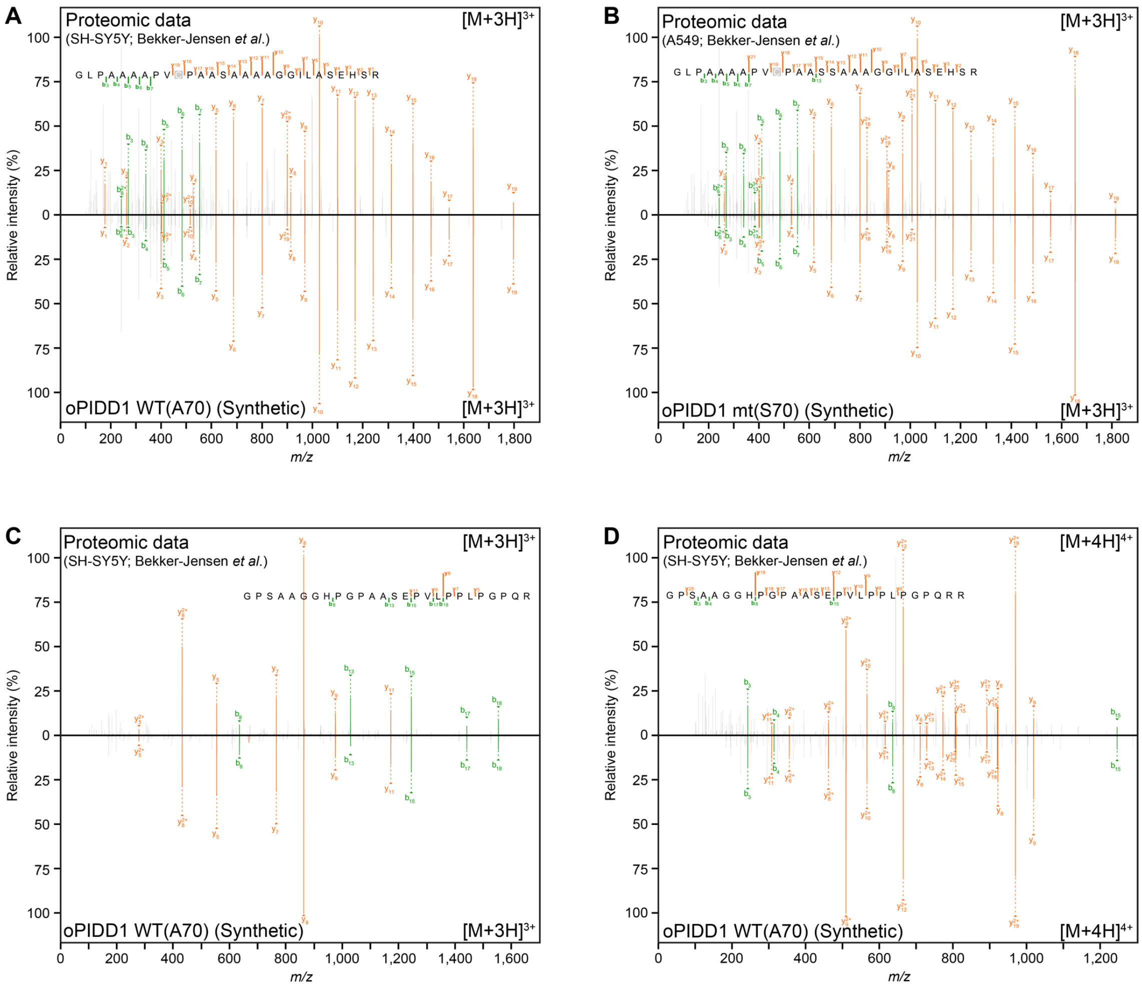


**Fig. S8. Mirror plots of MS/MS spectra from oPIDD1.**

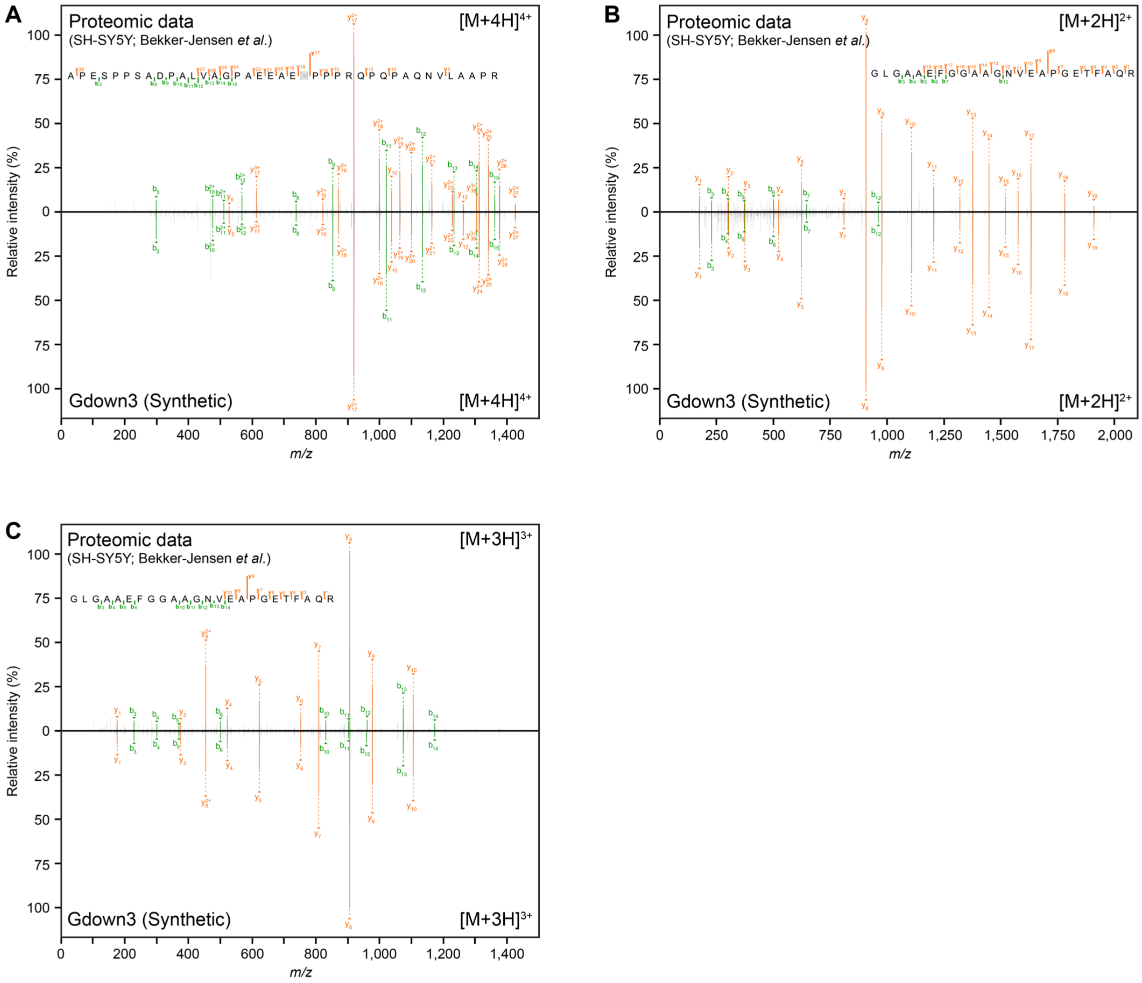


**Fig. S9. Mirror plots of MS/MS spectra from Gdown3.**

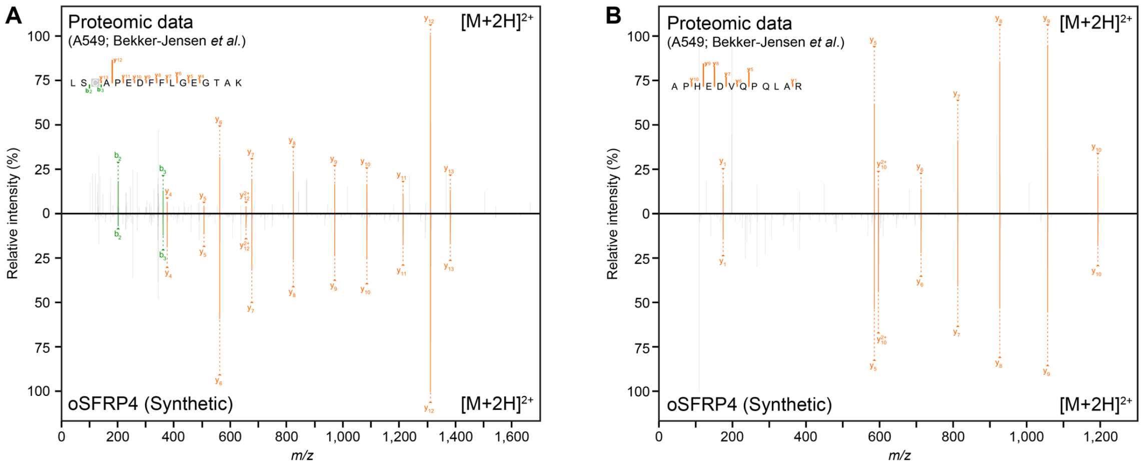


**Fig. S10. Mirror plots of MS/MS spectra from oSFRP4.**

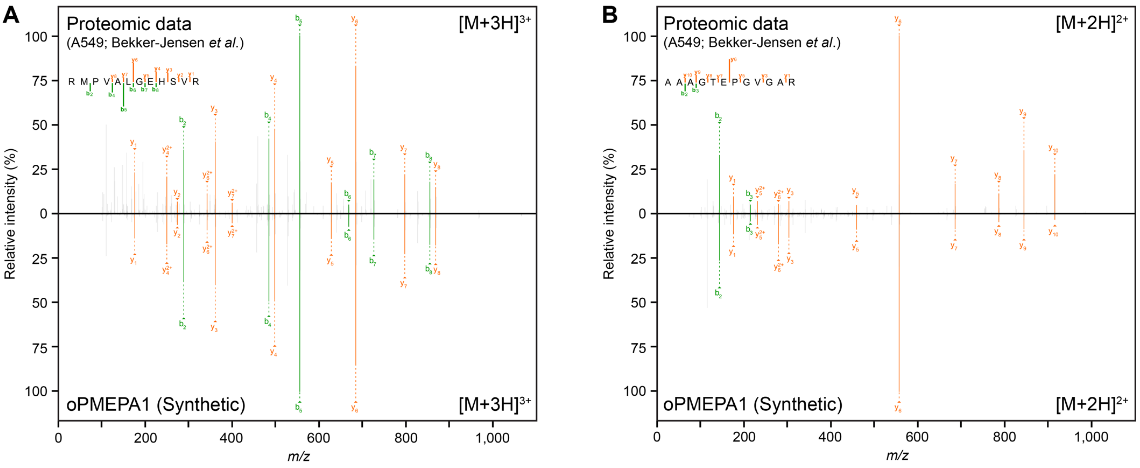


**Fig. S11. Mirror plots of MS/MS spectra from oPMEPA1.**

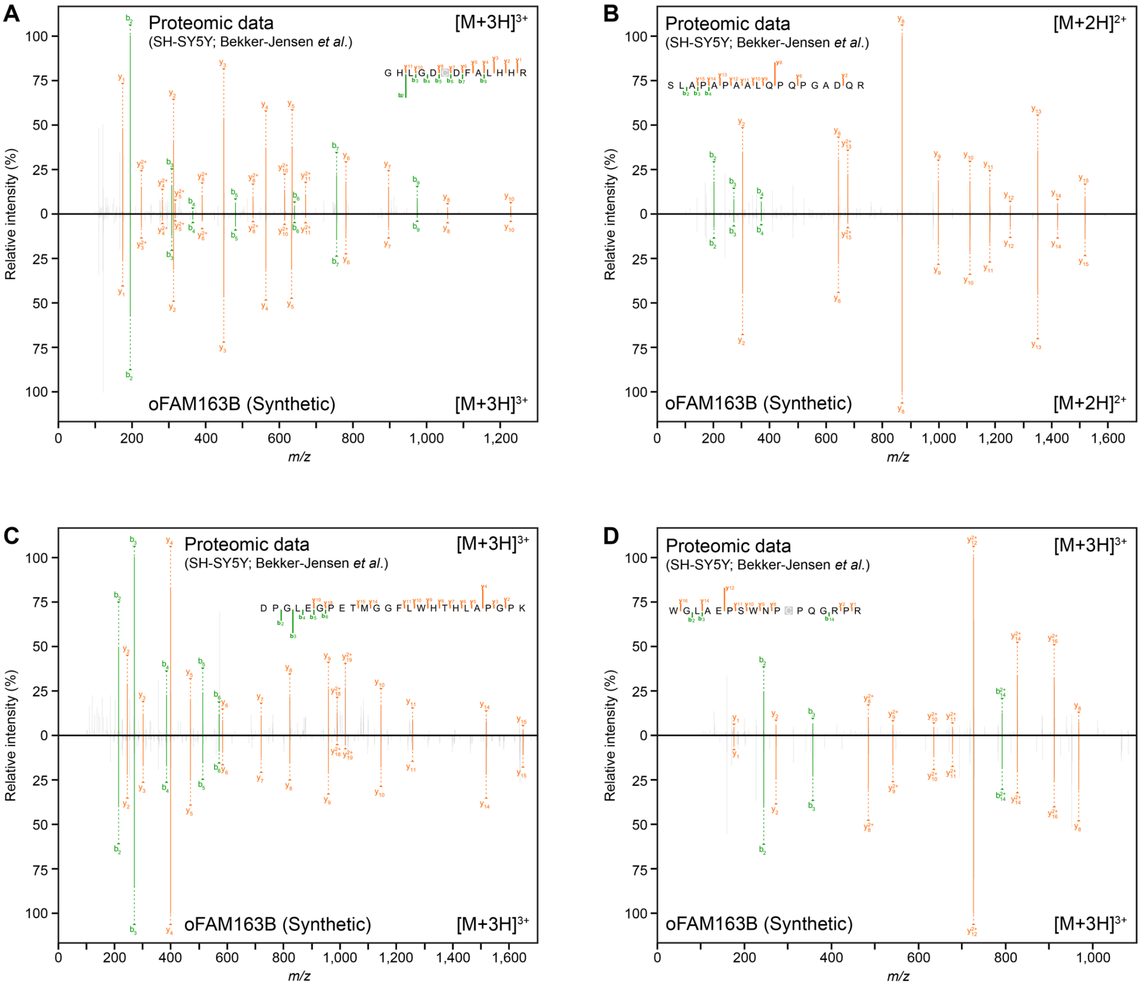


**Fig. S12. Mirror plots of MS/MS spectra from oFAM163B.**

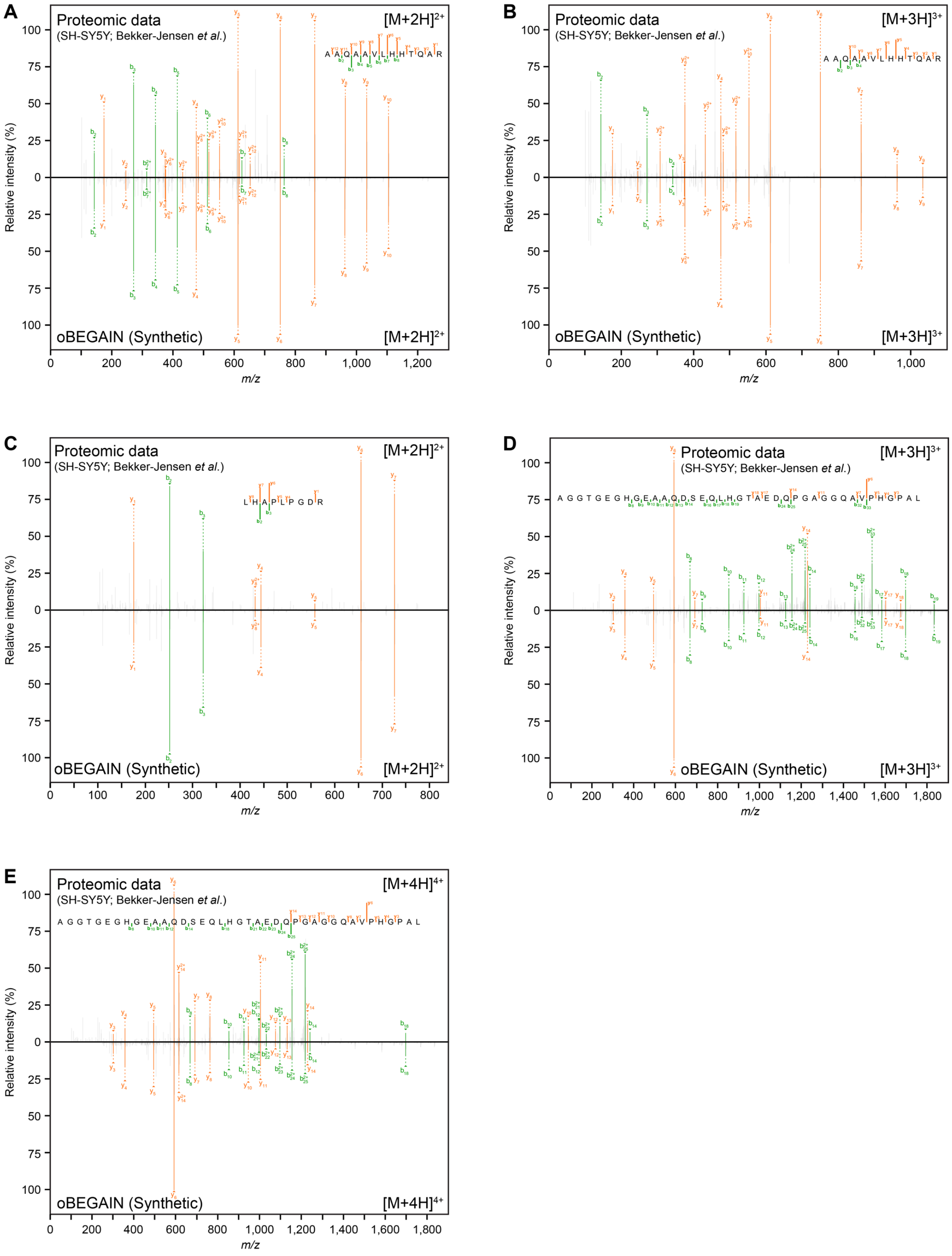


**Fig. S13. Mirror plots of MS/MS spectra from oBEGAIN.**

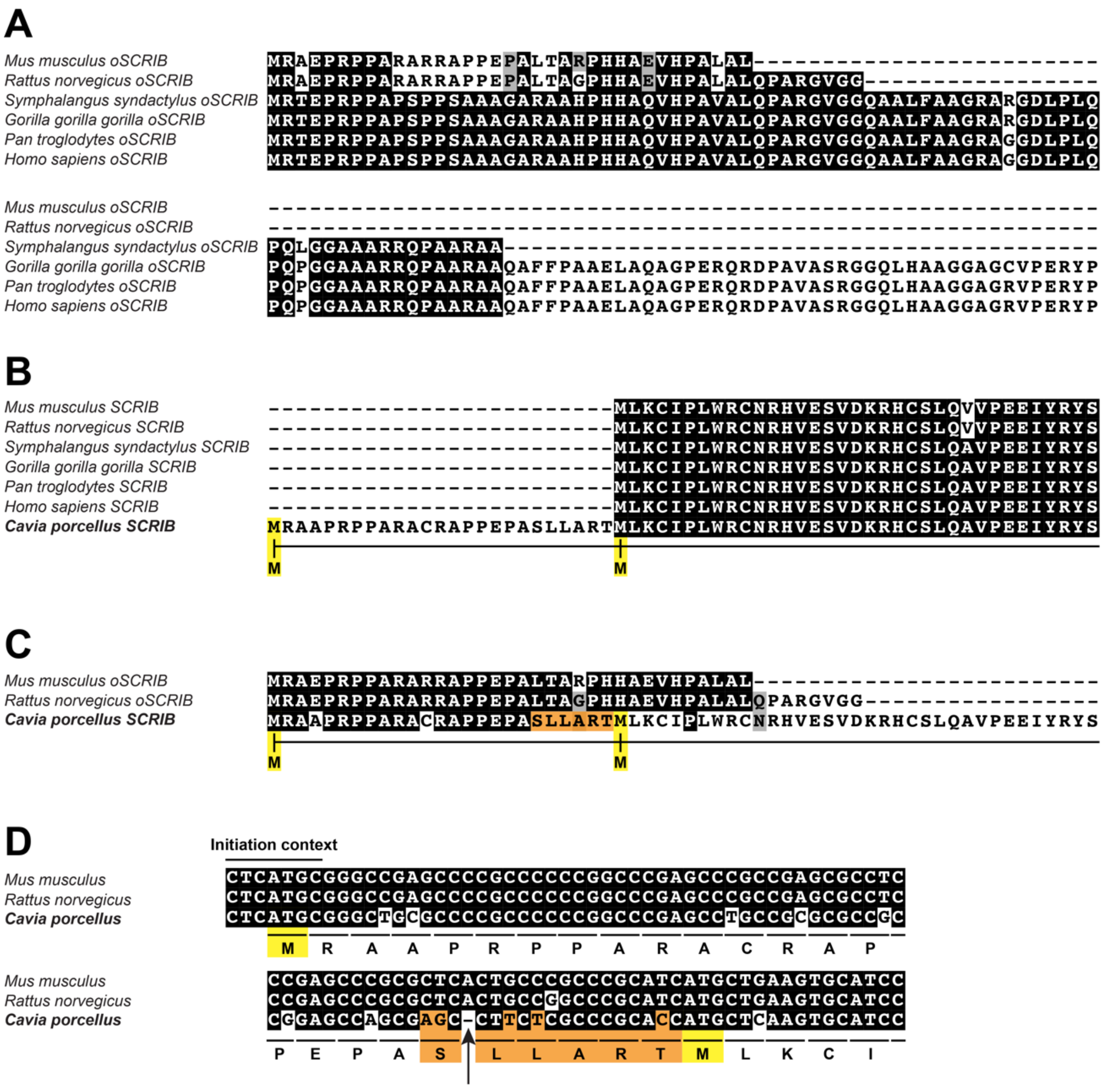


**Fig. S14. Sequence analyses of oSCRIB and SCRIB.** (**A, B**) Translation products of *oSCRIB* (**A**) and *SCRIB* (**B**) in animals. (**C, D**) Comparison of sequences of *Mus musculus*/*Rattus norvegicus* oSCRIB and *Cavia porcellus* SCRIB at the protein (**C**) and transcript (**D**) levels. The possible frameshift site is indicated by an arrow. Protein sequences were aligned and colored using GenomeNet ClustalW 2.1 (https://www.genome.jp/tools-bin/clustalw) and pyBoxshade (https://github.com/mdbaron42/pyBoxshade).

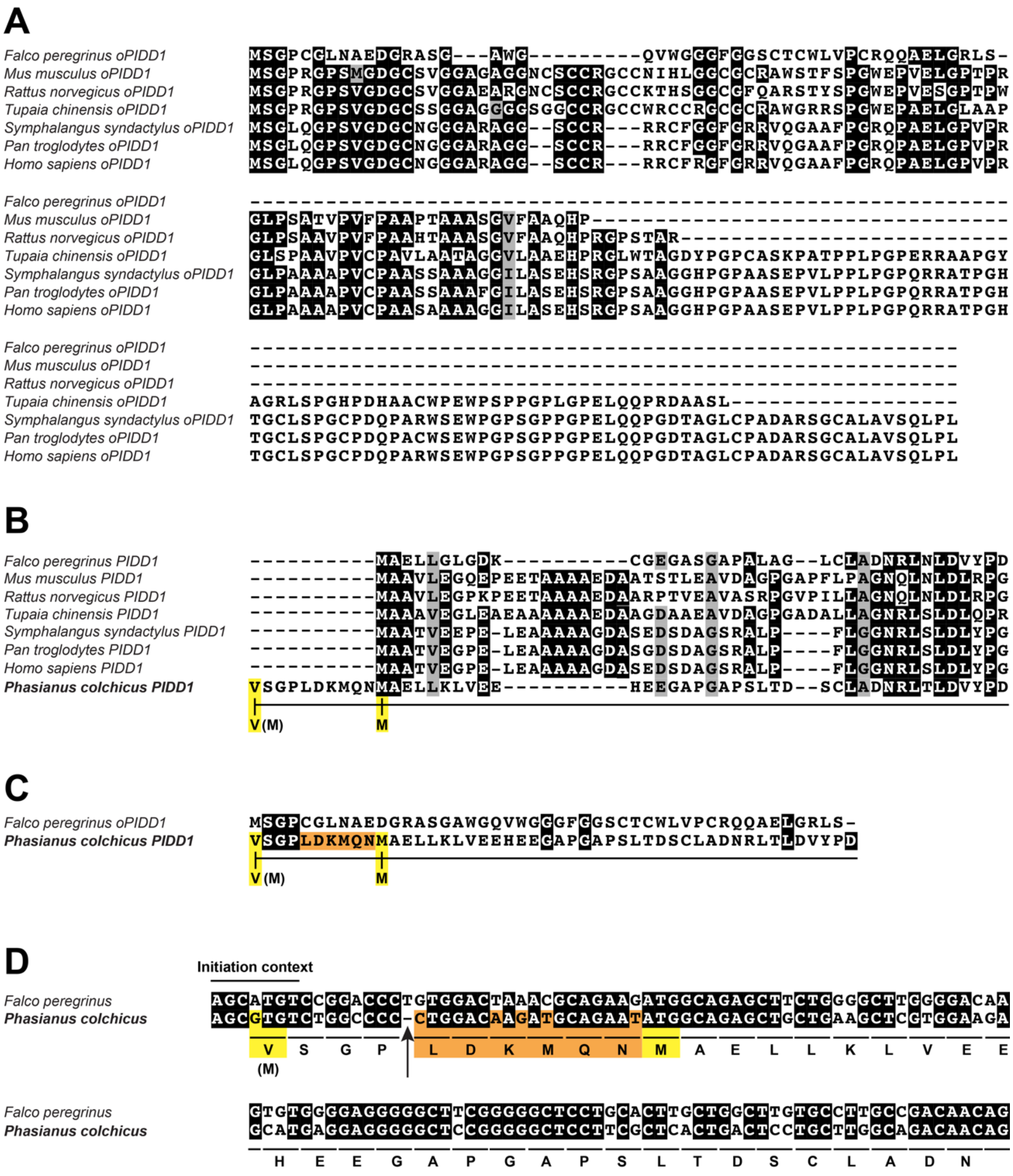

**Fig. S15. Sequence analyses of oPIDD1 and PIDD1.** (**A, B**) Translation products of *oPIDD1* (**A**) and *PIDD1* (**B**) in animals. (**C, D**) Comparison of the sequences of *Falco peregrinus* oPIDD1 and *Phasianus colchicus* PIDD1 at the protein (**C**) and transcript (**D**) levels. The possible frameshift site is indicated by an arrow. The exchange between an optimal ATG start codon and a suboptimal GTG start codon is also indicated. Protein sequences were aligned and colored using GenomeNet ClustalW 2.1 (https://www.genome.jp/tools-bin/clustalw) and pyBoxshade (https://github.com/mdbaron42/pyBoxshade).

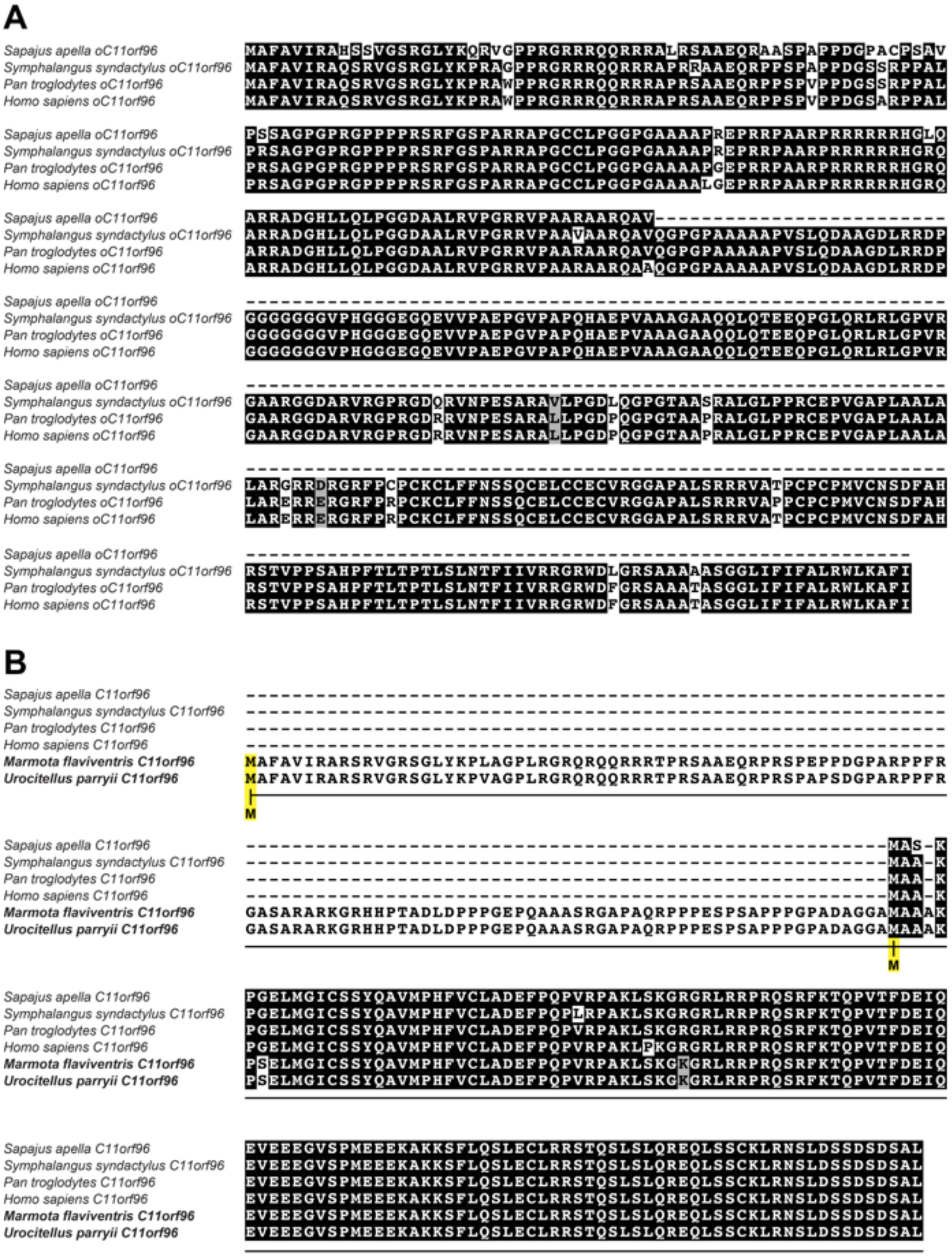

A
Sapajus apella oC11orf96 MAFAVIRAHSSVGSRGLYKQRVGPPRGRRRQQRRRALRSAAEQRAASPAPPDGPACPSAV
Symphalangus syndactylus oC11orf96 MAFAVIRAQSRVGSRGLYKPRAGPPRGRRRQQRRRAPRRAAEQRPPSPAPPDGSSRPPAL
Pan troglodytes oC11orf96 MAFAVIRAQSRVGSRGLYKPRAWPPRGRRRQQRRRAPRSAAEQRPPSPVPPDGSSRPPAL
Homo sapiens oC11orf96 MAFAVIRAQSRVGSRGLYKPRAWPPRGRRRQQRRRAPRSAAEQRPPSPVPPDGSARPPAL
Sapajus apella oC11orf96 PSSAGPGPRGPPPPRSRFGSPARRAPGCCLPGGPGAAAAPREPRRPAARPRRRRRRHGLQ
Symphalangus syndactylus oC11orf96 PRSAGPGPRGPPPPRSRFGSPARRAPGCCLPGGPGAAAAPREPRRPAARPRRRRRRHGRQ
Pan troglodytes oC11orf96 PRSAGPGPRGPPPPRSRFGSPARRAPGCCLPGGPGAAAAPGEPRRPAARPRRRRRRHGRQ
Homo sapiens oC11orf96 PRSAGPGPRGPPPPRSRFGSPARRAPGCCLPGGPGAAAALGEPRRPAARPRRRRRRHGRQ
Sapajus apella oC11orf96 ARRADGHLLQLPGGDAALRVPGRRVPAARAARQAV-------------------------
Symphalangus syndactylus oC11orf96 ARRADGHLLQLPGGDAALRVPGRRVPAAVAARQAVQGPGPAAAAAPVSLQDAAGDLRRDP
Pan troglodytes oC11orf96 ARRADGHLLQLPGGDAALRVPGRRVPAARAARQAVQGPGPAAAAAPVSLQDAAGDLRRDP
Homo sapiens oC11orf96 ARRADGHLLQLPGGDAALRVPGRRVPAARAARQAAQGPGPAAAAAPVSLQDAAGDLRRDP
Sapajus apella oC11orf96 ------------------------------------------------------------
Symphalangus syndactylus oC11orf96 GGGGGGGVPHGGGEGQEVVPAEPGVPAPQHAEPVAAAGAAQQLQTEEQPGLQRLRLGPVR
Pan troglodytes oC11orf96 GGGGGGGVPHGGGEGQEVVPAEPGVPAPQHAEPVAAAGAAQQLQTEEQPGLQRLRLGPVR
Homo sapiens oC11orf96 GGGGGGGVPHGGGEGQEVVPAEPGVPAPQHAEPVAAAGAAQQLQTEEQPGLQRLRLGPVR
Sapajus apella oC11orf96 ------------------------------------------------------------
Symphalangus syndactylus oC11orf96 GAARGGDARVRGPRGDQRVNPESARAVLPGDLQGPGTAASRALGLPPRCEPVGAPLAALA
Pan troglodytes oC11orf96 GAARGGDARVRGPRGDRRVNPESARALLPGDPQGPGTAAPRALGLPPRCEPVGAPLAALA
Homo sapiens oC11orf96 GAARGGDARVRGPRGDRRVNPESARALLPGDPQGPGTAAPRALGLPPRCEPVGAPLAALA
Sapajus apella oC11orf96 ------------------------------------------------------------
Symphalangus syndactylus oC11orf96 LARGRRDRGRFPCPCKCLFFNSSQCELCCECVRGGAPALSRRRVATPCPCPMVCNSDFAH
Pan troglodytes oC11orf96 LARERREGRFPRPCKCLFFNSSQCELCCECVRGGAPALSRRRVAPPCPCPMVCNSDFAH
Homo sapiens oC11orf96 LARERREGRFPRPCKCLFFNSSQCELCCECVRGGAPALSRRRVATPCPCPMVCNSDFAH
Sapajus apella oC11orf96 ---------------------------------------------------------
Symphalangus syndactylus oC11orf96 RSTVPPSAHPFTLTPTLSLNTFIIVRRGRWDLGRSAAAAASGGLIFIFALRWLKAFI
Pan troglodytes oC11orf96 RSTVPPSAHPFTLTPTLSLNTFIIVRRGRWDFGRSAAATASGGLIFIFALRWLKAFI
Homo sapiens oC11orf96 RSTVPPSAHPFTLTPTLSLNTFIIVRRGRWDFGRSAAATASGGLIFIFALRWLKAFI
B
Sapajus apella C11orf96 ------------------------------------------------------------
Symphalangus syndactylus C11orf96 ------------------------------------------------------------
Pan troglodytes C11orf96 ------------------------------------------------------------
Homo sapiens C11orf96 ------------------------------------------------------------
Marmota flaviventris C11orf96 MAFAVIRARSRVGRSGLYKPLAGPLRGRQRQQRRRTPRSAAEQRPRSPEPPDGPARPPFR
Urocitellus parryii C11orf96 MAFAVIRARSRVGRSGLYKPVAGPLRGRQRQQRRRTPRSAAEQRPRSPAPSDGPARPPFR
M
Sapajus apella C11orf96 -------------------------------------------------------MAS-K
Symphalangus syndactylus C11orf96 -------------------------------------------------------MAA-K
Pan troglodytes C11orf96 -------------------------------------------------------MAA-K
Homo sapiens C11orf96 -------------------------------------------------------MAA-K
Marmota flaviventris C11orf96 GASARARKGRHHPTADLDPPPGEPQAAASRGAPAQRPPPESPSAPPPGPADAGGAMAAAK
Urocitellus parryii C11orf96 GASARARKGRHHPTADLDPPPGEPQAAASRGAPAQRPPPESPSAPPPGPADAGGAMAAAK
M
Sapajus apella C11orf96 PGELMGICSSYQAVMPHFVCLADEFPQPVRPAKLSKGRGRLRRPRQSRFKTQPVTFDEIQ
Symphalangus syndactylus C11orf96 PGELMGICSSYQAVMPHFVCLADEFPQPLRPAKLSKGRGRLRRPRQSRFKTQPVTFDEIQ
Pan troglodytes C11orf96 PGELMGICSSYQAVMPHFVCLADEFPQPVRPAKLSKGRGRLRRPRQSRFKTQPVTFDEIQ
Homo sapiens C11orf96 PGELMGICSSYQAVMPHFVCLADEFPQPVRPAKLPKGRGRLRRPRQSRFKTQPVTFDEIQ
Marmota flaviventris C11orf96 PSELMGICSSYQAVMPHFVCLADEFPQPVRPAKLSKGKGRLRRPRQSRFKTQPVTFDEIQ
Urocitellus parryii C11orf96 PSELMGICSSYQAVMPHFVCLADEFPQPVRPAKLSKGKGRLRRPRQSRFKTQPVTFDEIQ
Sapajus apella C11orf96 EVEEEGVSPMEEEKAKKSFLQSLECLRRSTQSLSLQREQLSSCKLRNSLDSSDSDSAL
Symphalangus syndactylus C11orf96 EVEEEGVSPMEEEKAKKSFLQSLECLRRSTQSLSLQREQLSSCKLRNSLDSSDSDSAL
Pan troglodytes C11orf96 EVEEEGVSPMEEEKAKKSFLQSLECLRRSTQSLSLQREQLSSCKLRNSLDSSDSDSAL
Homo sapiens C11orf96 EVEEEGVSPMEEEKAKKSFLQSLECLRRSTQSLSLQREQLSSCKLRNSLDSSDSDSAL
Marmota flaviventris C11orf96 EVEEEGVSPMEEEKAKKSFLQSLECLRRSTQSLSLQREQLSSCKLRNSLDSSDSDSAL
Urocitellus parryii C11orf96 EVEEEGVSPMEEEKAKKSFLQSLECLRRSTQSLSLQREQLSSCKLRNSLDSSDSDSAL

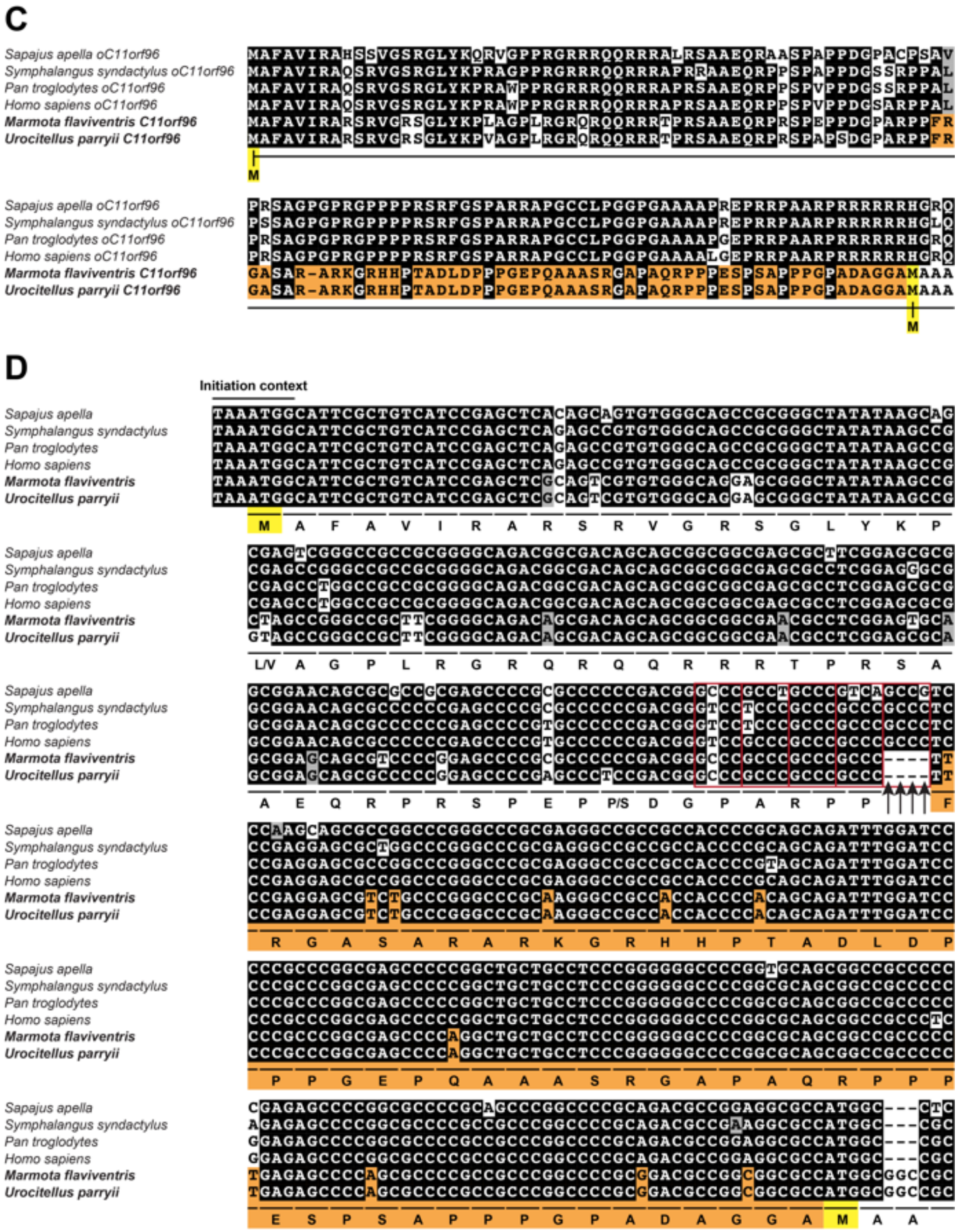

**Fig. S16. Sequence analyses of oC11orf96 and C11orf96.** (**A, B**) Translation products of *oC11orf96* (**A**) and *C11orf96* (**B**) in animals. (**C, D**) Comparison of sequences of *Sapajus apella*/*Symphalangus syndactylus*/*Pan troglodytes*/*Homo sapiens* oC11orf96 and *Marmota flaviventris*/*Urocitellus parryii* C11orf96 at the protein (**C**) and transcript (**D**) levels. The possible frameshift site is indicated by arrows. The red box outlines a tandem repeat of a GCCC nucleotide sequence. Protein sequences were aligned and colored using GenomeNet ClustalW 2.1 (https://www.genome.jp/tools-bin/clustalw) and pyBoxshade (https://github.com/mdbaron42/pyBoxshade).

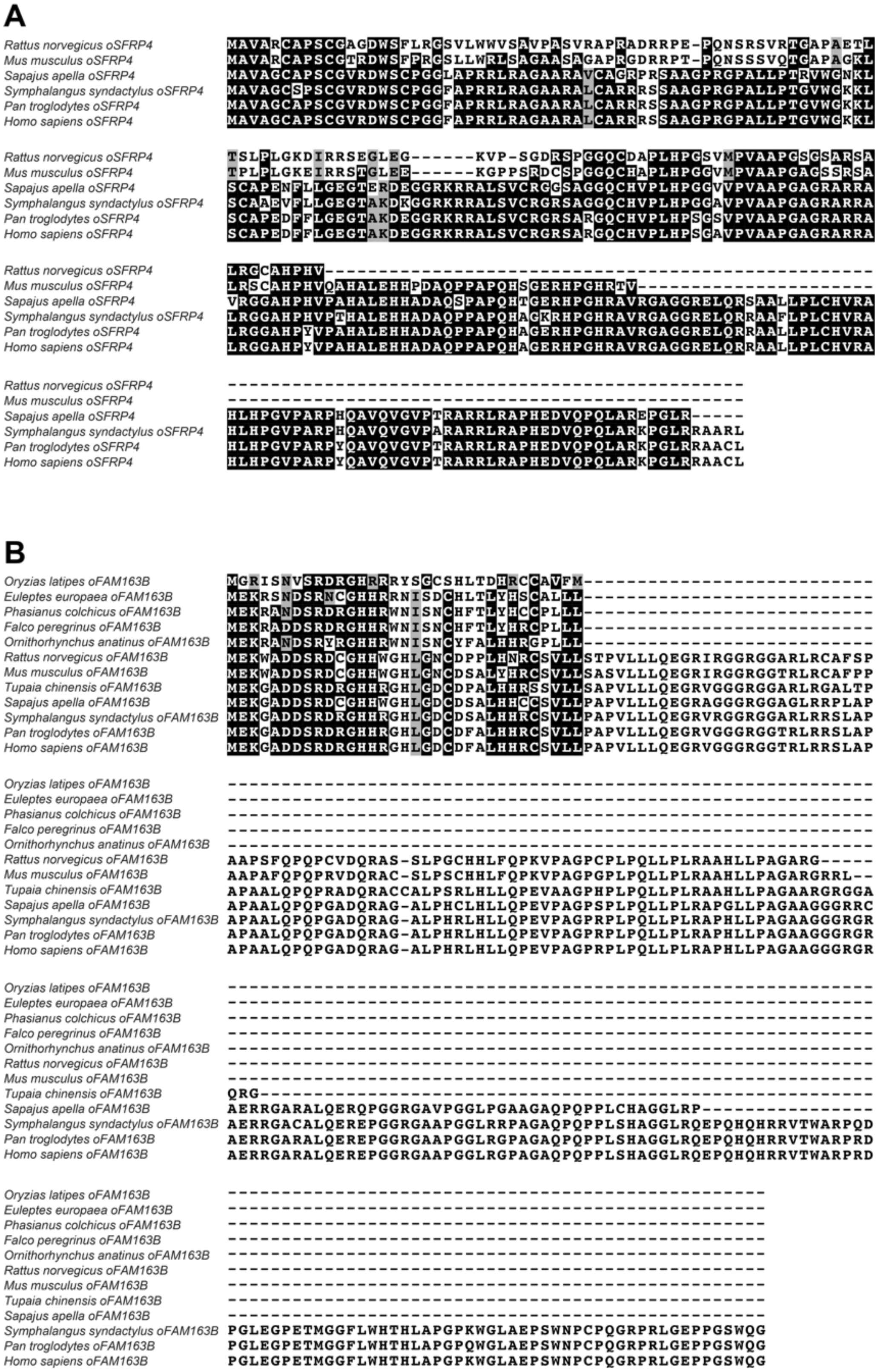
A
Rattus norvegicus oSFRP4
Mus musculus oSFRP4
Sapajus apella oSFRP4
Symphalangus syndactylus oSFRP4
Pan troglodytes oSFRP4
Homo sapiens oSFRP4
B
Oryzias latipes oFAM163B
Euleptes europaea oFAM163B
Phasianus colchicus oFAM163B
Falco peregrinus oFAM163B
Ornithorhynchus anatinus oFAM163B
Rattus norvegicus oFAM163B
Mus musculus oFAM163B
Tupaia chinensis oFAM163B
Sapajus apella oFAM163B
Symphalangus syndactylus oFAM163B
Pan troglodytes oFAM163B
Homo sapiens oFAM163B

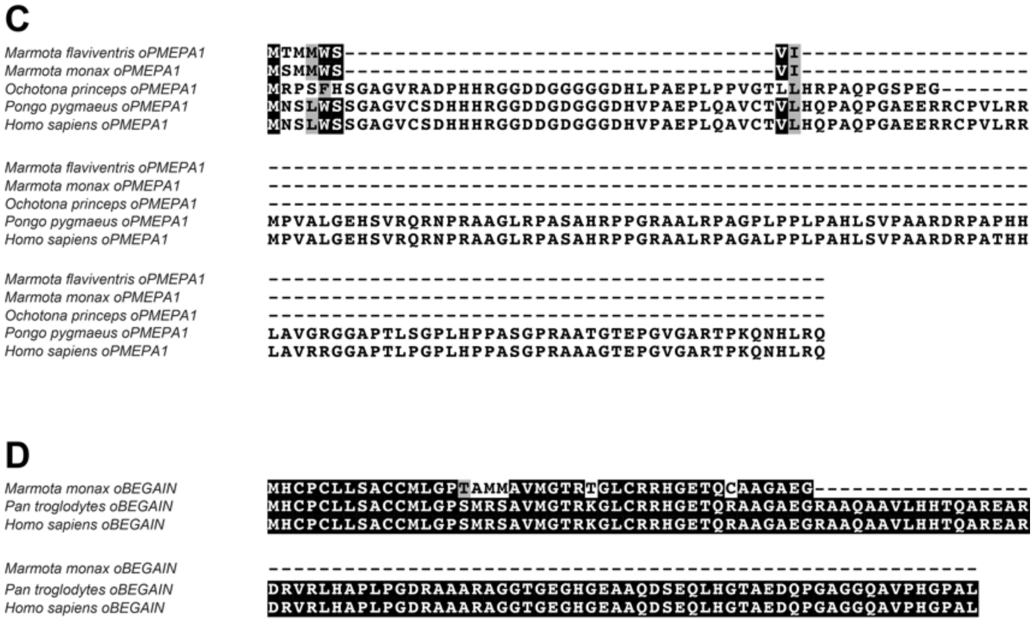


**Fig. S17. Sequence analyses of oSFRP4, oFAM163B, oPMEPA1, and oBEGAIN.** Translation products of *oSFRP4* (**A**), *oFAM163B* (**B**), *oPMEPA1* (**C**), and *oBEGAIN* (**D**) in animals.

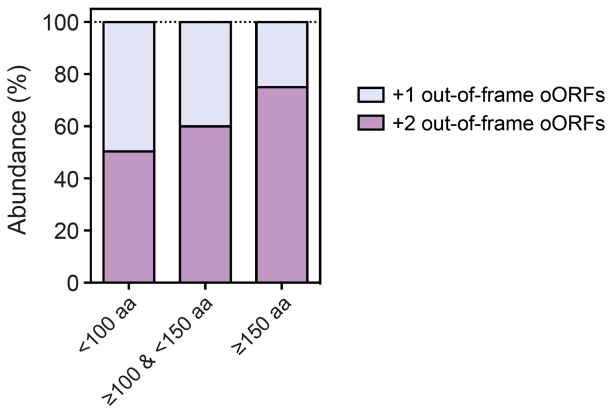


**Fig. S18. The ratio of +2 out-of-frame oORFs to +1 out-of-frame oORFs according to oORF length.** The results of Ribo-Seq-based translatomic analyses by Yang *et al.* were used to evaluate the ratio of +2 out-of-frame oORFs to +1 out-of-frame oORFs in humans. This value is based on the number of translatable oORFs that start with AUG and end with stop codons.

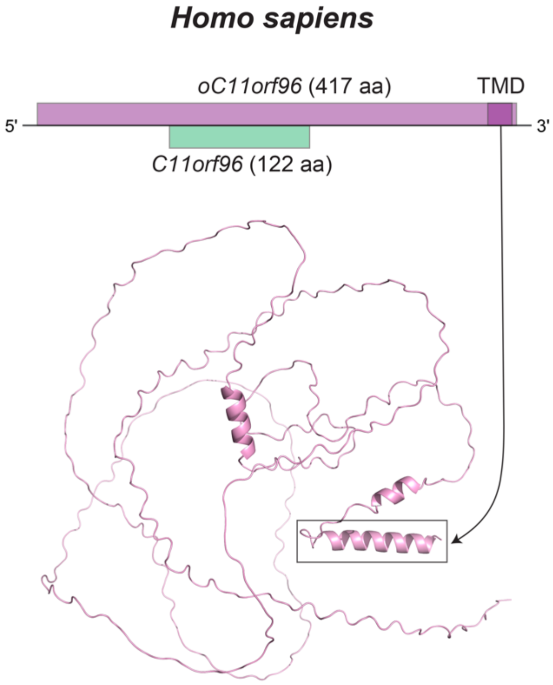


**Fig. S19. Three-dimensional structural modeling and functional domain analysis of oC11orf96 using AlphaFold2 and InterProScan.** TMD, putative transmembrane domain.

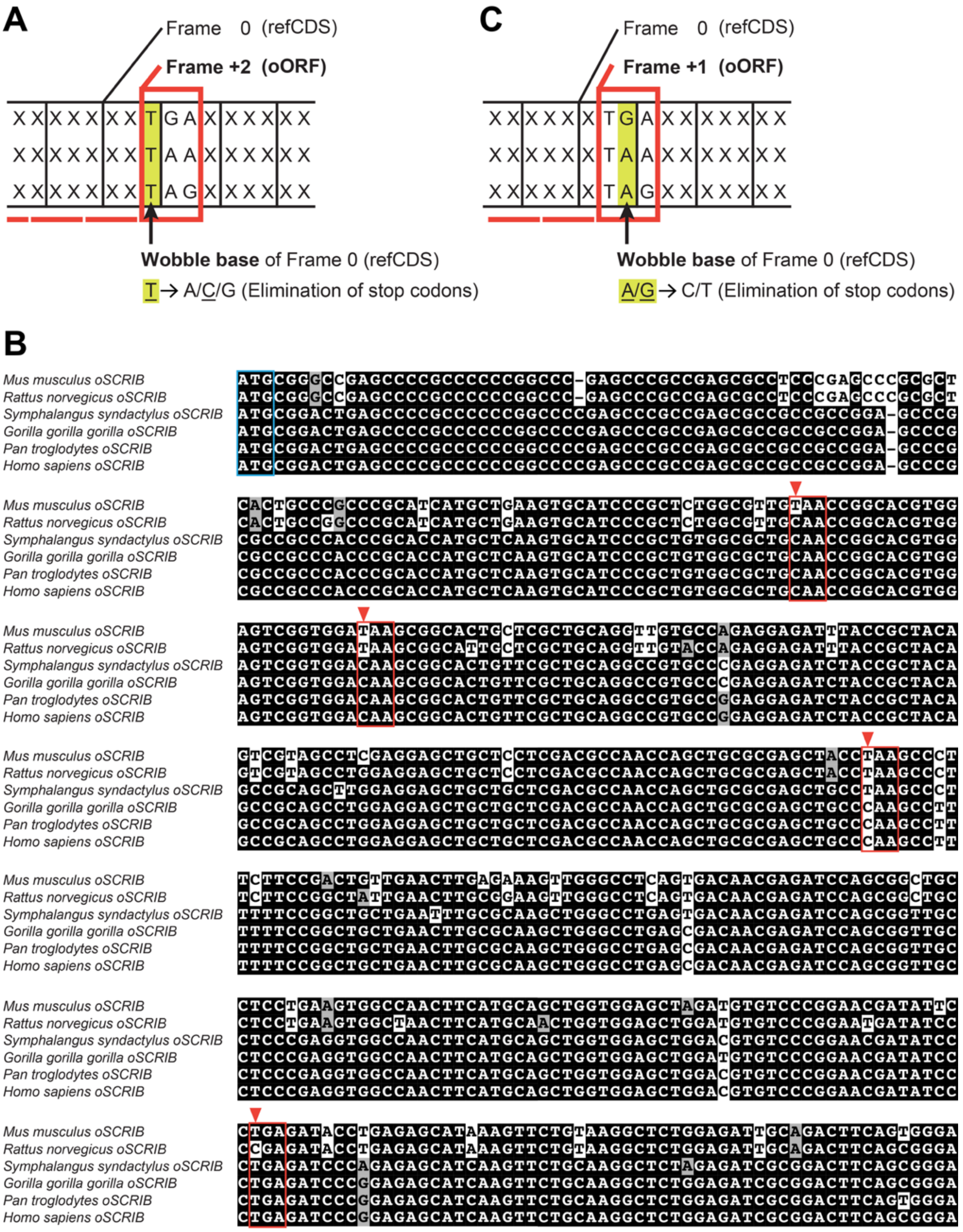


**Fig. S20. Advantage of +2 out-of-frame oORFs over +1 out-of-frame oORFs in the evolution by C-terminal extension.** (**A**) Mode of elimination of stop codons in +2 out-of-frame oORFs. (**B**) Representative example of elimination of stop codons in +2 out-of-frame oORFs. Thymine-to-cytosine (one-ring pyrimidines) transitions have occurred in stop codons of oSCRIB during mammalian evolution without changing genetic information in the refCDS (SCRIB). (**C**) Mode of elimination of stop codons in +1 out-of-frame oORFs.

**table S1**. Nucleotide and protein sequences of SEPs and LEPs in humans

| No | Gene name | Sequence (5' to 3')* |
| --- | --- | --- |
| 1 | *oSCRIB* | ATGCGGACTGAGCCCCGCCCCCCGGCCCCGAGCCCGCCGAGCGCCGCCGCCGGAGCCCGCGCCGCCCACCCGCACCATGCTCAAGTGCATCCCGCTGTGGCGCTGCAACCGGCACGTGGAGTCGGTGGACAAGCGGCACTGTTCGCTGCAGGCCGTGCCGGAGGAGATCTACCGCTACAGCCGCAGCCTGGAGGAGCTGCTGCTCGACGCCAACCAGCTGCGCGAGCTGCCCAAGCCTTTTTCCGGCTGCTGAACTTGCGCAAGCTGGGCCTGAGCGACAACGAGATCCAGCGGTTGCCTCCCGAGGTGGCCAACTTCATGCAGCTGGTGGAGCTGGACGTGTCCCGGAACGATATCCCTGA<br><br>Translation product:<br>MRTEPRPPAPSPPSAAAGARAAHPHHAQVHPAVALQPARGVGGQAALFAAGRAGGDLPLQPQPGGAAARRQPAARAAQAFFPAAELAQAGPERQRDPAVASRGGQLHAAGGAGRVPERYP |
| 2 | *alt-RPL36* | GTGGCGAGCGAGGCTGAAGGAGCCGGGACGCGGGGCTCTGGGCCTCGGGAACTGAGCCGGTACTCACCTCCGCCCCTTCTCCCCGTCGCTGTCCGCAGCCATGGCCCTACGCTACCCTATGGCCGTGGGCCTCAACAAGGGCCACAAAGTGACCAAGAACGTGAGCAAGCCCAGGCACAGCCGACGCCGCGGGCGTCTGACCAAACACACCAAGTTCGTGCGGGACATGATTCGGGAGGTGTGTGGCTTTGCCCCGTACGAGCGGCGCGCCATGGAGTTACTGAAGGTCTCCAAGGACAAACGGGCCCTCAAATTTATCAAGAAAAGGGTGGGGACGCACATCCGCGCCAAGAGGAAGCGGGAGGAGCTGAGCAACGTACTGGCCGCCATGAGGAAAGCTGCTGCCAAGAAAGACTGAGCCCCTCCCCTGCCCTCTCCCTGAAATAA<br><br>Translation product:<br>MASEAEGAGTRGSGPRELSRYSPPPLLPVAVRSHGPTLPYGRGPQQGPQSDQEREQAQAQPTPRASDQTHQVRAGHDSGGVWLCPVRAARHGVTEGLQGQTGPQIYQEKGGDAHPRQEEAGGAEQRTGRHEESCCQERLSPSPALSLK |
| 3 | *uMKKS1* | ATGAGCCTTCGGAACTTGTGGAGAGACTACAAAGTTTTGGTTGTTATGGTCCCTTTAGTTGGGCTCATACATTTGGGGTGGTACAGAATCAAAAGCAGCCCTGTTTTCCAAATACCTAAAAACGACGACATTCCTGAGCAAGATAGTCTGGGACTTTCAAATCTTCAGAAGAGCCAAATCCAGGGGAAGTAG<br><br>Translation product:<br>MSLRNLWRDYKVLVVMVPLVGLIHLGWYRIKSSPVFQIPKNDDIPEQDSLGLSNLQKSQIQGK |

*Boxes indicate the translational start and stop codons of each protein.

**table S1**. Nucleotide and protein sequences of SEPs and LEPs in humans (continued)

| No | Gene name | Sequence (5' to 3')* |
|---|---|---|
| 4 | *uKIAA0753* | ATGGCCGTGGGAGTGCCCACAGCTCCCGGGCTTCCAGGGCCAAT GGTAGGGGATACGGGCTCTGGGGATGACGGCTCAGGCATATGGG GAAGGAGACTGGCCACTGAGGGGCCCACAGCTGCCAGGGTTCCT GGACCTGTGGAGGGGGAGACAGGCTCTGAAGGTGTCTCAGGCCT GTGGAGAAGGAGACACGGTGGAGCCATTCCCGAGGCCGGGAGG GTGCCTCTGCCTCTGGGGGTGCTTGCAGCTGTAGGAGCTCCCATG GCAGGGCGTGCGCCTGCTGCGGTGTGGGTGACTGGGTCCTCAGG AGCAGAAGCGGACGTGGGTGTCTCAGGTCTGACCGTGCTGAGGA GACAGTCCACAGGGGGAGTGGGGGCTTCAGAGGATGAGACAGG AGGTGGGAATATTTTGGGACTGGCCGGGGGCAGCCAGGCTGTGG GGACATCTCACACCTGTGTGCCTGGGGCCAGGTTATGGAGCTGTC CAAGGTCTTTGGGGGAAGAAACAGCCTATGAGAATGCTCTAGGC ATGTCAGGGACAAGAACAGCTGTGGGGGTACCCTCAGTTTCAAG GGAGGAAATAGGCCTTGGCCATTTCAGAGACCACCTACAGCCAA GTGGGAGGAGGCTGGCTGGAGGAGTGTCTGCAGTCAGAGTGAGA GGGAGCAATTTGGGAGAGAGTTATGTGGGGGAGGCTAGATTGAG GGGGGCGTTCCAGTAG<br><br>Translation product:<br><br>MAVGVPTAPGLPGPMVGDTGSGDDGSGIWGRRLATEGPTAARVPG PVEGETGSEGVSGLWRRRHGGAIPEAGRVPLPLGVLAAVGAPMAGR APAAVWVTGSSGAEADVGVSGLTVLRRQSTGGVGASEDETGGGNI LGLAGGSQAVGTSHTCVPGARLWSCPRSLGEETAYENALGMSGTRT AVGVPSVSREEIGLGHFRDHLQPSGRRLAGGVSAVRVRGSNLGESY VGEARLRGAFQ |
| 5 | *uZBTB4* | ATGATGTCAGCGGCGGCGGCGGCGGCGGCAGCGGGGCCGGGCC GGGGCCAGGCGCTGGGGGGGCCAGGGCCGGAGCTGGAGCCGGA GCCGGAGCTGGAGGCGGGCAAAACCGGAGCGGCCGGGGGCGGC CACGGTGATGAGCCGGGCCTGGCGGACGCGGCGCCATCGCAGTC AGAAGGAAGAGGCCCAGCAGAAGGGACTCGCTGA<br><br>Translation product:<br><br>MMSAAAAAAAAGPGRGQALGGPGPELEPEPELEAGKTGAAGGGHG DEPGLADAAPSQSEGRGPAEGTR |

*Boxes indicate the translational start and stop codons of each protein.

**table S1**. Nucleotide and protein sequences of SEPs and LEPs in humans (continued)

| No | Gene name | Sequence (5' to 3')* |
|---|---|---|
| 6 | *oC11orf96* | ATGGCATTCGCTGTCATCCGAGCTCAGAGCCGTGTGGGCAGCCG<br>CGGGCTATATAAGCCGCGAGCCTGGCCGCCGCGGGGCAGACGGC<br>GACAGCAGCGGCGGCGAGCGCCTCGGAGCGCGGCGGAACAGCG<br>CCCCCCGAGCCCCGTGCCCCCCGACGGGTCCGCCCGCCCGCCCGC<br>CCTCCCGAGGAGCGCCGGCCCGGGCCCGCGAGGGCCGCCGCCAC<br>CCCGCAGCAGATTTGGATCCCCCGCCCGGCGAGCCCCCGGCTGCT<br>GCCTCCCGGGGGGCCCCGGCGCAGCGGCCGCCCTCGGAGAGCCC<br>CGGCGCCCCGCCGCCCGGCCCCGCAGACGCCGGAGGCGCCATGG<br>CCGCCAAGCCCGGCGAGCTGATGGGCATCTGCTCCAGTTACCAG<br>GCGGTGATGCCGCACTTCGTGTGCCTGGCCGACGAGTTCCCGCAG<br>CCCGTGCGGCCCGCCAAGCTGCCCAAGGGCCGGGGCCGGCTGCG<br>GCGGCCGCGCCAGTCTCGCTTCAAGACGCAGCCGGTGACCTTCGA<br>CGAGATCCAGGAGGTGGAGGAGGAGGGGGTGTCCCCCATGGAGG<br>AGGAGAAGGCCAAGAAGTCGTTCCTGCAGAGCCTGGAGTGCCTG<br>CGCCGCAGCACGCAGAGCCTGTCGCTGCAGCGGGAGCAGCTCAG<br>CAGCTGCAAACTGAGGAACAGCCTGGACTCCAGCGACTCCGACT<br>CGGCCCTGTAAGGGGCGCCGCCCGCGGGGGGGACGCGCGCGTCC<br>GCGGTCCGCGCGGGGACCGGCGTGTGAACCCCGAGAGTGCCCGC<br>GCCCTGCTCCCGGGGGACCCGCAAGGACCCGGGACCGCCGCTCC<br>TCGCGCGCTCGGACTCCCGCCCCGCTGCGAACCGGTCGGTGCGCC<br>CCTCGCCGCGCTCGCCCTGGCCCGGGAGCGCCGGGAGCGGGGCC<br>GCTTTCCTCGTCCTTGTAAATGTTTATTTTTAACTCTTCCCAGTG<br>CGAACTCTGCTGTGAGTGTGTGCGGGGAGGCGCGCCCGCGCTGA<br>GTCGGCGGCGGGTAGCCACTCCATGCCCTTGTCCGATGGTTTGCA<br>ACTCCGATTTGCACACCGCTCCACCGTGCCCCCCAGCGCACACC<br>CATTCACACTCACGCCAACACTCTCGCTGAACACTTTTATAATTG<br>TTAGGCGTGGCCGTTGGGACTTTGGGCGCAGCGCGGCTGCTACTG<br>CGTCTGGAGGATTGATATTTATTTTTGCATTGCGATGGCTGAAGG<br>CATTTATTTAA<br><br>Translation product:<br><br>MAFAVIRAQSRVGSRGLYKPRAWPPRGRRRQQRRRAPRSAAEQRPP<br>SPVPPDGSARPPALPRSAGPGPRGPPPPRSRFGSPARRAPGCCLPGGP<br>GAAAALGEPRRPAARPRRRRRRHGRQARRADGHLLQLPGGDAALR<br>VPGRRVPAARAARQAAQGPGPAAAAAPVSLQDAAGDLRRDPGGGG<br>GGGVPHGGGEGQEVVPAEPGVPAPQHAEPVAAAGAAQQLQTEEQP<br>GLQRLRLGPVRGAARGGDARVRGPRGDRRVNPESARALLPGDPQG<br>PGTAAPRALGLPPRCEPVGAPLAALALARERRERGRFPRPCKCLFFN<br>SSQCELCCECVRGGAPALSRRRVATPCPCPMVCNSDFAHRSTVPPSA<br>HPFTLTPTLSLNTFIIVRRGRWDFGRSAAATASGGLIFIFALRWLKAFI |
| 7 | *oPIDD1* | ATGTCTGGCCTCCAAGGACCGTCGGTGGGCGATGGCTGCAACGG<br>TGGAGGGGCCAGAGCTGGAGGCAGCTGCTGCCGCAGGAGATGCT<br>TCAGAGGATTCGGACGCAGGGTCCAGGGCGCTGCCTTTCCTGGGC<br>GGCAACCGGCTGAGCTTGGACCTGTACCCCGGGGGCTGCCAGCA<br>GCTGCTGCACCTGTGTGTCCAGCAGCCTCTGCAGCTGCTGCAGGT<br>GGAATTCTTGCGTCTGAGCACTCACGAGGACCCTCAGCTGCTGGA<br>GGCCACCCTGGCCCAGCTGCCTCAGAGCCTGTCCTGCCTCCGCTC<br>CCTGGTCCTCAAAGGAGGGCAACGCCGGGACACACTGGGTGCCT<br>GTCTCCGGGGTGCCCTGACCAACCTGCCCGCTGGTCTGAGTGGCC<br>TGGCCCATCTGGCCCACCTGGACCTGAGCTTCAACAGCCTGGAGA<br>CACTGCCGGCCTGTGTCCTGCAGATGCGAGGTCTGGGTGCGCTCT<br>TGCTGTCTCACAACTGCCTCTCTGA<br><br>Translation product:<br><br>MSGLQGPSVGDGCNGGGARAGGSCCRRRCFRGFGRRVQGAAFPGR<br>QPAELGPVPRGLPAAAAPVCPAASAAAAGGILASEHSRGPSAAGGH<br>PGPAASEPVLPPLPGPQRRATPGHTGCLSPGCPDQPARWSEWPGPSG<br>PPGPELQQPGDTAGLCPADARSGCALAVSQLPL |

*Boxes indicate the translational start and stop codons of each protein.

**table S1**. Nucleotide and protein sequences of SEPs and LEPs in humans (continued)

| No | Gene name | Sequence (5' to 3')* |
|---|---|---|
| 8 | *Gdown3* | Transcript variant 1<br>ATGGCGACTCCCGCTCGTGCCCCGGAGTCACCGCCGTCCGCGGAT<br>CCGGCGCTAGTAGCGGGGCCTGCCGAGGAAGCCGAGTGCCCGCC<br>GCCGCGCCAGCCTCAGCCCGCGCAGAATGTGCTCGCTGCCCCGCG<br>GCTTCGAGCCCCAAGCTCCCGAGGACTTGGCGCAGCGGAGTTTG<br>GTGGAGCTGCGGGAAATGTTGAAGCGCCAGGAGAGACTTTTGCG<br>CAACGAAAAATTCATTTGCAAATTGCCCGACAAAGGTAA<br><br>Translation product:<br>MATPARAPESPPSADPALVAGPAEEAECPPPRQPQPAQNVLAAPRLR<br>APSSRGLGAAEFGGAAGNVEAPGETFAQRKIHLQIARQR<br><br>Transcript variant 2<br>ATGGCGACTCCCGCTCGTGCCCCGGAGTCACCGCCGTCCGCGGAT<br>CCGGCGCTAGTAGCGGGGCCTGCCGAGGAAGCCGAGTGCCCGCC<br>GCCGCGCCAGCCTCAGCCCGCGCAGAATGTGCTCGCTGCCCCGCG<br>GCTTCGAGCCCCAAGCTCCCGAGGACTTGGCGCAGCGGAGTTTG<br>GTGGAGCTGCGGGAAATGTTGAAGCGCCAGGAGAGACTTTTGCG<br>CAACGAGTTACAGAGGATCACCATTGCGGACCAAGGTGA<br><br>Translation product:<br>MATPARAPESPPSADPALVAGPAEEAECPPPRQPQPAQNVLAAPRLR<br>APSSRGLGAAEFGGAAGNVEAPGETFAQRVTEDHHCGPR |
| 9 | *oSFRP4* | ATGGCCGTGGCTGGCTGCGCTCCGAGCTGCGGAGTCCGGGACTG<br>GAGCTGCCCGGGCGGGTTCGCGCCCCGAAGGCTGAGAGCTGGCG<br>CTGCTCGTGCCCTGTGTGCCAGACGGCGGAGCTCCGCGGCCGGAC<br>CCCGCGGCCCCGCTTTGCTGCCGACTGGAGTTTGGGGGAAGAAA<br>CTCTCCTGCGCCCCAGAGGATTTCTTCCTCGGCGAAGGGACAGCG<br>AAAGATGAGGGTGGCAGGAAGAGAAGGGCGCTTTCTGTCTGCCG<br>GGGTCGCAGCGCGAGAGGGCAGTGCCATGTTCCTCTCCATCCTAG<br>TGGCGCTGTGCCTGTGGCTGCACCTGGCGCTGGGCGTGCGCGGCG<br>CGCCCTGCGAGGCGGTGCGCATCCCTATGTGCCGGCACATGCCCT<br>GGAACATCACGCGGATGCCCAACCACCTGCACCACAGCACGCAG<br>GAGAACGCCATCCTGGCCATCGAGCAGTACGAGGAGCTGGTGGA<br>CGTGAACTGCAGCGCCGTGCTGCGCTTCTTCCTCTGTGCCATGTA<br>CGCGCCCATTTGCACCCTGGAGTTCCTGCACGACCCTATCAAGCC<br>GTGCAAGTCGGTGTGCCAACGCGCGCGCGACGACTGCGAGCCCC<br>TCATGAAGATGTACAACCACAGCTGGCCCGAAAGCCTGGCCTGC<br>GACGAGCTGCCTGTCTATGA<br><br>Translation product:<br>MAVAGCAPSCGVRDWSCPGGFAPRRLRAGAARALCARRRSSAAGP<br>RGPALLPTGVWGKKLSCAPEDFFLGEGTAKDEGGRKRRALSVCRGR<br>SARGQCHVPLHPSGAVPVAAPGAGRARRALRGGAHPYVPAHALEH<br>HADAQPPAPQHAGERHPGHRAVRGAGGRELQRRAALLPLCHVRAH<br>LHPGVPARPYQAVQVGVPTRARRLRAPHEDVQPQLARKPGLRRAA<br>CL |

*Boxes indicate the translational start and stop codons of each protein.

**table S1**. Nucleotide and protein sequences of SEPs and LEPs in humans (continued)

| No | Gene name | Sequence (5' to 3')* |
|---|---|---|
| 10 | *oPMEPA1* | ATGAATTCGCTCTGGTCTAGCGGAGCTGGAGTTTGTTCAGATCAT<br>CATCATCGTGGTGGTGATGATGGTGATGGTGGTGGTGATCACGTG<br>CCTGCTGAGCCACTACAAGCTGTCTGCACGGTCCTTCATCAGCCG<br>GCACAGCCAGGGGCGGAGGAGAGAAGATGCCCTGTCCTCAGAAG<br>GATGCCTGTGGCCCTCGGAGAGCACAGTGTCAGGCAACGGAATC<br>CCAGAGCCGCAGGTCTACGCCCCGCCTCGGCCCACCGACCGCCTG<br>GCCGTGCCGCCCTTCGCCCAGCGGGAGCGCTTCCACCGCTTCCAG<br>CCCACCTATCCGTACCTGCAGCACGAGATCGACCTGCCACCCACC<br>ATCTCGCTGTCAGACGGGGAGGAGCCCCCACCCTACCAGGGCCC<br>CTGCACCCTCCAGCTTCGGGACCCCGAGCAGCAGCTGGAACTGA<br>ACCGGGAGTCGGTGCGCGCACCCCCAAACAGAACCATCTTCGAC<br>AGTGA<br><br>Translation product:<br>MNSLWSSGAGVCSDHHHRGGDDGDGGGDHVPAEPLQAVCTVLHQ<br>PAQPGAEERRCPVLRRMPVALGEHSVRQRNPRAAGLRPASAHRPPG<br>RAALRPAGALPPLPAHLSVPAARDRPATHHLAVRRGGAPTLPGPLHP<br>PASGPRAAAGTEPGVGARTPKQNHLRQ |
| 11 | *oFAM163B* | ATGGAGAAGGGGGCGGATGACAGCCGGGACCGTGGTCATCACCG<br>GGGGCATCTTGGCGACTGTGATTTTGCTCTGCATCATCGCTGTTCT<br>GTGCTACTGCCGGCTCCAGTACTACTGCTGCAAGAAGGACGAGTC<br>GGAGGAGGACGAGGAGGAACCAGACTTCGCCGTTCACTCGCACC<br>TGCCCCCGCTGCACTCCAACCGCAACCTGGTGCTGACCAACGGGC<br>CGGCGCTCTACCCCACCGCCTCCACCTCCTTCAGCCAGAAGTCCC<br>CGCAGGCCCGCGCCCTCTGCCGCAGCTGCTCCCACTGCGAGCCCC<br>CCACCTTCTTCCTGCAGGAGCCGCCGGAGGAGGAAGAGGACGTG<br>CTGAACGGCGGGGAGCGCGTGCTCTACAAGAGCGTGAGCCAGGA<br>GGACGTGGAGCTGCCCCCGGGGGGCTTCGGGGGCCTGCAGGCGC<br>TCAACCCCAACCGCCTCTCAGCCATGCGGGAGGCCTTCGCCAGGA<br>GCCGCAGCATCAGCACCGACGTGTGACCTGGGCCCGCCCCCGGG<br>ATCCTGGCCTTGAAGGTCCTGAGACAATGGGGGGATTCCTTTGGC<br>ACACCCACCTGGCCCCAGGCCCCAAGTGGGGCTTGGCTGAGCCCT<br>CCTGGAATCCCTGTCCTCAGGGGAGGCCCAGGCTTGGGGAGCCC<br>CCAGGCTCGTGGCAGGGATAA<br><br>Translation product:<br>MEKGADDSRDRGHHRGHLGDCDFALHHRCSVLLPAPVLLLQEGRV<br>GGGRGGTRLRRSLAPAPAALQPQPGADQRAGALPHRLHLLQPEVPA<br>GPRPLPQLLPLRAPHLLPAGAAGGGRGRAERRGARALQEREPGGRG<br>AAPGGLRGPAGAQPQPPLSHAGGLRQEPQHQHRRVTWARPRDPGL<br>EGPETMGGFLWHTHLAPGPKWGLAEPSWNPCPQGRPRLGEPPGSW<br>QG |
| 12 | *oBEGAIN* | ATGCACTGTCCGTGTTTATTGAGCGCCTGCTGTATGCTGGGACCT<br>AGCATGAGAAGCGCTGTCATGGGGACCCGGAAAGGCCTCTGCCG<br>CAGACATGGAGAAACTCAGCGCGCTGCAGGAGCAGAAGGGCGA<br>GCTGCGCAAGCGGCTGTCCTACACCACACACAAGCTCGAGAAGC<br>TCGAGACCGAGTTCGACTCCACGCGCCACTACCTGGAGATCGAG<br>CTGCGGCGCGCGCAGGAGGAACTGGAGAAGGTCACGGAGAAGCT<br>GCGCAGGATTCAGAGCAACTACATGGCACTGCAGAGGATCAACC<br>AGGAGCTGGAGGACAAGCTGTACCGCATGGGCCAGCACTATGA<br><br>Translation product:<br>MHCPCLLSACCMLGPSMRSAVMGTRKGLCRRHGETQRAAGAEGR<br>AAQAAVLHHTQAREARDRVRLHAPLPGDRAAARAGGTGEGHGEA<br>AQDSEQLHGTAEDQPGAGGQAVPHGPAL |

*Boxes indicate the translational start and stop codons of each protein.

**table S2**. List of SEPs and LEPs in Concatemer 1/2

| No | Protein name | Concatemer |
|---|---|---|
| 1 | oSCRIB | 1 |
| 2 | alt-RPL36 | 1 |
| 3 | uMKKS1 | 1 |
| 4 | uKIAA0753 | 1 |
| 5 | uZBTB4 | 1 |
| 6 | oC11orf96 | 1 |
| 7 | oPIDD1 WT(A70)/mt(S70) | 2 |
| 8 | Gdown3 | 2 |
| 9 | oSFRP4 | 2 |
| 10 | oPMEPA1 | 2 |
| 11 | oFAM163B | 2 |
| 12 | oBEGAIN | 2 |

**table S3**. List of plasmids

| No | Name | Sequence (5' to 3')* |
|---|---|---|
| 1 | pEX-A2J2-Concatemer1 | AGATATCCATATGTTTACTAGTAACAATGGGACATCATCATCATCATCACGATTACGACATCCCA<br>ACGACCGAAAACCTGTATTTTCAGGGCAAGGCTGCTCATCCGCACCATGCACAAGTCCACCCTGC<br>CGTAGCGTTACAGCCGGCCCGCGGTGTCGGCGGACAAGCGGCTTTGTTCGCGGCTGGAAGAGCC<br>GGTGGAGACCTGCCATTGCAGCCGCAGCCTGGGGGAGCAGCCGCTCGCGCAGCTCAAGCCTTCT<br>TCCCAGCGGCGGAGCTAGCGCAGGCGGGGCCCGAGAGGGCCGCGGCTCGGCATGGGGTCACAG<br>AAGGCCTTCAAGGCCAGACGGGCCCCCAAATTTATCAGGAGAAAAGTAGCCCAGTCTTTCAAAT<br>TCCAAAAAATGATGACATACCAGAACAAGACTCCCTTGGTCTTTCGAACCTGCAGAAGGTGCCG<br>GGTCCGGTCGAAGGAGAGACGGGGTCCGAGGGAGTGAGCGGCTTGTGGCGAAGCCTGGGTGAG<br>GAAACAGCGTACGAGAACGCCTTAGGTATGAGCGGCACTCGTGGCTCGAACTTGGGCGAGAGCT<br>ACGTCGGTGAAGCTCGGGGTCAAGCGCTCGGGGGCCCCGGCCCAGAGCTAGAACCTGAGCCGGA<br>GCTTGAGGCCGGTAAGACCGGTGCCGCCGGGGGCGGGCACGGAGACGAACCTGGCCTCGCTGAT<br>GCTGCCCCATCACAATCTGAGGGCAGGGGTCCTGCCGAGGGTACTAGAGCAGCTGCTCGCAGGG<br>CACCAGGTTGCTGCCTGCCTGGCGGCCCAGGCGCGGCAGCCGCTCTCGGAGAACCCCGCCAAGC<br>CGCACAAGGACCTGGACCCGCCGCCGCCGCTGCCCCGGTATCTCTCCAAGACGCAGCGGGGGAC<br>CTCCGTCAGGCCGCGCAAGGGCCCGGACCAGCAGCCGCGGCTGCGCCAGTTTCACTCCAAGATG<br>CCGCAGGCGACCTGAGGAGAGCGGCGGCTCGGCGAGTGAATCCTGAGTCTGCACGGGCACTCTT<br>GCCAGGCGATCCTCAGGGCCCCGGAACCGCCGCGCCCCGGCTGGCTGGGGACGGCGCTCTGCAA<br>AGGCCCGCCGGCGTTGGCTGGCAGGTGGGCAGATCAGGGCCCGAACCCCTGCCTATTCCGGCCC<br>GCGCGGTGGCCGGAGCCGCGGCGGGCGCCGGGGGAAGGCGCGCCGCTGCGAGGATGGTGCCGG<br>AGCCGGGCGATCACGCAGGCGTGGCCGCTGGCGAAGCTGCCGGACGGGTGCACGGCCAAGACG<br>CCACCGATGATTGTGAACTGGCCGCGGCTGCCGCCGGCGAACTGGGCGAGGGCCCCGGCTGAGT<br>CGACTTTCCCGGGTAACTAACTAAGGATCCTTTGAGCTCGGTACCGATATCAATGCCGTGAGCAC<br>AAAAGCGACCATAGTCGGGATTTTACCCGCTTATCTGCGGAAGGTCATTGGGAACGCATCTTCCA<br>CAAATGGGCTGAGCGCAAAAATGCCGAACGTATCTGGCTGACCTATGCCATTCCGTGGGAACGA<br>CACTACTTTTCAATTGGTGCACTCTCAGTACAATCTGCTCTGATGCCGCATAGTTAAGCCAGCCCC<br>GACACCCGCCAACACCCGCTGACGCGCCCTGACGGGCTTGTCTGCTCCCGGCATCCGCTTACAGA<br>CAAGCTGTGACCGTCTCCGGAAAATCAAAGGATCTTCTTGAGATCCTTTTTTTGAGCTGCATGTGT<br>CAGAGGTTTTCACCGTCATCACCGAAACGCGCGAGACGAAAGGGCCTCGTGATACGCCTATTTTT<br>ATAGGTTAATGTCATGATAATAATGGTTTCTTAGACGTCAGGTGGCACTTTTCGGGGAAATGTGC<br>GCGGAACCCCTATTTGTTTATTTTTCTAAATACATTCAAATATGTATCCGCTCATGAGACAATAAC<br>CCTGATAAATGCTTCAATAATATTGAAAAAGGAAGAGTATGAGTATTCAACATTTCCGTGTCGCC<br>CTTATTCCCTTTTTTGCGGCATTTTGCCTTCCTGTTTTTGCTCACCCAGAAACGCTGGTGAAAGTA<br>AAAGATGCTGAAGATCAGTTGGGTGCACGAGTGGGTTACATCGAACTGGATCTCAACAGCGGTA<br>AGATCCTTGAGAGTTTTCGCCCCGAAGAACGTTTTCCAATGATGAGCACTTTTAAAGTTCTGCTAT<br>GTGGCGCGGTATTATCCCGTATTGACGCCGGGCAAGAGCAACTCGGTCGCCGCATACACTATTCT<br>CAGAATGACTTGGTTGAGTACTCACCAGTCACAGAAAAGCATCTTACGGATGGCATGACAGTAA<br>GAGAATTATGCAGTGCTGCCATAACCATGAGTGATAACACTGCGGCCAACTTACTTCTGACAACG<br>ATCGGAGGACCGAAGGAGCTAACCGCTTTTTTGCACAACATGGGGGATCATGTAACTCGCCTTGA<br>TCGTTGGGAACCGGAGCTGAATGAAGCCATACCAAACGACGAGCGTGACACCACGATGCCTGTA<br>GCAATGGCAACAACGTTGCGCAAACTATTAACTGGCGAACTACTTACTCTAGCTTCCCGGCAACA<br>ATTAATAGACTGGATGGAGGCGGATAAAGTTGCAGGACCACTTCTGCGCTCGGCCCTTCCGGCTG<br>GCTGGTTTATTGCTGATAAATCTGGAGCCGGTGAGCGTGGGAGTCGCGGTATCATTGCAGCACTG<br>GGGCCAGATGGTAAGCCCTCCCGTATCGTAGTTATCTACACGACGGGGAGTCAGGCAACTATGG<br>ATGAACGAAATAGACAGATCGCTGAGATAGGTGCCTCACTGATTAAGCATTGGTAACTGTCAGA<br>CCAAGTTTACTCATATATACTTTAGATTGATTTAAAACTTCATTTTTAATTTAAAAGGATCTAGGT<br>GAAGATCCTTTTTGATAATCTCATGACCAAAATCCCTTAACGTGAGTTTTCGTTCCACTGAGCGTC<br>AGACCCCGTAGAAAAGATCAAAGGATCTTCTTGAGATCCTTTTTTTCTGCGCGTAATCTGCTGCTT<br>GCAAACAAAAAAACCACCGCTACCAGCGGTGGTTTGTTTGCCGGATCAAGAGCTACCAACTCTTT<br>TTCCGAAGGTAACTGGCTTCAGCAGAGCGCAGATACCAAATACTGTTCTTCTAGTGTAGCCGTAG<br>TTAGGCCACCACTTCAAGAACTCTGTAGCACCGCCTACATACCTCGCTCTGCTAATCCTGTTACCA<br>GTGGCTGCTGCCAGTGGCGATAAGTCGTGTCTTACCGGGTTGGACTCAAGACGATAGTTACCGGA<br>TAAGGCGCAGCGGTCGGGCTGAACGGGGGGTTCGTGCACACAGCCCAGCTTGGAGCGAACGACC<br>TACACCGAACTGAGATACCTACAGCGTGAGCTATGAGAAAGCGCCACGCTTCCCGAAGGGAGAA<br>AGGCGGACAGGTATCCGGTAAGCGGCAGGGTCGGAACAGGAGAGCGCACGAGGGAGCTTCCAG<br>GGGGAAACGCCTGGTATCTTTATAGTCCTGTCGGGTTTCGCCACCTCTGACTTGAGCGTCGATTTT<br>TGTGATGCTCGTCAGGGGGGCGGAGCCTATGGAAAAACGCCAGCAACGCGGCCTTTTTACGGTT<br>CCTGGCCTTTTGCTGGCCTTTTGCTCACATGTTCTTTCCTGCGTTATCCCCTGATTCTGTGGATAAC<br>CGTATTACCGCCTTTGAGTGAGCTGATACCGCTCGCCGCAGCCGAACGACCGAGCGCAGCGAGT<br>CAGTGAGCGAGGAAGCGGAAGAGCAAATAATAAAAAAGCCGGATTAATAATCTGGCTTTTTATA<br>TTCTCTGCCCAATACGCAAACCGCCTCTCCCCGCGCGTTGGCCGATTCATTAATGCAGCTGGCAC<br>GACAGGCAAAGACTTCTACGGTTGGGCAATCGAACGCCACTATTCAGCGCATGACCTGTTTAAG<br>GCGCATAGCTTTGATCATGCCGGTAGAGCACGTTGGGAAGGACATCGTTGGGCACATCGTGAAC<br>ATGCCAACATGTTTTGGGCGCATGAAGGCATTTGGCGTATCCACTGACT |

*Boxes indicate the translational start and stop codons of the protein (Concatemer1). Underlines indicate the primer annealing sites. Gray highlight indicates the pEX-A2J2 vector sequence.

**table S3**. List of plasmids (continued)

| No | Name | Sequence (5' to 3')* |
|---|---|---|
| 2 | pEX-A2J2-Concatemer2 | AGATATCCATATGTTTACTAGTAACAATGGGACATCATCATCATCATCACGATTACGACATCCCA<br>ACGACCGAAAACCTGTATTTTCAGGGCAAGGGCTTGCCAGCTGCGGCAGCACCAGTATGTCCTGC<br>GGCATCGGCCGCGGCGGCAGGGGGCATCCTGGCTTCGGAACACAGCCGCGGATTACCAGCCGCC<br>GCAGCCCCAGTGTGCCCTGCCGCATCCAGCGCTGCTGCTGGTGGAATTCTCGCATCCGAGCATTC<br>CAGGGGGCCATCGGCCGCCGGTGGGCATCCTGGACCCGCGGCTTCTGAGCCCGTGCTTCCCCCTC<br>TGCCTGGTCCTCAACGTGGACCAAGTGCGGCCGGTGGGCATCCGGGGCCAGCAGCATCAGAGCC<br>AGTTCTACCCCCACTCCCAGGTCCGCAAAGACGAGCCGCTGCTCGTGCAACCCCGGCTAGAGCCC<br>CAGAGTCACCCCCGAGCGCTGATCCTGCGCTAGTCGCAGGCCCGGCCGAAGAAGCGGAGTGTCC<br>TCCCCCGAGACAGCCCCAACCAGCGCAGAACGTTCTTGCTGCGCCTAGGGGTCTCGGTGCGGCA<br>GAATTCGGTGGGGCGGCCGGTAACGTCGAGGCCCCAGGGGAGACATTCGCTCAACGGTTATCTT<br>GTGCACCGGAGGACTTCTTTTTGGGTGAGGGGACCGCGAAGGCGCCGCACGAGGACGTTCAGCC<br>GCAGCTAGCTCGGGCGGCGGCGCGTAGGATGCCAGTCGCACTTGGAGAGCACTCCGTCCGCGCG<br>GCCGCCGGGACGGAACCGGGAGTGGGCGCCCGCGGCCATCTTGGCGACTGCGACTTCGCCCTCC<br>ATCATCGATCATTGGCTCCAGCTCCCGCGGCTCTCCAGCCCCAGCCCGGCGCCGATCAACGCGCG<br>GCCGCTCGCGATCCGGGTCTCGAGGGACCCGAGACTATGGGAGGATTTCTCTGGCACACCCACCT<br>CGCGCCGGGACCGAAGTGGGGCCTGGCCGAACCCAGCTGGAATCCGTGTCCCCAGGGGAGGCCG<br>CGCGCGGCCCAAGCCGCCGTGCTGCACCACACTCAAGCTCGGCTGCACGCCCCACTGCCAGGCG<br>ACCGGGCCGGCGGAACGGGCGAAGGTCACGGCGAGGCCGCCCAAGACAGTGAGCAGCTCCACG<br>GCACAGCCGAAGATCAACCAGGCGCCGGCGGCCAGGCCGTGCCACACGGCCCAGCCCTCTGAGT<br>CGACTTTCCCGGGTAACTAACTAAGGATCCTTTGAGCTCGGTACCGATATCAATGCCGTGAGCAC<br>AAAAGCGACCATAGTCGGGATTTTACCCGCTTATCTGCGGAAGGTCATTGGGAACGCATCTTCCA<br>CAAATGGGCTGAGCGCAAAAATGCCGAACGTATCTGGCTGACCTATGCCATTCCGTGGGAACGA<br>CACTACTTTTCAATTGGTGCACTCTCAGTACAATCTGCTCTGATGCCGCATAGTTAAGCCAGCCCC<br>GACACCCGCCAACACCCGCTGACGCGCCCTGACGGGCTTGTCTGCTCCCGGCATCCGCTTACAGA<br>CAAGCTGTGACCGTCTCCGGAAAATCAAAGGATCTTCTTGAGATCCTTTTTTTGAGCTGCATGTGT<br>CAGAGGTTTTCACCGTCATCACCGAAACGCGCGAGACGAAAGGGCCTCGTGATACGCCTATTTTT<br>ATAGGTTAATGTCATGATAATAATGGTTTCTTAGACGTCAGGTGGCACTTTTCGGGGAAATGTGC<br>GCGGAACCCCTATTTGTTTATTTTTCTAAATACATTCAAATATGTATCCGCTCATGAGACAATAAC<br>CCTGATAAATGCTTCAATAATATTGAAAAAGGAAGAGTATGAGTATTCAACATTTCCGTGTCGCC<br>CTTATTCCCTTTTTTGCGGCATTTTGCCTTCCTGTTTTTGCTCACCCAGAAACGCTGGTGAAAGTA<br>AAAGATGCTGAAGATCAGTTGGGTGCACGAGTGGGTTACATCGAACTGGATCTCAACAGCGGTA<br>AGATCCTTGAGAGTTTTCGCCCCGAAGAACGTTTTCCAATGATGAGCACTTTTAAAGTTCTGCTAT<br>GTGGCGCGGTATTATCCCGTATTGACGCCGGGCAAGAGCAACTCGGTCGCCGCATACACTATTCT<br>CAGAATGACTTGGTTGAGTACTCACCAGTCACAGAAAAGCATCTTACGGATGGCATGACAGTAA<br>GAGAATTATGCAGTGCTGCCATAACCATGAGTGATAACACTGCGGCCAACTTACTTCTGACAACG<br>ATCGGAGGACCGAAGGAGCTAACCGCTTTTTTGCACAACATGGGGGATCATGTAACTCGCCTTGA<br>TCGTTGGGAACCGGAGCTGAATGAAGCCATACCAAACGACGAGCGTGACACCACGATGCCTGTA<br>GCAATGGCAACAACGTTGCGCAAACTATTAACTGGCGAACTACTTACTCTAGCTTCCCGGCAACA<br>ATTAATAGACTGGATGGAGGCGGATAAAGTTGCAGGACCACTTCTGCGCTCGGCCCTTCCGGCTG<br>GCTGGTTTATTGCTGATAAATCTGGAGCCGGTGAGCGTGGGAGTCGCGGTATCATTGCAGCACTG<br>GGGCCAGATGGTAAGCCCTCCCGTATCGTAGTTATCTACACGACGGGGAGTCAGGCAACTATGG<br>ATGAACGAAATAGACAGATCGCTGAGATAGGTGCCTCACTGATTAAGCATTGGTAACTGTCAGA<br>CCAAGTTTACTCATATATACTTTAGATTGATTTAAAACTTCATTTTTAATTTAAAAGGATCTAGGT<br>GAAGATCCTTTTTGATAATCTCATGACCAAAATCCCTTAACGTGAGTTTTCGTTCCACTGAGCGTC<br>AGACCCCGTAGAAAAGATCAAAGGATCTTCTTGAGATCCTTTTTTTCTGCGCGTAATCTGCTGCTT<br>GCAAACAAAAAAACCACCGCTACCAGCGGTGGTTTGTTTGCCGGATCAAGAGCTACCAACTCTTT<br>TTCCGAAGGTAACTGGCTTCAGCAGAGCGCAGATACCAAATACTGTTCTTCTAGTGTAGCCGTAG<br>TTAGGCCACCACTTCAAGAACTCTGTAGCACCGCCTACATACCTCGCTCTGCTAATCCTGTTACCA<br>GTGGCTGCTGCCAGTGGCGATAAGTCGTGTCTTACCGGGTTGGACTCAAGACGATAGTTACCGGA<br>TAAGGCGCAGCGGTCGGGCTGAACGGGGGGTTCGTGCACACAGCCCAGCTTGGAGCGAACGACC<br>TACACCGAACTGAGATACCTACAGCGTGAGCTATGAGAAAGCGCCACGCTTCCCGAAGGGAGAA<br>AGGCGGACAGGTATCCGGTAAGCGGCAGGGTCGGAACAGGAGAGCGCACGAGGGAGCTTCCAG<br>GGGGAAACGCCTGGTATCTTTATAGTCCTGTCGGGTTTCGCCACCTCTGACTTGAGCGTCGATTTT<br>TGTGATGCTCGTCAGGGGGGCGGAGCCTATGGAAAAACGCCAGCAACGCGGCCTTTTTACGGTT<br>CCTGGCCTTTTGCTGGCCTTTTGCTCACATGTTCTTTCCTGCGTTATCCCCTGATTCTGTGGATAAC<br>CGTATTACCGCCTTTGAGTGAGCTGATACCGCTCGCCGCAGCCGAACGACCGAGCGCAGCGAGT<br>CAGTGAGCGAGGAAGCGGAAGAGCAAATAATAAAAAAGCCGGATTAATAATCTGGCTTTTTATA<br>TTCTCTGCCCAATACGCAAACCGCCTCTCCCCGCGCGTTGGCCGATTCATTAATGCAGCTGGCAC<br>GACAGGCAAAGACTTCTACGGTTGGGCAATCGAACGCCACTATTCAGCGCATGACCTGTTTAAG<br>GCGCATAGCTTTGATCATGCCGGTAGAGCACGTTGGGAAGGACATCGTTGGGCACATCGTGAAC<br>ATGCCAACATGTTTTGGGCGCATGAAGGCATTTGGCGTATCCACTGACT |

*Boxes indicate the translational start and stop codons of the protein (Concatemer2). Underlines indicate the primer annealing sites. Gray highlight indicates the pEX-A2J2 vector sequence.

**table S3**. List of plasmids (continued)

| No | Name | Sequence (5' to 3')* |
|---|---|---|
| 3 | pEX-A2J2-*oPIDD1*-*PIDD1*-*sfGFP* | GAATTCACGCGTAGGCGCCGAGCGCCCGCTGAGCAGCCACCCTTTGCGCGCCGCCTGCAGCGCAGCTT<br>CCCCGGGCGCTGCCTGGACAGGCCTGCCTGCGTGCTGGGACATGTCTGGCCTCCAAGGACCGTCGGTG<br>GGCGATGGCTGCAACGGTGGAGGGGCCAGAGCTGGAGGCAGCTGCTGCCGCAGGAGATGCTTCAGAG<br>GATTCGGACGCAGGGTCCAGGGCGCTGCCTTTCCTGGGCGGCAACCGGCTGAGCTTGGACCTGTACCC<br>CGGGGGCTGCCAGCAGCTGCTGCACCTGTGTGTCCAGCAGCCTCTGCAGCTGCTGCAGGTGGAATTCTT<br>GCGTCTGAGCACTCACGAGGACCCTCAGCTGCTGGAGGCCACCCTGGCCCAGCTGCCTCAGAGCCTGT<br>CCTGCCTCCGCTCCCTGGTCCTCAAAGGAGGGCAACGCCGGGACACACTGGGTGCCTGTCTCCGGGGT<br>GCCCTGACCAACCTGCCCGCTGGTCTGAGTGGCCTGGCCCATCTGGCCCACCTGGACCTGAGCTTCAAC<br>AGCCTGGAGACACTGCCGGCCTGTGTCCTGCAGATGCGAGGTCTGGGTGCGCTCTTGCTGTCTCACAAC<br>TGCCTCTCTGAGGGTTCTGCTGGAAGCGCAGCCGGCTCCGGGGAGTTCTCTAAGGGTGAGGAGCTATT<br>CACAGGCGTGGTACCCATCTTGGTAGAGCTGGACGGGGACGTCAACGGTCACAAATTCTCAGTGCGAG<br>GAGAGGGCGAGGGTGACGCTACTAACGGCAAGTTAACACTGAAGTTTATCTGCACGACAGGCAAGTTG<br>CCCGTTCCTTGGCCAACGCTTGTCACTACCCTGACTTACGGGGTGCAGTGTTTCAGTCGCTACCCGGAC<br>CATATGAAAAGACACGACTTTTTAAGTCCGCGATGCCCGAAGGCTACGTTCAAGAGCGGACAATATC<br>CTTTAAAGACGACGGTACCTACAAGACACGTGCCGAAGTCAAGTTCGAGGGAGATACTTTAGTCAACA<br>GGATCGAACTCAAAGGAATTGACTTCAAAGAGGACGGGAATATCCTCGGACATAAACTGGAGTACAAC<br>TTTAACTCGCACAACGTTTATATCACTGCCGATAAACAGAAGAACGGAATAAAAGCCAACTTTAAAAT<br>TAGACACAACGTGGAAGACGGAAGCGTGCAGCTCGCAGACCATTATCAACAGAACACCCCCATTGGAG<br>ACGGACCAGTGCTGCTGCCAGATAATCATTATCTCTCTACCCAGAGTGTGCTGAGCAAGGATCCAAAC<br>GAAAAGAGGGATCATATGGTGCTGCTTGAGTTCGTGACCGCTGCAGGGATTACCCACGGCATGGACGA<br>ACTTTATAAGGGCTCACACCACCACCATCACCATTAGGCCCCACAGACTTTTAGGCTGGCCCAGATATT<br>CCCCAGTGGATGGGCAGAGCCCCCACCTTCAAGTCTCTCCAGTGTGTGGGGACGGGTCCCTGTGAGCA<br>ACAAAACTGCACTGTTTCTTTCACCTCGTCGACCCCGGGATGCCGTGAGCACAAAAGCGACCATAGTCG<br>GGATTTTACCCGCTTATCTGCGGAAGGTCATTGGGAACGCATCTTCCACAAATGGGCTGAGCGCAAAA<br>ATGCCGAACGTATCTGGCTGACCTATGCCATTCCGTGGGAACGACACTACTTTTCAATTGGTGCACTCT<br>CAGTACAATCTGCTCTGATGCCGCATAGTTAAGCCAGCCCCGACACCCGCCAACACCCGCTGACGCGC<br>CCTGACGGGCTTGTCTGCTCCCGGCATCCGCTTACAGACAAGCTGTGACCGTCTCCGGAAAATCAAAGG<br>ATCTTCTTGAGATCCTTTTTTTGAGCTGCATGTGTCAGAGGTTTTCACCGTCATCACCGAAACGCGCGAG<br>ACGAAAGGGCCTCGTGATACGCCTATTTTTATAGGTTAATGTCATGATAATAATGGTTTCTTAGACGTC<br>AGGTGGCACTTTTCGGGGAAATGTGCGCGGAACCCCTATTTGTTTATTTTTCTAAATACATTCAAATAT<br>GTATCCGCTCATGAGACAATAACCCTGATAAATGCTTCAATAATATTGAAAAAGGAAGAGTATGAGTA<br>TTCAACATTTCCGTGTCGCCCTTATTCCCTTTTTTGCGGCATTTTGCCTTCCTGTTTTTGCTCACCCAGAA<br>ACGCTGGTGAAAGTAAAAGATGCTGAAGATCAGTTGGGTGCACGAGTGGGTTACATCGAACTGGATCT<br>CAACAGCGGTAAGATCCTTGAGAGTTTTCGCCCCGAAGAACGTTTTCCAATGATGAGCACTTTTAAAGT<br>TCTGCTATGTGGCGCGGTATTATCCCGTATTGACGCCGGGCAAGAGCAACTCGGTCGCCGCATACACTA<br>TTCTCAGAATGACTTGGTTGAGTACTCACCAGTCACAGAAAAGCATCTTACGGATGGCATGACAGTAA<br>GAGAATTATGCAGTGCTGCCATAACCATGAGTGATAACACTGCGGCCAACTTACTTCTGACAACGATC<br>GGAGGACCGAAGGAGCTAACCGCTTTTTTGCACAACATGGGGGATCATGTAACTCGCCTTGATCGTTG<br>GGAACCGGAGCTGAATGAAGCCATACCAAACGACGAGCGTGACACCACGATGCCTGTAGCAATGGCA<br>ACAACGTTGCGCAAACTATTAACTGGCGAACTACTTACTCTAGCTTCCCGGCAACAATTAATAGACTGG<br>ATGGAGGCGGATAAAGTTGCAGGACCACTTCTGCGCTCGGCCCTTCCGGCTGGCTGGTTTATTGCTGAT<br>AAATCTGGAGCCGGTGAGCGTGGGAGTCGCGGTATCATTGCAGCACTGGGGCCAGATGGTAAGCCCTC<br>CCGTATCGTAGTTATCTACACGACGGGGAGTCAGGCAACTATGGATGAACGAAATAGACAGATCGCTG<br>AGATAGGTGCCTCACTGATTAAGCATTGGTAACTGTCAGACCAAGTTTACTCATATATACTTTAGATTG<br>ATTTAAAACTTCATTTTTAATTTAAAAGGATCTAGGTGAAGATCCTTTTTGATAATCTCATGACCAAAAT<br>CCCTTAACGTGAGTTTTCGTTCCACTGAGCGTCAGACCCCGTAGAAAAGATCAAAGGATCTTCTTGAGA<br>TCCTTTTTTTCTGCGCGTAATCTGCTGCTTGCAAACAAAAAAACCACCGCTACCAGCGGTGGTTTGTTTG<br>CCGGATCAAGAGCTACCAACTCTTTTTCCGAAGGTAACTGGCTTCAGCAGAGCGCAGATACCAAATAC<br>TGTTCTTCTAGTGTAGCCGTAGTTAGGCCACCACTTCAAGAACTCTGTAGCACCGCCTACATACCTCGC<br>TCTGCTAATCCTGTTACCAGTGGCTGCTGCCAGTGGCGATAAGTCGTGTCTTACCGGGTTGGACTCAAG<br>ACGATAGTTACCGGATAAGGCGCAGCGGTCGGGCTGAACGGGGGGTTCGTGCACACAGCCCAGCTTGG<br>AGCGAACGACCTACACCGAACTGAGATACCTACAGCGTGAGCTATGAGAAAGCGCCACGCTTCCCGAA<br>GGGAGAAAGGCGGACAGGTATCCGGTAAGCGGCAGGGTCGGAACAGGAGAGCGCACGAGGGAGCTTC<br>CAGGGGGAAACGCCTGGTATCTTTATAGTCCTGTCGGGTTTCGCCACCTCTGACTTGAGCGTCGATTTTT<br>GTGATGCTCGTCAGGGGGGCGGAGCCTATGGAAAAACGCCAGCAACGCGGCCTTTTTACGGTTCCTGG<br>CCTTTTGCTGGCCTTTTGCTCACATGTTCTTTCCTGCGTTATCCCCTGATTCTGTGGATAACCGTATTACC<br>GCCTTTGAGTGAGCTGATACCGCTCGCCGCAGCCGAACGACCGAGCGCAGCGAGTCAGTGAGCGAGGA<br>AGCGGAAGAGCAAATAATAAAAAAGCCGGATTAATAATCTGGCTTTTTATATTCTCTGCCCAATACGC<br>AAACCGCCTCTCCCCGCGCGTTGGCCGATTCATTAATGCAGCTGGCACGACAGGCAAAGACTTCTACG<br>GTTGGGCAATCGAACGCCACTATTCAGCGCATGACCTGTTTAAGGCGCATAGCTTTGATCATGCCGGTA<br>GAGCACGTTGGGAAGGACATCGTTGGGCACATCGTGAACATGCCAACATGTTTTGGGCGCATGAAGGC<br>ATTTGGCGTATCCACTGACT |

*Boxes indicate the translational start and stop codons of the proteins (oPIDD1 and PIDD1-sfGFP). Green, pink, and yellow highlights correspond to *PIDD1* (C-terminally truncated), flexible linker, and *sfGFP* sequences, respectively. Underlines indicate the primer annealing sites. Gray highlight indicates the pEX-A2J2 vector sequence.

**table S3**. List of plasmids (continued)

| No | Name | Sequence (5' to 3')* |
|---|---|---|
| 4 | pEX-A2J2-*oSFRP4*-*SFRP4*-*sfGFP* | GAATTCACGCGTGGGGGAGCCCGCGCCGCGGCTGCAGCTGCCAAGGGAGCGTTCCGAGCCCACGTCAG<br>GGGAGGTGTCGGGATAAATAGGGTCCCGCAATGGCCGTGGCTGGCTGCGCTCCGAGCTGCGGAGTCCG<br>GGACTGGAGCTGCCCGGGCGGGTTCGCGCCCCGAAGGCTGAGAGCTGGCGCTGCTCGTGCCCTGTGTG<br>CCAGACGGCGGAGCTCCGCGGCCGGACCCCGCGGCCCCGCTTTGCTGCCGACTGGAGTTTGGGGGAAG<br>AAACTCTCCTGCGCCCCAGAGGATTTCTTCCTCGGCGAAGGGACAGCGAAAGATGAGGGTGGCAGGAA<br>GAGAAGGGCGCTTTCTGTCTGCCGGGGTCGCAGCGCGAGAGGGCAGTGCCATGTTCCTCTCCATCCTA<br>GTGGCGCTGTGCCTGTGGCTGCACCTGGCGCTGGGCGTGCGCGGCGCGCCCTGCGAGGCGGTGCGCAT<br>CCCTATGTGCCGGCACATGCCCTGGAACATCACGCGGATGCCCAACCACCTGCACCACAGCACGCAGG<br>AGAACGCCATCCTGGCCATCGAGCAGTACGAGGAGCTGGTGGACGTGAACTGCAGCGCCGTGCTGCGC<br>TTCTTCCTCTGTGCCATGTACGCGCCCATTTGCACCCTGGAGTTCCTGCACGACCCTATCAAGCCGTGCA<br>AGTCGGTGTGCCAACGCGCGCGCGACGACTGCGAGCCCCTCATGAAGATGTACAACCACAGCTGGCCC<br>GAAAGCCTGGCCTGCGACGAGCTGCCTGTCTATGACGGTTCTGCTGGAAGCGCAGCCGGCTCCGGGGA<br>GTTCTCTAAGGGTGAGGAGCTATTCACAGGCGTGGTACCCATCTTGGTAGAGCTGGACGGGGACGTCA<br>ACGGTCACAAATTCTCAGTGCGAGGAGAGGGCGAGGGTGACGCTACTAACGGCAAGTTAACACTGAA<br>GTTTATCTGCACGACAGGCAAGTTGCCCGTTCCTTGGCCAACGCTTGTCACTACCCTGACTTACGGGGT<br>GCAGTGTTTCAGTCGCTACCCGGACCATATGAAAAGACACGACTTTTTTAAGTCCGCGATGCCCGAAGG<br>CTACGTTCAAGAGCGGACAATATCCTTTAAAGACGACGGTACCTACAAGACACGTGCCGAAGTCAAGT<br>TCGAGGGAGATACTTTAGTCAACAGGATCGAACTCAAAGGAATTGACTTCAAAGAGGACGGGAATATC<br>CTCGGACATAAACTGGAGTACAACTTTAACTCGCACAACGTTTATATCACTGCCGATAAACAGAAGAA<br>CGGAATAAAAGCCAACTTTAAAATTAGACACAACGTGGAAGACGGAAGCGTGCAGCTCGCAGACCATT<br>ATCAACAGAACACCCCCATTGGAGACGGACCAGTGCTGCTGCCAGATAATCATTATCTCTCTACCCAGA<br>GTGTGCTGAGCAAGGATCCAAACGAAAAGAGGGATCATATGGTGCTGCTTGAGTTCGTGACCGCTGCA<br>GGGATTACCCACGGCATGGACGAACTTTATAAGGGCTCACACCACCACCATCACCATTGAGCTAACTA<br>GTTTCCAAAGCGGAGACTTCCGACTTCCTTACAGGATGAGGCTGGGCATTGCCTGGGACAGCCTATGTA<br>AGGCCATGTGCCCCTTGCCCTAACAACTCGTCGACCCCGGGATGCCGTGAGCACAAAAGCGACCATAG<br>TCGGGATTTTACCCGCTTATCTGCGGAAGGTCATTGGGAACGCATCTTCCACAAATGGGCTGAGCGCAA<br>AAATGCCGAACGTATCTGGCTGACCTATGCCATTCCGTGGGAACGACACTACTTTTCAATTGGTGCACT<br>CTCAGTACAATCTGCTCTGATGCCGCATAGTTAAGCCAGCCCCGACACCCGCCAACACCCGCTGACGCG<br>CCCTGACGGGCTTGTCTGCTCCCGGCATCCGCTTACAGACAAGCTGTGACCGTCTCCGGAAAATCAAAG<br>GATCTTCTTGAGATCCTTTTTTTGAGCTGCATGTGTCAGAGGTTTTCACCGTCATCACCGAAACGCGCGA<br>GACGAAAGGGCCTCGTGATACGCCTATTTTTATAGGTTAATGTCATGATAATAATGGTTTCTTAGACGT<br>CAGGTGGCACTTTTCGGGGAAATGTGCGCGGAACCCCTATTTGTTTATTTTTCTAAATACATTCAAATAT<br>GTATCCGCTCATGAGACAATAACCCTGATAAATGCTTCAATAATATTGAAAAAGGAAGAGTATGAGTA<br>TTCAACATTTCCGTGTCGCCCTTATTCCCTTTTTTGCGGCATTTTGCCTTCCTGTTTTTGCTCACCCAGAA<br>ACGCTGGTGAAAGTAAAAGATGCTGAAGATCAGTTGGGTGCACGAGTGGGTTACATCGAACTGGATCT<br>CAACAGCGGTAAGATCCTTGAGAGTTTTCGCCCCGAAGAACGTTTTCCAATGATGAGCACTTTTAAAGT<br>TCTGCTATGTGGCGCGGTATTATCCCGTATTGACGCCGGGCAAGAGCAACTCGGTCGCCGCATACACTA<br>TTCTCAGAATGACTTGGTTGAGTACTCACCAGTCACAGAAAAGCATCTTACGGATGGCATGACAGTAA<br>GAGAATTATGCAGTGCTGCCATAACCATGAGTGATAACACTGCGGCCAACTTACTTCTGACAACGATC<br>GGAGGACCGAAGGAGCTAACCGCTTTTTTGCACAACATGGGGGATCATGTAACTCGCCTTGATCGTTG<br>GGAACCGGAGCTGAATGAAGCCATACCAAACGACGAGCGTGACACCACGATGCCTGTAGCAATGGCA<br>ACAACGTTGCGCAAACTATTAACTGGCGAACTACTTACTCTAGCTTCCCGGCAACAATTAATAGACTGG<br>ATGGAGGCGGATAAAGTTGCAGGACCACTTCTGCGCTCGGCCCTTCCGGCTGGCTGGTTTATTGCTGAT<br>AAATCTGGAGCCGGTGAGCGTGGGAGTCGCGGTATCATTGCAGCACTGGGGCCAGATGGTAAGCCCTC<br>CCGTATCGTAGTTATCTACACGACGGGGAGTCAGGCAACTATGGATGAACGAAATAGACAGATCGCTG<br>AGATAGGTGCCTCACTGATTAAGCATTGGTAACTGTCAGACCAAGTTTACTCATATATACTTTAGATTG<br>ATTTAAAACTTCATTTTTAATTTAAAAGGATCTAGGTGAAGATCCTTTTTGATAATCTCATGACCAAAAT<br>CCCTTAACGTGAGTTTTCGTTCCACTGAGCGTCAGACCCCGTAGAAAAGATCAAAGGATCTTCTTGAGA<br>TCCTTTTTTTCTGCGCGTAATCTGCTGCTTGCAAACAAAAAAACCACCGCTACCAGCGGTGGTTTGTTTG<br>CCGGATCAAGAGCTACCAACTCTTTTTCCGAAGGTAACTGGCTTCAGCAGAGCGCAGATACCAAATAC<br>TGTTCTTCTAGTGTAGCCGTAGTTAGGCCACCACTTCAAGAACTCTGTAGCACCGCCTACATACCTCGC<br>TCTGCTAATCCTGTTACCAGTGGCTGCTGCCAGTGGCGATAAGTCGTGTCTTACCGGGTTGGACTCAAG<br>ACGATAGTTACCGGATAAGGCGCAGCGGTCGGGCTGAACGGGGGGTTCGTGCACACAGCCCAGCTTGG<br>AGCGAACGACCTACACCGAACTGAGATACCTACAGCGTGAGCTATGAGAAAGCGCCACGCTTCCCGAA<br>GGGAGAAAGGCGGACAGGTATCCGGTAAGCGGCAGGGTCGGAACAGGAGAGCGCACGAGGGAGCTTC<br>CAGGGGGAAACGCCTGGTATCTTTATAGTCCTGTCGGGTTTCGCCACCTCTGACTTGAGCGTCGATTTTT<br>GTGATGCTCGTCAGGGGGGCGGAGCCTATGGAAAAACGCCAGCAACGCGGCCTTTTTACGGTTCCTGG<br>CCTTTTGCTGGCCTTTTGCTCACATGTTCTTTCCTGCGTTATCCCCTGATTCTGTGGATAACCGTATTACC<br>GCCTTTGAGTGAGCTGATACCGCTCGCCGCAGCCGAACGACCGAGCGCAGCGAGTCAGTGAGCGAGGA<br>AGCGGAAGAGCAAATAATAAAAAAGCCGGATTAATAATCTGGCTTTTTATATTCTCTGCCCAATACGC<br>AAACCGCCTCTCCCCGCGCGTTGGCCGATTCATTAATGCAGCTGGCACGACAGGCAAAGACTTCTACG<br>GTTGGGCAATCGAACGCCACTATTCAGCGCATGACCTGTTTAAGGCGCATAGCTTTGATCATGCCGGTA<br>GAGCACGTTGGGAAGGACATCGTTGGGCACATCGTGAACATGCCAACATGTTTTGGGCGCATGAAGGC<br>ATTTGGCGTATCCACTGACT |

*Boxes indicate the translational start and stop codons of the proteins (oSFRP4 and SFRP4-sfGFP). Green, pink, and yellow highlights correspond to *SFRP4* (C-terminally truncated), flexible linker, and *sfGFP* sequences, respectively. Underlines indicate the primer annealing sites. Gray highlight indicates the pEX-A2J2 vector sequence.

**table S4**. List of PCR primers

| No | Primer name | Primer sequence (5' to 3') |
|---|---|---|
| 1 | Fw1-C | GAAATAATTTTGTTTAACTTTAAGAAGGAGATATACCAATGGGACATCATCATCATCATCACGATTACG |
| 2 | Fw1-PG1 | CATTTTACATTCTACAACTACATAACTAACTAGTAACAATGTCTGGCCTCCAAGGACCGTCGGTGGGC |
| 3 | Fw1-PG2 | CATTTTACATTCTACAACTACATAACTAACTAGTAACAAGGTCTGGCCTCCAAGGACCGTCGGTGGGC |
| 4 | Fw1-SFG1 | CATTTTACATTCTACAACTACATAACTAACTAGTAACAATGGCCGTGGCTGGCTGCGCTCCGAGCTGC |
| 5 | Fw1-SFG2 | CATTTTACATTCTACAACTACATAACTAACTAGTAACAAGGGCCGTGGCTGGCTGCGCTCCGAGCTGC |
| 6 | DsRed-Fw1 | CGCCTAGGCGTCACCGGCCATCACTAGTCGGACGGGGCGGCGAGACGCTACGCGTCGCCACCATGGCCTCCTCCGAGGACGTCATC |
| 7 | DsRed-Rv1 | TTATGATCTAGAGTCGGCGCGCCTCTACAGGAACAGGTGGTGGCGGCCCTCGGC |
| 8 | DsRed-Fw2 | CTGTAGAGGCGCGCCGACTCTAGATCATAATCAGCCATACCACATTTGTAGAGG |
| 9 | DsRed-Rv2 | CGACTAGTGATGGCCGGTGACGCCTAGGCGTAGAGTAGCAGAACGACGTCTTGATGGGATCTGACGGTTCACTAAACCAGCTCTGC |
| 10 | DsRed-Fw3 | CGCTACAGGGCGCGTTAGTTATTAATAGTAATCAATTACGGGGTC |
| 11 | DsRed-Rv3 | CGAAAAGTGCCACCTGACGCCTTAAGATACATTGATGAGTTTGGAC |
| 12 | AmCyan-Fw | GTATCTTAAGGCGTCAGGTGGCACTTTTCGGGGAAATGTGCGCG |
| 13 | AmCyan-Rv | TACTATTAATAACTAACGCGCCCTGTAGCGGCGCATTAAGCGCGG |
| 14 | oORF-244aa-Fw | TCGAGCTCAAGCTTCGAATTCTGCAGTCGACGCCCCCATGGCCGCTGCAGCCACCGGTCGCCACCATGGCCCTGTCCAACAAGTTCA |
| 15 | oORF-14aa-Fw | CGGTCGCCACCATGGCCCTGTCCAATAAGTTCATCGGCGACGACATGAAGATGAC |
| 16 | oORF-14aa-Rv | GTCATCTTCATGTCGTCGCCGATGAACTTATTGGACAGGGCCATGGTGGCGACCG |
| 17 | oORF-31aa-Fw | GGCGACGACATGAAGATGACCTACCACATGGACGGCTGCGTTAACGGCCACTACTTCACCG |
| 18 | oORF-31aa-Rv | CGGTGAAGTAGTGGCCGTTAACGCAGCCGTCCATGTGGTAGGTCATCTTCATGTCGTCGCC |
| 19 | oORF-60aa-Fw | CCTTCAAGGTGACCATGGCTAACGGCGGCCCCCTGGCCTTCTCCTTC |
| 20 | oORF-60aa-Rv | GAAGGAGAAGGCCAGGGGGCCGCCGTTAGCCATGGTCACCTTGAAGG |
| 21 | oORF-182aa-Fw | GACCGCCTTCCTGATGCTGCAGGGCGGCGGTAACTACAGATGCCAGTTCCACACCTC |
| 22 | oORF-182aa-Rv | GAGGTGTGGAACTGGCATCTGTAGTTACCGCCGCCCTGCAGCATCAGGAAGGCGGTC |

**Data. S1. Investigation of Ribo-Seq-based translatomic datasets in humans.**

**Data. S2. Data mining of NCBI ClinVar for the identification of oORFs (USURPs) arising from pathogenic frameshift polymorphisms in humans.**
